# Multi-scale spatial mapping of cell populations across anatomical sites in healthy human skin and basal cell carcinoma

**DOI:** 10.1101/2023.08.08.551504

**Authors:** Clarisse Ganier, Pavel Mazin, Gabriel Herrera-Oropeza, Xinyi Du-Harpur, Matthew Blakeley, Jeyrroy Gabriel, Alexander Predeus, Batuhan Cakir, Martin Prete, Nasrat Harun, Jean-Francois Darrigrand, Alexander Haiser, Saranya Wyles, Tanya Shaw, Sarah A. Teichmann, Muzlifah Haniffa, Fiona M. Watt, Magnus D. Lynch

**Affiliations:** Centre for Gene Therapy and Regenerative Medicine, King’s College London, Guy’s Hospital, Great Maze Pond, London, UK; Wellcome Sanger Institute, Wellcome Genome Campus, Hinxton, Cambridge CB10 1SA, UK; Centre for Developmental Neurobiology, Institute of Psychiatry, Psychology and Neuroscience, King’s College London, London SE1 1UL, UK; The Francis Crick Institute, 1 Midland Road, London, UK; Department of Dermatology, Mayo Clinic, 200 First St. SW, Rochester, MN 55905; Centre for Inflammation Biology & Cancer Immunology, King’s College London, London SE1 1UL, UK; Biosciences Institute, Newcastle University, Newcastle upon Tyne NE2 4HH, UK; Theory of Condensed Matter Group, Cavendish Laboratory, University of Cambridge, JJ Thomson Ave, Cambridge CB3 0HE, UK; Department of Dermatology and NIHR Newcastle Biomedical Research Centre, Newcastle Hospitals NHS Foundation Trust, Newcastle upon Tyne NE1 4LP, UK; Directors’ Unit, EMBL, Meyerhofstr. 1, 69117 Heidelberg, Germany; St John’s Institute of Dermatology, King’s College London, London, UK

**Keywords:** Human Cell Atlas, Facial Skin, Body Skin, Basal Cell Carcinoma, Spatial Transcriptomics, *In situ* sequencing, Single Cell Sequencing, Cancer Associated Fibroblasts, Pericytes

## Abstract

Our understanding of how human skin cells differ according to anatomical site and tumour formation is limited. To address this we have created a multi-scale spatial atlas of healthy skin and basal cell carcinoma (BCC), incorporating *in vivo* optical coherence tomography, single cell RNA sequencing, spatial global transcriptional profiling and *in situ* sequencing. Computational spatial deconvolution and projection revealed the localisation of distinct cell populations to specific tissue contexts. Although cell populations were conserved between healthy anatomical sites and in BCC, mesenchymal cell populations including fibroblasts and pericytes retained signatures of developmental origin. Spatial profiling and *in silico* lineage tracing support a hair follicle origin for BCC and demonstrate that cancer-associated fibroblasts are an expansion of a *POSTN*+ subpopulation associated with hair follicles in healthy skin. *RGS5+* pericytes are also expanded in BCC suggesting a role in vascular remodelling. We propose that the identity of mesenchymal cell populations is regulated by signals emanating from adjacent structures and that these signals are repurposed to promote the expansion of skin cancer stroma. The resource we have created is publicly available in an interactive format for the research community.

**Significance statement:** Single cells RNA sequencing has revolutionised cell biology, enabling high resolution analysis of cell types and states within human tissues. Here, we report a comprehensive spatial atlas of adult human skin across different anatomical sites and basal cell carcinoma (BCC) - the most common form of skin cancer - encompassing *in vivo* optical coherence tomography, single cell RNA sequencing, global spatial transcriptomic profiling and in situ sequencing. In combination these modalities have allowed us to assemble a comprehensive nuclear-resolution atlas of cellular identity in health and disease.

## Introduction

Human skin comprises two main layers, the epidermis and dermis; is home to cells of the innate and adaptive immune systems; and is extensively vascularised and innervated. Skin in different body sites performs specialised functions that are reflected in differences in the nature and density of adnexal structures – hair follicles (HFs), sebaceous glands and sweat glands. As a result of its exposure to UV light and other carcinogens, the skin is susceptible to developing cancer and basal cell carcinoma (BCC) is the most common type of cancer affecting humans (1).

In order to gain a fuller understanding of skin cell organisation and function in health and disease, a number of research teams, including our own, have used single cell RNA sequencing (scRNAseq) to create a comprehensive catalogue of the cell types present in healthy adult skin, developing skin and common inflammatory skin diseases (2). However, we currently lack a full understanding of whether the cell types in skin from different anatomical sites differ and we do not have high resolution spatial maps of where specific cell types are located.

The issue of cellular identity is particularly interesting because during development the dermis of the trunk and limbs is derived from somatic mesoderm whereas the dermis of the anterior head and neck is derived from cranial neural crest (3). The question of spatial cellular location is also important since lineage tracing studies in mice have shown that functionally distinct subpopulations of dermal cells have different spatial organisation (4–6).

A further important question is how the cell populations present in healthy skin relate to those in skin cancer (7). BCC is a locally invasive skin cancer composed of islands of tumour cells embedded in a dense fibroblastic stroma along with vascular elements and preferentially affects the face (8). The cell of origin of BCC has long been a subject of debate, with some studies pointing to a HF origin (9, 10). Growth of a BCC requires both expansion of stromal elements and the growth of new blood vessels (11). Altered proliferation of epithelial elements is driven primarily by cell-intrinsic genetic alterations (12), including *PTCH1, TP53, NOTCH1, NOTCH2* and *FAT1* mutations in the case of BCC (13). However the origins and drivers of the associated stromal changes are not fully understood. Fibroblasts within tumour stroma are referred to as ‘cancer associated Fibroblasts’ (CAFs) (14) and are considered as a potential target for therapeutic strategies (15) but it is not clear whether they have a distinct cellular identity compared to healthy dermal fibroblasts.

The goals of the Human Cell Atlas (16) are to define the states of all cell populations within tissues in both health and disease and to understand their spatial localisation with respect to one another. Here, we report a comprehensive spatial atlas of cell populations in human skin across multiple scales incorporating Visium spatial transcriptomics (ST) at 55μm resolution, *in situ* sequencing (ISS) of transcripts at subcellular resolution, <10μm-resolution optical coherence tomography (OCT) imaging of live human skin and scRNAseq of skin from multiple body sites and BCC. Our analysis has allowed us to create maps of cellular populations at single cell resolution for healthy skin at different anatomical sites and for BCC. We believe this will be a valuable resource for the community.

## Results

### Cartography of a multi-scale atlas of healthy human skin and basal cell carcinoma

To understand the landscape of skin cell populations, we studied healthy skin across multiple anatomical sites and BCC (Fig. 1A). The human skin exhibits substantial morphological variation across anatomical sites, such as differences in epidermal thickness, HF, sebaceous gland, and eccrine coil density (SI Appendix, Fig. 1A, B).

**Figure 1.**
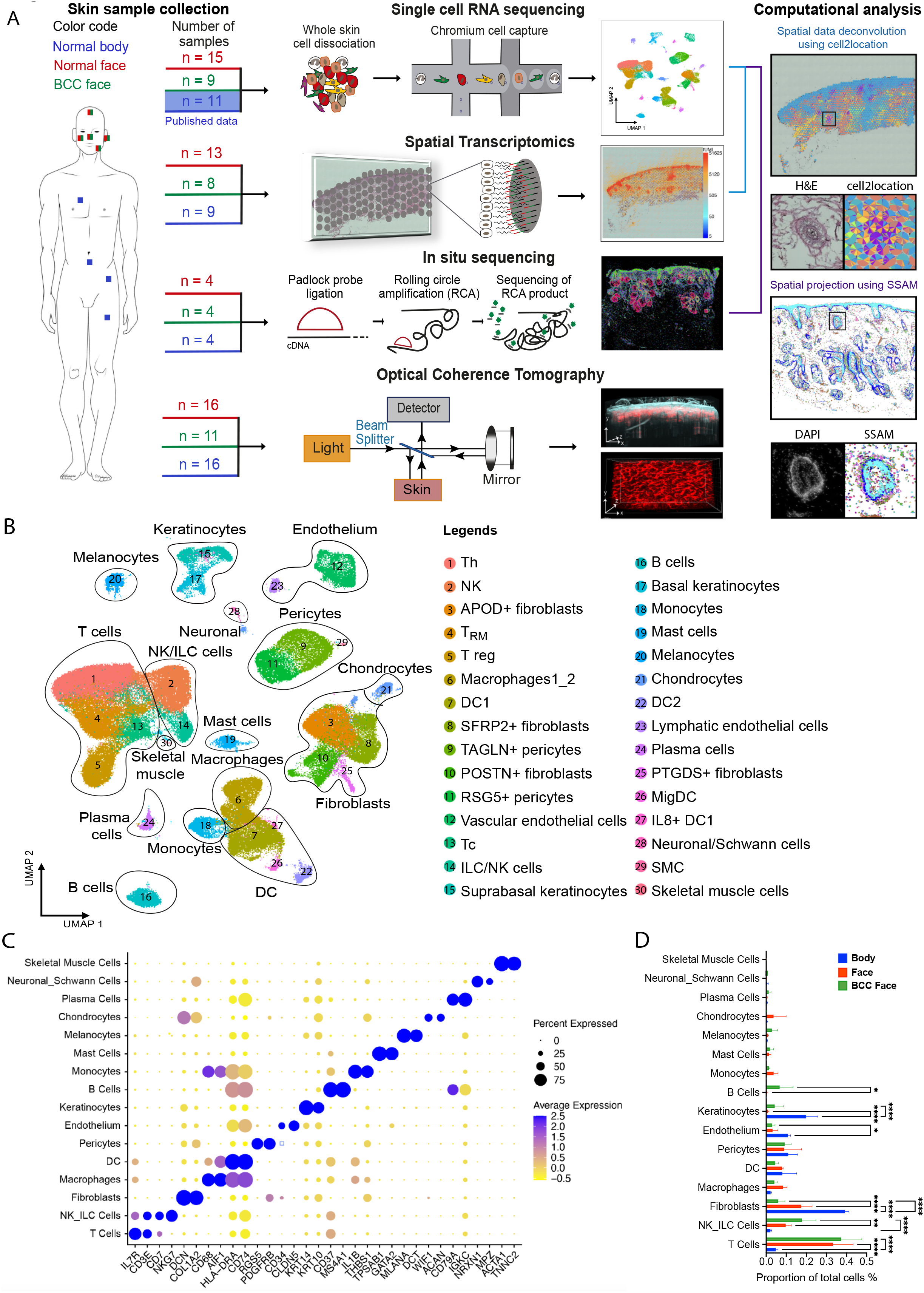
Cartography of a multi-scale human skin atlas across anatomical sites and cancer. A) ScRNAseq was performed for skin samples derived from multiple facial sites and compared with previously published scRNAseq of body sites. We also profiled skin samples from these conditions with global spatial transcriptomics (ST) which permitted us to examine the global transcriptional profile at 55μm resolution; *in situ* sequencing which mapped skin at subcellular resolution with a custom panel of 165 genes (SI Appendix, Table 2, 3 and 4) designed to represent all the cellular populations present in human skin; and optical coherence tomography imaging which permitted us to assess the morphology of the skin at 10μm *in vivo*. The number of samples for each modality is indicated. The location and identity of cells were inferred probabilistically in the global ST data and in the ISS data according to our annotated scRNA-seq dataset through computational analyses. B) Uniform Manifold Approximation and Projection (UMAP) and clustering of 155,401 cells from 33 donors (11 healthy body sites, 14 healthy face sites, and 8 BCC patients) representing 30 skin cell populations and 16 cell types. Each cell is represented by a single point with the two dimensional location of this point on the UMAP plot corresponding to the global transcriptional state of the cell. Cells with similar transcriptional profile form clusters. C) Dotplot showing well known marker genes specific to each skin cell type. For each cell type the percentage of cells expressing the marker (diameter) and the average log2 normalized expression (color) is shown. D) Abundance of skin cell types in healthy face and body areas and in BCC from the face. The relative proportion of sequenced cells assigned to each cellular population is illustrated for body, face and BCC face. Significant differences (One way ANOVA-test) are indicated (* <0.05, **<0.01, ****<0.0001).

Previous scRNAseq studies of human skin have largely focused on the skin of the trunk and limbs(2, 17–20). However, BCC is more common in facial skin (7, 8). We therefore augmented existing publicly available datasets through scRNAseq of full thickness skin biopsies from 15 healthy donors and 8 BCCs across 5 facial skin sites (Ear, Nose, Cheek, Forehead, Temple). Additionally, we used two complementary approaches to localize cells within tissue sections: global ST (Visium) and targeted ISS(21, 22). We also quantified skin vasculature structure with angiographic OCT across anatomical sites. OCT images of *in vivo* human skin obtained at 10μm resolution through the principle of low-coherence interferometry(23) yielded high resolution 3D images of cutaneous vasculature (Fig. 1A, SI Appendix, Fig. 1C). OCT showed differences in the dermal density of vascular networks between face and body skin (SI Appendix, Fig. 1D).

### scRNAseq in healthy skin and BCC samples

We sequenced 233,379 cells, of which 134,839 cells passed quality control (QC). We integrated our sequenced cells with two publicly available datasets from 11 healthy body sites (Inguinal, Arm) to give a combined dataset of 155,401 cells (Fig. 1B; SI Appendix, Fig. 2A, B, C). Unsupervised clustering revealed a total of 30 clusters corresponding to 16 cell types (Fig. 1B, SI Appendix, Fig. 2D). Cell types and subcelltypes were annotated with previously described markers (2, 17, 19, 24–26) (Fig. 1C, SI Appendix Fig 2E and Table 1).

The mesenchymal cells that constitute the dermis of the face are derived from neural crest(3), whereas those of the body are derived from somatic mesoderm. Previous studies have revealed developmental gene expression differences in fibroblasts at different anatomical sites in both human and mouse (27, 28). We confirmed that adult skin cells retained signatures of their embryological origins and we found differences in expression of Hox genes and mesenchymal neural crest markers between face, including BCC, and body skin in scRNAseq and spatial transcriptomic datasets (SI Appendix, Fig. 3A, B), particularly for fibroblasts, pericytes, SMC, vascular endothelial cells and chondrocytes (SI Appendix, Fig. 3C, D). These findings demonstrate that signatures associated with developmental origin can be detected in a cell type-specific manner in cells derived from adult face and body skin.

Although, we found skin cell populations to be conserved (Fig. 1B); we identified a mesenchymal population that was present only in a full thickness ear sample and on the basis of marker expression (including high expression of ACAN and COL2A1), we identified these as extra-articular chondrocytes (Fig. 1B, C, D; SI Appendix, Fig. 2E, F). We additionally identified a skeletal muscle cell population that was present only in a forehead sample (Fig. 1B, C, D; SI Appendix, Fig. 2E, F). The face contains 20 distinct flat skeletal muscles that attach to the skull, including the forehead(29). Those two cell types are not strictly skin cell types and were not represented across all the samples; therefore we did not study them further.

Differences in the relative abundance of cell types were observed. T cells were over-represented in facial skin and BCC compared to body skin. NK/ILC and B cells were largely over-represented in BCC compared to healthy skin (face and body sites). NK/ILC were also more abundant in face compared to body sites. Fibroblasts were more abundant in body skin compared to facial skin. The decrease is even more important in BCC. A difference in the abundance of keratinocytes was also observed between face and body skin, which likely reflects both differences in epidermal thickness and in the efficiency of keratinocyte isolation (Fig. 1D, SI Appendix, Fig. 2D).

### Mapping of skin cell populations using global spatial transcriptomics and targeted *in situ* **sequencing**

Having catalogued cell populations across anatomical sites and in BCC, we localized them within the tissue by combining scRNAseq data with global ST and targeted ISS (21, 22).

We prepared ST libraries for 9 body, 13 face and 8 BCC sections (Fig. 1A, SI Appendix, Fig. 4A). The presence of high-quality RNA throughout sections was verified through housekeeping gene expression via multiplex RNA FISH (SI Appendix, Fig. 4B). The percentage of mitochondrial genes and Unique Molecular Identifier (UMI) counts in the Visium sections confirmed that the libraries were of high quality (SI Appendix, Fig. 4C, D), and the few low-quality spots were removed. To deconvolve cell types present within each Visium 55μm grid spot, we used a published algorithm, cell2location(22) - a Bayesian model that can resolve fine-grained cell types in ST data to create comprehensive cellular maps of diverse tissues, permitting the inference of cell types present on the basis of an annotated scRNAseq dataset (Fig. 1A, B).

An advantage of ST is the ability to analyse the entire transcriptome in whole tissue sections. However, resolution is limited to 55μm grid spots and highly expressed genes can dominate sequencing libraries in these spots. To compensate these, we performed ISS. This approach utilises rolling-circle amplification in combination with barcoded padlock-probe circularization (30).

Based on published scRNAseq data (2, 17, 19, 20, 24–26) and our sequencing of facial skin, we defined a panel of 165 marker genes representing all skin cell populations (SI Appendix, Table 2, 3, 4). The presence of high-quality RNA was verified through housekeeping gene expression via ISS workflow (SI Appendix, Fig. 4E). We then performed ISS of 4 body, 4 face and 4 BCC sections (Fig. 1A, SI Appendix, Fig. 4A, F). According to Qian *et al*.,(21), 50 optimally-chosen markers were sufficient for accurate classification of 28 cell populations. Since the number of cell populations present in our human skin dataset is comparable to their analysis, our panel of 165 marker genes should be adequate for cell typing. To do so, we used the SSAM pipeline (31), which allows inference of cell types from mRNA signals in a cell segmentation-free manner by correlating gene expression with an annotated scRNAseq dataset. To demonstrate the power of those two computational spatial methods, we predicted and projected the main skin cell types such as the keratinocyte clusters, fibroblast clusters, endothelial cells, pericytes and SMC clusters onto the ST sections in regard to the localisation of ISS reads representing those main skin cell types (SI Appendix, Fig. 5A, B, C).

We next applied the same methodologies to the 30 clusters found in our dataset (Fig. 1B; SI Appendix, Fig. 2D, E and Table 1) to create a spatial skin atlas. To facilitate the use of the atlas, we generated a public web interface (https://spatial-skin-atlas.cellgeni.sanger.ac.uk/), where researchers can explore spatial gene expression patterns and skin cell population predictions. This resource contains an explorer called CELLxGENE for scRNA-seq data and ST data and another explorer called Vitessce (32) for ISS data.

### The architecture of cutaneous blood vessels reflects vessel size and anatomical site and is altered in BCC

In healthy skin, the cutaneous vasculature is organised as a superficial (in the papillary dermis) and a deep (in the reticular dermis) vascular plexuses connected by perforating vessels. In BCC, tumour epithelial cells are embedded in a dense vascular and fibroblastic stroma. Growth of the tumour to a macroscopic size requires both expansion of stromal elements and the growth of new blood vessels (11). Since the cutaneous vasculature is a dynamic 3D network, histological sections do not accurately capture its structure in vivo. Therefore we used angiographic (speckle contrast) OCT (33) to generate 3D images of vasculature density across facial and body sites from the same 16 healthy individuals (Fig. 2A, B) and of matched healthy and BCC areas from the same 11 patients (Fig. 2C, D).

**Figure 2.**
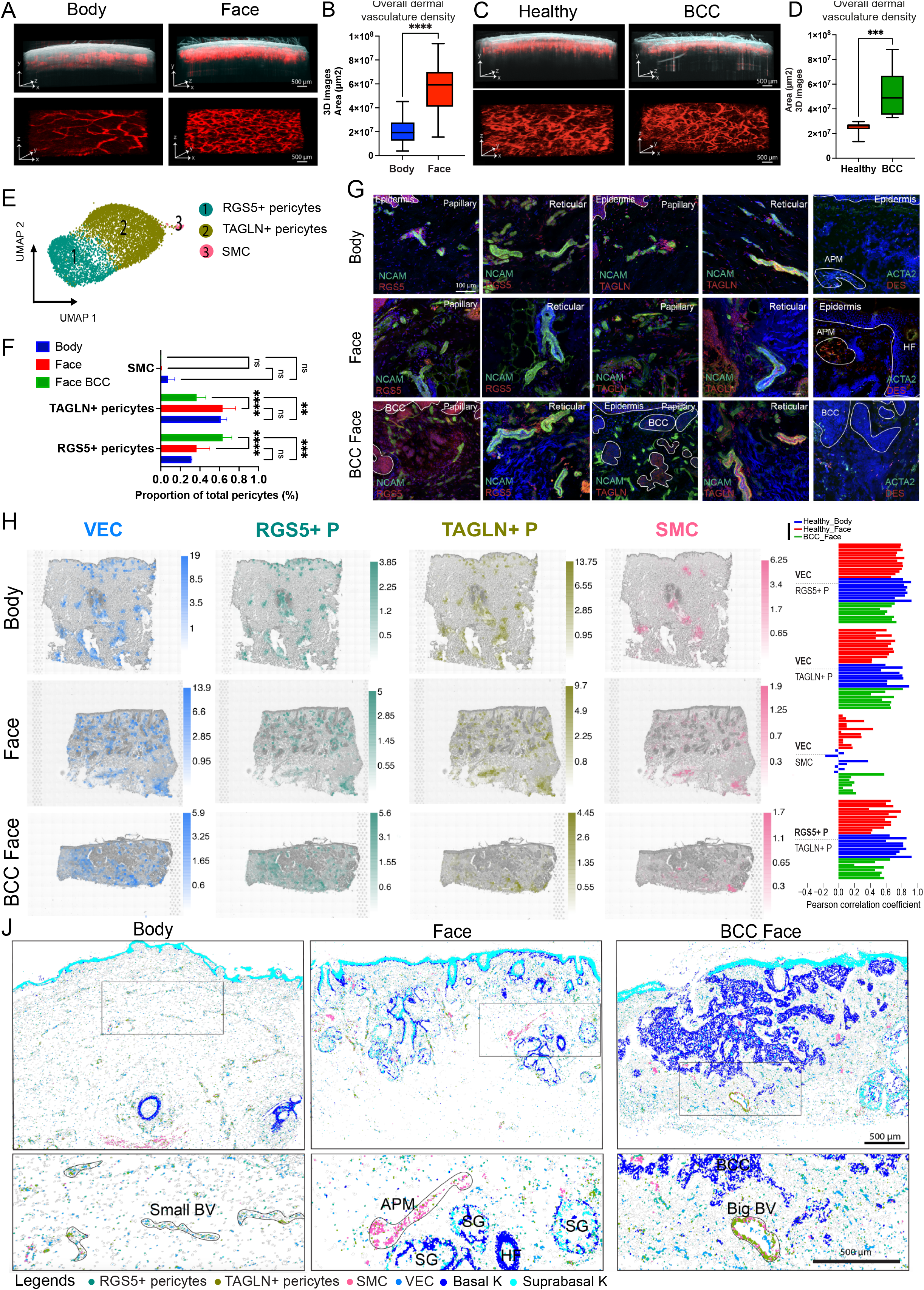
Healthy morphology of cutaneous vasculature and localisation of vascular cell populations and their alterations in the stroma of BCC. A) 3D morphology of vascular networks is compared for representative OCT scans of facial and body skin from the same individuals. OCT images (500 frames) illustrate reflectivity (grayscale), overlaid with the blood flow (red) (upper panel). The skin vasculature network is also shown as a 3D computational reconstruction (lower panel). Image size 6 mm × 6 mm; scale bar: 500um. B) Quantification of skin vascular density across facial and body sites from the same individuals. Vascular density was quantified for a total of 6 sites, 3 facial sites and 3 body sites, across 16 individuals. Statistically significant differences (One way ANOVA-test) are indicated are indicated (****<0.0001). C) Comparison of vascular architecture in healthy skin area and BCC sites from the same patients. Image size 6 mm × 6 mm; scale bar: 500um. D) Quantification of skin vascular density between healthy and BCC sites from the same patients. Vascular density was quantified for a total of 3 facial sites across 11 individuals. Statistically significant differences (One way ANOVA-test) are indicated (***<0.001)). E) UMAP plot of pericytes (*RGS5*+ pericytes, *TAGLN*+ pericytes), smooth muscle cells (SMC) and vascular endothelial cells (VEC) subpopulations present in healthy skin (face and body) and BCC. F) Comparison of the abundance of different pericyte subpopulations (as a proportion of total pericytes) in healthy facial and body skin and BCC. Statistically significant differences (One way ANOVA-test) are indicated are indicated (ns, not significant, **<0.01, ***<0.001, ****<0.0001). G) Immunostaining of NCAM, RGS5 and TAGLN in papillary and reticular dermis in healthy body and face skin and in BCC. RNAscope imaging of *ACTA2* and *DES* in healthy body and face skin and in BCC (DAPI, blue). H) VEC and pericyte cell populations annotated in scRNAseq data were computationally predicted on global ST sections using cell2location. Predicted cell abundances shown by color gradients per spot into tissue architecture (H&E) images (representative samples are shown). I) Cell cluster colocalization analysis for VEC with *RGS5+, TAGLN+* pericytes and SMC. Barplot shows pearson correlation coefficients of cell2location predictions per spot normalized cell abundances across all spots of Visium samples (each individual bar represents a Visium sample). J) Computational spatial mapping of VEC and pericyte subpopulations in *ISS* sections using SSAM (representative samples are shown). Localization of basal and suprabasal keratinocytes are shown in order to indicate skin structures.

In healthy skin, vascular density was significantly higher in facial skin compared to body sites in both papillary and reticular dermis (Fig. 2A, B; SI Appendix, Fig. 1C, D). However, we found no significant differences in segment length, number of segments, tube thickness or number of branching points in face skin samples (SI Appendix, Fig. 1E).

In BCC patients, OCT images showed a disorganised vascular plexus with tortuous vessels (Fig. 2C; SI Appendix, Fig. 6A), as previously published(34, 35). Quantification revealed a significant increase in blood vessel density in the tumour compared to adjacent skin from the same patient (Fig. 2D).

We next analysed the vascular cell populations present in our integrated scRNAseq dataset. In contrast to previous studies (20), we were able to identify only one population of VEC and one of LEC (Fig. 1B). Regarding peri-vascular cell populations, we found three populations of pericytes (Fig. 2E). In keeping with previous studies (20), there were two major pericyte populations, which we denoted RGS5+ and TAGLN+ based on high expression of those genes (SI Appendix, Fig. 2E). We additionally identified a third population expressing high levels of DES and ACTA2 which is a gene signature of SMC (SI Appendix, Fig. 2E). Gene ontology (GO) analysis revealed SMC to have signatures of muscle and contractile function, whereas RGS5+ pericytes exhibited signatures of blood vessel development and morphogenesis. TAGLN+ pericytes showed to have signature in both blood vessel development and contractile function (SI Appendix, Fig. 6B).

Despite the increased density of blood vessels in facial skin, we did not find a significant difference in the abundance of VEC, LEC (Fig. 1D) or pericyte subpopulations (Fig. 2F) between healthy face and body skin. However, there was a selective expansion of the RGS5+ pericytes and a reduction in TAGLN+ pericytes in BCC compared to healthy skin (face and body) (Fig. 2F).

Immunostaining of RGS5 in papillary and reticular dermis showed RGS5+ cells to be in close contact with NCAM+ cells, a pan vascular marker, in healthy skin and BCC (Fig. 2G). TAGLN+ cells colocalized and were in close contact with NCAM+ cells in papillary and reticular dermis and in small and large blood vessels in healthy face and body skin. The colocalization of TAGLN+ pericytes with small blood vessels was lost in BCC areas (Fig. 2G). In healthy skin, DES and ACTA2 RNA hybridization was observed in the arrector pili muscle (APM) cells in proximity to HFs. In BCC, SMC (ACTA2+ DES+ cells) were diffusely distributed within the stroma (Fig. 2G). RGS5, TAGLN and ACTA2 distributions were confirmed in the global ST data and in ISS (SI Appendix, Fig. 6C, D).

In healthy skin, ST deconvolution analysis revealed VEC to be localized in superficial and deep vascular plexuses (Fig. 2H). Both RGS5+ and TAGLN+ pericytes colocalized with VEC and with one another throughout the dermis (Fig. 2H). In BCC, the distribution of RGS5+ pericytes closely paralleled with VEC, whereas the distribution of TAGLN+ pericytes was more focal and less abundant (Fig. 2H). This is in contrast to healthy skin. In both healthy skin and BCC, SMC were adjacent to HFs and in the lower dermis, suggesting an association with the deep vascular plexus (Fig. 2H). Pearson correlation coefficients (PCC) of spot normalized cell abundances using the cell2location predictions in all the ST samples confirmed a strong colocalization of RGS5+ and TAGLN+ pericytes with VEC and with one another. Although fewer in number, TAGLN+ pericytes in BCC still colocalized with VEC (Fig. 2I).

We also applied CellPhoneDB(36), a receptor-ligand interaction analysis, this revealed a high number of predicted interactions (> 100) between RGS5+ and TAGLN+ pericytes and VEC (SI Appendix, Fig. 6E). Significant biological ligand–receptor interactions are predicted (SI Appendix, Fig. 6F) for RGS5+ pericytes with VEC and TAGLN+ pericytes with VEC as those predicted cell clusters are part of the same spatial microenvironment in ST data.

At spatial single cell level, RGS5+ and TAGLN+ pericytes were also localized to both small and large blood vessels. SMC were restricted to larger vessels and additionally colocalized with the APM (Fig. 2J; SI Appendix, Fig. 6C, D).

### Fibroblast subpopulations are spatially restricted within the dermis of healthy skin and BCC stroma

Previous studies in both the mouse (4, 5) and human (2, 6, 17, 19, 20, 24, 25) have revealed the presence of multiple fibroblast subpopulations in the dermis. The number of subpopulations found in human skin varies between studies and spatial information at the tissue level and level of anatomical sites is lacking.

Unbiased clustering of our integrated scRNAseq dataset identified 4 main populations of fibroblasts which we designated APOD+, SFRP2+, PTGDS+ and POSTN+ fibroblasts based on specific gene expression (Fig. 3A). The same number of fibroblast clusters was present in face, body and BCC skin. However, there were significant differences in their abundance and distribution.

**Figure 3.**
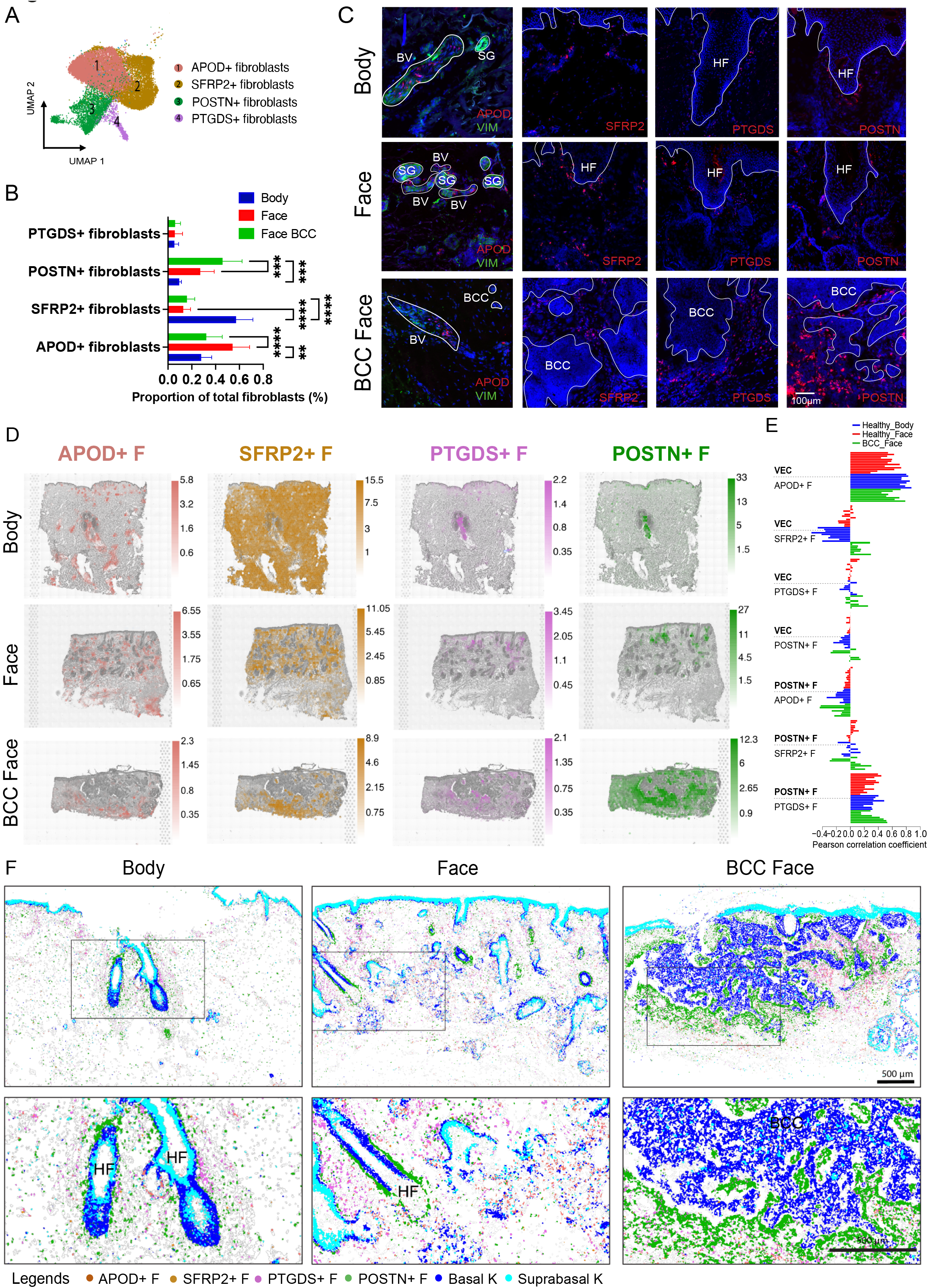
Localisation of fibroblast cell populations in healthy human skin and expansion of *POSTN*+ fibroblast subpopulations in the stroma of BCC. A) UMAP plot of four main fibroblast subpopulations in human healthy skin and BCC (*APOD*+ fibroblasts; *SFRP2*+ fibroblasts; *POSTN*+ fibroblasts; *PTGDS*+ fibroblasts). B) Comparison of the abundance of fibroblast subpopulations (proportion of total fibroblasts) between body and facial skin and BCC face. Statistically significant differences (One way ANOVA-test) are indicated (ns, not significant, **<0.01, ***<0.001), ****<0.0001). C) RNAscope of specific markers of each fibroblast subpopulation: *APOD, SFRP2, PTGDS, POSTN* (red), DAPI (blue) and immunostaining of VIM in healthy skin in BCC tissue sections. D) Fibroblast cell populations annotated in scRNAseq data were computationally predicted on global ST sections using cell2location. Predicted cell abundances shown by color gradients per spot into tissue architecture (H&E) images (representative samples are shown). E) Cell cluster colocalization analysis for VEC with *APOD+, SFRP2+, PTGDS+ and POSTN+* fibroblasts and *POSTN+* fibroblasts with *APOD+, SFRP2+,* and *PTGDS+* fibroblasts. Barplot shows pearson correlation coefficients of cell2location predictions per spot normalized cell abundances across all spots of Visium samples (each individual bar represents a Visium sample). F) Computational spatial mapping of fibroblast subpopulations in ISS sections using SSAM (representative samples are shown). Localization of basal and suprabasal keratinocytes are shown in order to indicate skin structures.

*APOD*+ fibroblasts were overrepresented in healthy facial skin in comparison with body skin and BCC (Fig. 3B). They highly co-expressed *APOD* and *APOE*, also expressed in immune cells (SI Appendix, Fig. 2E). GO analysis revealed potential functions in cell migration and mobilisation, characteristic of immune cells and implicated in humoral immune responses (SI Appendix, Fig. 7A). *APOD in situ* hybridization coupled with vimentin (VIM) immunostaining showed an association of *APOD*+ fibroblasts and blood vessels (VIM+) (Fig. 3C). The APOD distribution is shown in global ST data and ISS data (SI Appendix, Fig. 7B, C). ST deconvolution analysis and PCC confirmed *APOD*+ fibroblasts to be colocalized with VEC in facial and body areas and in BCC (Fig. 3D, E). Receptor-ligand analysis revealed a high number of predicted interactions (>100) between *APOD*+ fibroblasts and VEC (SI Appendix, Fig. 7D, E). The increased prevalence of *APOD*+ fibroblasts in facial skin (Fig. 3B) is in keeping with the increased vascular density compared to body skin (Fig. 2B, SI Appendix, Fig. 1C, D).

*SFRP2*+ fibroblasts were over-represented in body sites compared to facial sites and BCC (Fig. 3B). They co-expressed high levels of *SFRP2* and *WISP2* (SI Appendix, Fig. 2E). GO terms included extracellular matrix (ECM) structure and organisation, but also negative regulation of BMP signalling, which is known to control HF growth(37), which are characteristics of reticular fibroblasts (SI Appendix, Fig. 7A). Moreover, *WISP2* is a Wnt inhibitor(38). Epidermal Wnt signalling is important in the regulation of HF development and cycling(39). *SFRP2* RNAscope spots were present throughout the dermis (Fig. 3C). The distribution of *SFRP2* is confirmed in global ST data and ISS data (SI Appendix, Fig. 7B, C). ST deconvolution analysis confirmed *SFRP2*+ fibroblasts to be present throughout the dermis (Fig. 3D). The greater abundance of *SFRP2*+ fibroblasts in body skin (Fig. 3B) could be related to denser collagen and the lower quantity of epithelial structures in the body compared to face (SI Appendix, Fig. 1A, B).

The abundance of the *PTGDS*+ fibroblasts was similar between face and body skin and BCC (Fig. 3B). GO terms included ECM organisation and remodelling but also collagen fibril organisation and response to TGF-ß, which are characteristics of papillary fibroblasts (SI Appendix, Fig. 7A). *PTGDS* RNA spots were located in the papillary dermis in healthy face and body areas and around the tumour islets in BCC (Fig. 3C). The *PTGDS* distribution is shown in global ST data and ISS data (SI Appendix, Fig. 7B, C). *ST* and ISS predictions confirmed *PTGDS*+ fibroblasts to be located in papillary dermis and in proximity to follicular structures including eccrine coils. Furthermore, computational prediction in ISS, showed *PTGDS*+ fibroblasts to be frequently localized next to the upper HF areas such as bulge and infundibulum (Fig. 3F).

*POSTN*+ fibroblasts showed a trend of higher abundance in face skin compared to body skin and was significantly higher in BCC compared to healthy skin (Fig. 3B). Like *PTGDS*+ fibroblasts, GO terms showed function in ECM deposition and organisation and collagen fibril organisation but also characteristic of endoderm formation and development (SI Appendix, Fig. 7A), which is interesting given that cancer progression shares some gene expression profiles with developmental pathways (40). *POSTN* RNAscope spots was mainly around HFs in body and face skin and massively increased around the tumour islands in BCC (Fig. 3C). Expression of *POSTN* in ST and ISS confirmed these results (SI Appendix, Fig. 7B, C). ST deconvolution and PCC analysis showed *POSTN*+ fibroblasts to colocalized with *PTGDS*+ fibroblasts in both healthy body and face (Fig. 3D, E). Computational prediction in ISS revealed a localization next to lower shaft and bulb of the HF in healthy skin (Fig. 3F). Receptor-ligand analysis revealed a high number of predicted interactions (> 150) between *POSTN*+ and *PTGDS*+ fibroblasts (SI Appendix, Fig. 7D); multiple collagen subtypes and integrins are implicated. This suggests reciprocal cell-cell communication involving different fibroblast subpopulations within the HF niche (SI Appendix, Fig. 7D).

ST deconvolution and ISS projection confirmed that the expanded *POSTN*+ fibroblasts were localized in proximity to tumour islands (Fig. 3D, F). Other fibroblast subpopulations were also present in the stroma but distant from the tumour islands.

### Lack of distinct epithelial cell populations in BCC

The intimate association of the POSTN+ fibroblasts with epithelial tumour islands could potentially be mediated by an inductive signal arising from the epithelial cells. We hypothesised that since the POSTN+ fibroblasts were associated with HFs in healthy skin and expanded in BCC, the epithelial compartment of BCC would have characteristics of abnormal HFs. To test this, we explored the epithelial cell populations in BCC compared to healthy epidermis.

To be able to analyse keratinocytes in the interfollicular epidermis (IFE) and in follicular structures, we enriched our datasets with scRNAseq of 6217 cells from IFE and pilosebaceous units (PSU) microdissected from healthy scalp, area containing many follicular structures, of which 4565 cells passed the QC steps (SI Appendix, Fig. 8A). In order to perform high resolution clustering and refine cell annotations, we integrated our sequenced cells with one publicly available scRNAseq dataset derived from healthy scalp skin (20,561 epithelial cells; SI Appendix, Fig. 8B) (18). We used those cells as a template for the initial steps of clustering (Fig. 4A) and then analysed only our dataset comprising 11,966 epithelial cells from healthy skin (face and body) and BCC.

**Figure 4.**
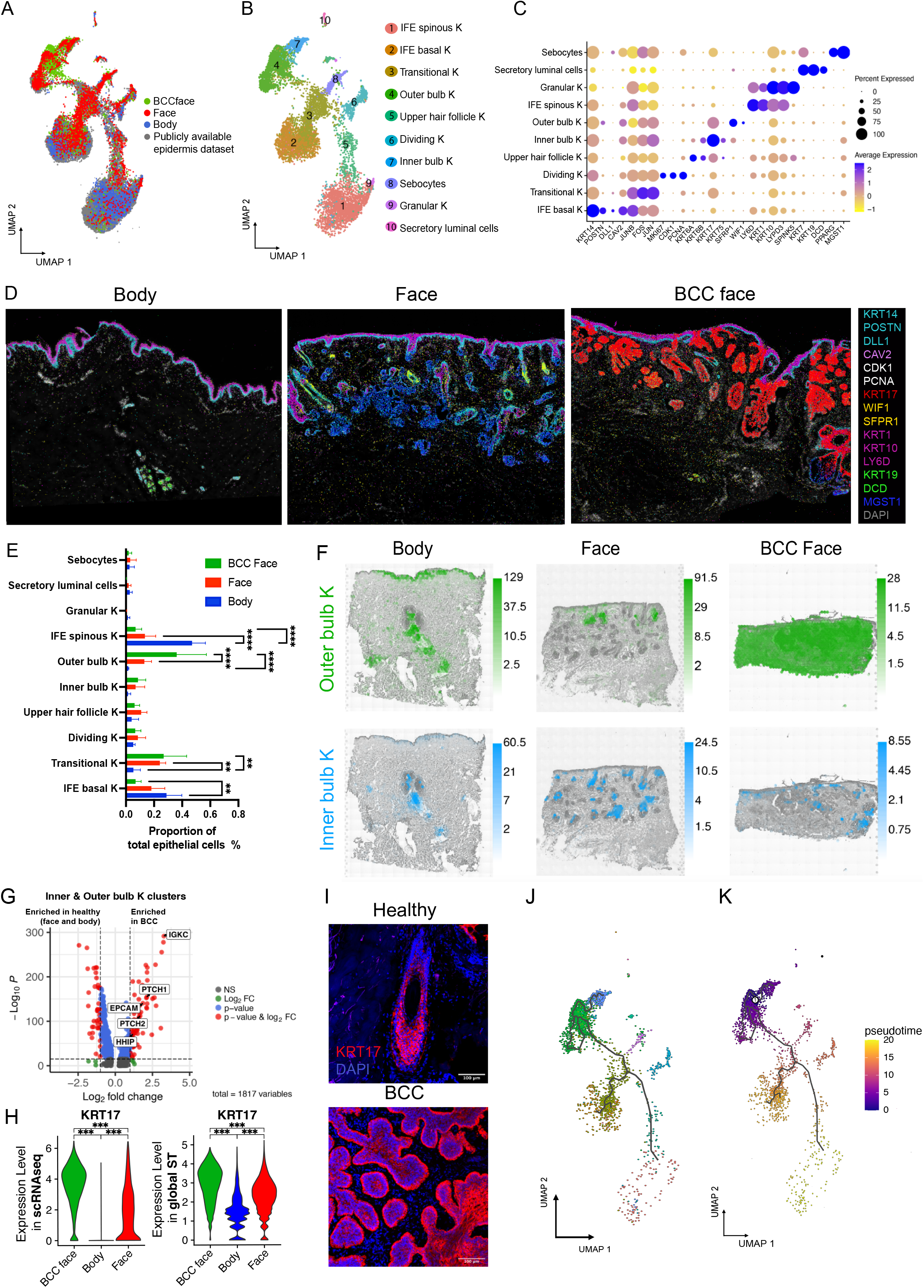
Lack of distinct epithelial cell populations between normal skin and BCC. A) UMAP of body, face skin (including IFE and PSU cells) and BCC integrated with a publicly available healthy scalp skin dataset to perform a high resolution clustering clustering. B) Unsupervised clustering identified a total of 10 clusters. C) Dot plot showing average log2 normalized expression of specific marker genes for each epithelial subpopulation. D) Spatial localisation of ISS reads for the different epithelial subpopulations in ISS images from a representative healthy body, face and BCC face sample (DAPI, grey). *KRT14, POSTN, DLL1* represented the IFE basal K; Dividing K co-expressed *CDK1* and *PCNA; KRT17* in the inner bulb cluster. *SFRP1; WIF1* marked the outer HF bulb. *KRT1, KRT10* and *LY6D* marked spinous IFE keratinocytes; *KRT19* and *DCD* marked sweat glands; *MGST1* marked sebaceous glands. Markers such as FOS, JUN or KRT6A and KRT6B, highly expressed by transitional K and upper HF K, were not included in our ISS gene panel and then not represented. E) Comparison of the abundance of the epithelial subpopulations between body, facial skin and BCC face. Statistically significant differences (One way ANOVA-test) are indicated (**<0.01, ****<0.0001). F) Cell2location prediction of keratinocyte populations corresponding to inner and outer bulb clusters. Predicted cell abundances shown by color gradients per spot. Three representative samples are shown. G) Volcano plot highlighting differential expression of *PTCH1/2, HHIP*, *EPCAM* and *IGKC* in BCC compared to healthy cells in the inner and outer bulb K clusters. H) Differential expression of *KRT17* in scRNAseq data (left panel) and global ST data (right panel) between BCC and healthy skin (face and body). Statistically significant differences (One way ANOVA-test) are indicated (***<0.001). I) Immunostaining of KRT17 in healthy face skin and BCC (DAPI, blue). J) Single cell trajectory gene analysis using Monocle 3 showing root nodes in BCC epithelial cells. K) Monocle pseudotime showing given lineage for each cell based on the distance from the cell of origin in BCC epithelial cells.

Unbiased clustering identified a total of 10 cell populations comprising undifferentiated (basal) and differentiated (spinous and granular) keratinocytes (K) specific to the IFE, dividing and transitional K, and K specific to adnexal structures such as upper HF K, inner and outer bulb K, sebocytes, and sweat gland secretory luminal cells (Fig. 4B, C; SI Appendix, Fig. 8C and Table 5). We plotted key markers in ISS (Fig. 4D).

Although we did not find any distinct epithelial cluster for BCC, markers in the hedgehog signalling pathway such as PTCH1/2, HHIP; EPCAM which is use as a diagnostic marker for BCC (41) and IGKC which is a immunologic marker of solid cancer (42) were highly expressed in BCC basal keratinocytes compared to healthy (SI Appendix, Fig. 8D). However, we found an increase of outer bulb K in BCC face compared to healthy. We also found an increase of transitional K in BCC and in healthy face compared to body skin. Surprisingly, there was also an increase of IFE basal and spinous K in healthy body compared to healthy face and BCC (Fig. 4E). ST deconvolution revealed epithelial clusters to localize to distinct regions within the epidermis and cutaneous appendages (SI Appendix, Fig. 8D). Tumour regions exhibited signatures of the inner bulb K and, predominantly, of the outer bulb K (Fig. 4F). Additionally, BCC cells in the inner and outer bulb K clusters showed high expression of PTCH1/2, HHIP (hedgehog signalling pathway), EPCAM and IGKC compared to healthy cells in these same clusters (Fig. 4G).

The most striking difference in BCC versus healthy was the high expression of KRT17 in the nodules and invasive tumours (Fig. 4D). This was confirmed in the scRNAseq data and independently in the ST data (Fig. 4H, SI Appendix, Fig. 8F, G). Immunostaining of KRT17 showed high expression in the inner bulb and inner root sheath and a low expression in the outer bulb and outer root sheath in healthy skin and high expression in BCC tumour islets (Fig. 4I). These results lead us to hypothesise that BCC tumour cells retain the transcriptional signatures of their cells of origin.

We then performed in silico single cell trajectory and pseudotime gene analysis using Monocle3 (43) in BCC keratinocytes (Fig. 4J, K) and independently in all the epithelial cells (SI Appendix, Fig. 8H). This trajectory inference method revealed numerous ‘root nodes’ in the outer bulb cluster in BCC (Fig. 4J). Pseudotime showed the given lineage for each cell based on the distance from the cell of origin, such as outer bulb K in the BCC (Fig. 4K).

Collectively, our data suggest that malignant epithelial cells of BCC could arise from cells of the inner and outer HF bulb. Selective expansion of *POSTN*+ fIbroblasts in BCC that are associated with the HF bulb in healthy skin strengthened this hypothesis.

## Discussion

Through a combination of scRNAseq, OCT, global ST and targeted ISS, we have created a cellular-resolution atlas of cell populations across multiple anatomical sites in healthy human skin and BCC. Using computational spatial mapping, we probabilistically inferred the localisation of cell populations in tissue sections. Mesenchymal cell subpopulations exhibited characteristic spatial distributions within the dermis and these same cell populations were repurposed in the stroma of BCC.

We found that analogous skin cell populations were present in body and facial skin despite differences in morphology and embryological origin. However, in keeping with a previous study (27), we observed differences in Hox gene expression in fibroblasts derived from the body versus the face. We additionally showed differences in expression of neural crest markers, not only in fibroblasts but also in other cell types including pericytes, Schwann cells, VEC and melanocytes. These findings indicate that fibroblast, pericyte, and VEC differentiation programmes can be concurrently activated in dermal lineages derived from cranial neural crest and somatic mesoderm. This is of particular interest in regard to oncogenic specificity. We know that oncogenic alterations to DNA are not transforming in all cellular contexts (7). It has been shown that anatomic position determines oncogenic responses due to pre-existing transcriptional programmes of the cell of origin (44).

In line with previous studies, we identified two main pericyte subpopulations (2, 17–20) and a SMC cluster that also had a peri-vascular location. *RGS5*+ and *TAGLN*+ pericytes corresponded to the populations reported by Reynolds *et al.*(*20*). Gene ontology and the location of *TAGLN*+ pericytes suggest a role in blood vessel development and contractility, similar to mesh/thin-strand pericytes previously described in other organs (45). In BCC, there was expansion of the *RGS5*+ pericytes and a reduction of *APOD*+ fibroblasts and *TAGLN*+ pericytes in BCC. The distribution of the *RGS5*+ pericyte subpopulation closely paralleled with VEC, whereas the distribution of the *APOD*+ fibroblasts and *TAGLN*+ pericytes was more focal. This contrasts with healthy skin where *APOD*+ fibroblasts, *RGS5*+ and *TAGLN*+ pericytes populations largely colocalize, implying selective expansion of the *RGS5*+ pericytes during tumour angiogenesis. Genetic ablation of *RSG5* in a mouse cancer model reduces tumour angiogenesis(46). These observations suggest that the *RGS5*+ pericytes may have a role in vascular remodelling during cancer neovascularization and therefore enhancing tumour progression.

We identified four main fibroblasts subpopulations localized to the following spatial contexts: in association with blood vessels (*APOD*+ fibroblasts); in the interstitial dermis with a concentration in the reticular dermis (*SFRP2*+ fibroblasts); concentrated adjacent to HFs, near the upper HF and in the papillary dermis (*PTGDS*+ fibroblasts); and near the HF bulb (*POSTN*+ fibroblasts). Whilst the number of fibroblast subpopulations identified in previous studies has varied, our demonstration of distinct spatial contexts for each of the populations sets a minimum bound on this number. It is probable that additional fibroblast populations will be identified in the future by enriching scRNAseq datasets in mesenchymal cells through increasing the depth of reads and the improvement of the spatial transcriptomic technologies.

In nodular and infiltrative BCCs, cancer-associated fibroblasts correspond to fibroblast subpopulations in healthy skin, with selective expansion of the *POSTN*+ fibroblasts. It is present adjacent to the tumour islands, suggesting that malignant epithelial cells that retain transcriptional signatures of the HF could induce the *POSTN*+ fibroblasts in adjacent fibroblasts or promote selective expansion of the healthy population. POSTN has been previously found in invasive BCCs (47, 48) but was not compared with healthy skin from the same sites.

The spatial localization of mesenchymal populations relative to epithelial appendages or epithelial tumours suggests that reciprocal signalling is important in the establishment and maintenance of cellular identity. We identified multiple subpopulations of keratinocytes within the skin that correlate with analogous subpopulations in adult mouse skin (49), although we did not find subpopulations specific to the hair bulge, but instead found two subpopulations which localize to the hair bulb. This could reflect differences in human skin architecture and the fact that 90% of human HFs are in anagen compared to mouse HFs (50).

The cell of origin of human BCC has been a subject of debate. Lineage tracing in the mouse supports a follicular origin (10, 51) and keratin expression in human BCC is in keeping with this model(9). *KRT17* was upregulated in BCC. In mice, *KRT17* is implicated in hair cycle regulation and tissue repair; genetic ablation inhibits tumour development and growth (52, 53). *KRT17* is also overexpressed in other cancers suggesting a more general role in tumorigenesis (54). However, murine studies additionally showed that activation of oncogenic hedgehog signalling in the bulge stem cells is not able to induce BCC(55) whereas this activation in the IFE can initiate BCC in mice(56). Spatial prediction of keratinocyte subpopulations and *in silico* lineage tracing support a follicular origin, specifically the bulb area. This data does not exclude the possibility of oncogenic mutations occurring in the IFE cells in human BCCs.

An understanding of the location and potential function of cell populations in healthy skin and how they are subverted in skin cancer is likely to inform therapeutic approaches. This could include inhibition, ablation or selective expansion of specific fibroblast or pericyte subpopulations. The rational design of such therapeutics will require a mechanistic understanding of the intrinsic transcriptional regulators of these cell populations and the mechanisms through which reciprocal signalling is coordinated with neighbouring cells.

In summary, our cellular-resolution map of human healthy skin and cutaneous malignancy is a valuable resource for the skin research and dermatology communities and forms a cornerstone of the Human Cell Atlas.

## Materials and methods

### Ethics

The study was sponsored by Guy’s and St Thomas’ NHS Foundation Trust and King’s College London and was subject to both institutional and external research ethics council (REC) review (REC reference 19/NE/0063). Following informed consent, excess skin samples were obtained from patients undergoing skin surgery.

### Single cell RNA sequencing and Data Analysis of scRNAseq

*SI Appendix*

### Spatial transcriptomics (10X Visium) and Data Analysis of ST

*SI Appendix*

### In situ sequencing and Data Analysis for ISS

*SI Appendix*

### Histology

*SI Appendix*

### Immunohistochemistry

*SI Appendix*

### RNAscope

*SI Appendix*

### Optical Coherence Tomography Imaging

*SI Appendix*

### Data availability

The raw and processed datasets for scRNAseq and Visium ST generated in the current study are available on ArrayExpress under ‘E-MTAB-13084’ and ‘E-MTAB-13085’ accessions and an interactive tool to display scRNAseq, ST and ISS data is available at https://spatial-skin-atlas.cellgeni.sanger.ac.uk/.

### Key Resources Table

*SI Appendix*

## Acknowledgements

This work was funded by grants to M.D.L. from the Wellcome Trust (211276/E/18/Z) and to F.M.W. from the Wellcome Trust (096540/Z/11/Z). *Additional acknowledgements in SI Appendix*.

