## Supplementary material for "Multi-scale spatial mapping of cell populations across anatomical sites in healthy human skin and basal cell carcinoma": SI Appendix

### Title Page

Affiliations:

Keywords:

### Supplementary Materials and Methods

#### Single cell RNA sequencing

Whole skin biopsies were kept on ice immediately after resection for no longer than 2hrs before processing. Characteristics of healthy donors and BCC patients (conditions, location, age, gender, number of cells) are summarised in SI Appendix, Fig. 2A, 6D. Biopsies were either directly processed for dissociation into cell suspension or kept in MACS Tissue Storage Solution (Miltenyi Biotec, Surrey, U.K, cat. no. 130-100-008) overnight. The enzymatic and mechanical dissociation of whole skin biopsies was performed using the Whole Skin Dissociation kit for human material (Miltenyi Biotec, Surrey, U.K, cat. no. 130-101-540). Larger specimens were reduced in size by pre-cutting into several small pieces prior to enzymatic digestion and the quantity of enzymes used was adjusted according to the size of the sample. Biopsies were incubated with the enzymes for 3 hours in a shaking water bath at 37°C. Cell suspensions were then filtered through 70µm cell strainer and centrifuged at 1500rpm for 10 minutes at 4°C. The collected cells were then frozen in 10%DMSO in Fetal Bovine Serum (FBS) and stored in liquid nitrogen.

To enrich epithelial populations, we microdissected pilosebaceous unit (PSU) cells from a healthy scalp skin donor. After treating the skin with Dispase (Sigma, Gillingham, Dorset, U.K.) overnight at 4°C, PSUs were microdissected under a dissecting stereomicroscope. PSUs and residual epidermis (IFE) were then separated into two tubes and digested with a mixture of trypsin-EDTA (Gibco, Paisley, U.K) and Versene (Gibco, 1.33:1) for 5 minutes at 37$^{o}$C. Cold DMEM with 10% FBS was added and the suspension was then passed sequentially through 70µm cell strainers in Falcon tubes. The collected cells were then frozen.

For library preparation, cells were thawed and resuspended in PBS with 2% FBS and DAPI (0.05mg/ml). Dead cells were removed through flow cytometry of the final cell suspension. Immediately after flow cytometry, ~20,000 to 100,000 live single cells were loaded onto a Chromium chip and libraries were prepared through droplet encapsulation on the Chromium controller (10X Genomics; Pleasanton, CA, USA) using the Single Cell 3' reagent kits following the manufacturer's protocol. Library quantification was performed using the Qubit dsDNA HS Assay Kit (Life Technologies, Waltham, MA, USA), and cDNA integrity was assessed using D1000 ScreenTapes (Agilent Technologies). Libraries were sequenced using an Illumina HiSeq 4000 device.

#### Data Analysis of scRNAseq

Sequencing data was analysed with Cell Ranger, v6.1.1 (10X Genomics). Downstream analyses were completed with Seurat (version 4.1.1). In order to combine our sequenced facial skin cells with body skin cells, we took advantage of publicly available scRNAseq datasets from Tabib *et al.*(17) and Solé-Boldo *et al.*(19). In order to characterise the epithelial populations of human skin, we combined our dataset from PSUs and IFE with a publicly available healthy scalp skin dataset(18). These datasets were combined using the merge() function from Seurat (Satija Lab, New York Genome Center, New York) and then a Seurat object was created with "min. cells = 3" and "min.features = 200." Cells with more than 5,000 expressed genes were removed to eliminate possible cell doublets, and cells with more than 5% mitochondrial reads were discarded to eliminate apoptotic cells. Seurat reference-based integration protocols were applied, which permitted the assembly of multiple, distinct single-cell RNA-sequencing data sets into an integrated reference and the correction of batch effects from interindividual differences. First, preprocessing of the data included log normalization of the unique molecular identifier counts and identification of the 2,000 most variable genes per sample. Then, using the FindIntegrationAnchors() function with default parameters and 20 canonical correlation analysis dimensions, integration anchors were identified across all samples. These anchors were used to integrate the data (using the IntegrateData() function, with the first 20 correlation analysis dimensions and default parameters). The integrated data were used for cell clustering and visualization with Seurat, which used the 2,000 most variable genes of the integrated data set as input. Next, data were scaled using the ScaleData() function. Principal component analysis (PCA) dimensions were calculated with the RunPCA() function. Unsupervised clustering of the data was performed with the FindNeighbors() and FindClusters() functions. The first 20 PCA dimensions were used for the FindNeighbors() function to construct a shared nearest-neighbour graph. of all the data. Then, the cells were clustered using the function FindClusters() with a shared nearest-neighbour modularity optimization-based clustering algorithm with a resolution of 0.65. Finally, the RunUMAP() function was used with default parameters and 20 PCA dimensions for visualization. FindAllMarkers() uses a Wilcoxon rank-sum test to identify the representative genes of each cluster. These representative genes were used to establish the cell identity of each cluster, in addition to well-known markers defined in the literature for human skin cell types. The average expression of a particular set of marker genes was used for cell type identification and projected into a Uniform Manifold Approximation and Projection (UMAP).

Differentially expressed genes were identified using the Wilcoxon sum rank test with a fold change cut-off of 0.25 (natural log scale) and only positive markers were returned. Genes that were detected in a minimum of 10% of cells in either of the two compared clusters were used.
Gene ontology (GO) term enrichment analysis was performed using ShinyGO 0.77 (57), inputting all significantly differentially expressed genes and assessing enrichment of Biological Process GO terms.

Putative cell-cell interactions were identified using CellPhoneDB (version 3.0.2; database v4.0.0) (36). CellPhoneDB uses a curated database of ligands, receptors and their interactions to identify ligand-receptor pairs that are highly expressed, and exhibit cell type specificity. We used fine-resolution cell types and "statistical_analysis" methods to identify the cell type-specific interactions. Interactions with p values of 0.05 or less, and mean normalized expression of 0.5 or higher, were considered in the further analysis. Microenvironments defined from the spatial transcriptomics experiments were used to refine the results of the analysis. Custom visualisations of specific interacting cell type pairs were generated using R/ggplot2.

We performed pseudotemporal trajectory and pseudotime using Monocle 3 v0.1.3 (60). The UMAP space from the Seurat package was used as an input of the reduced dimensional function in Monocle 3.

#### Spatial transcriptomics (10X Visium)

Fresh frozen OCT-embedded skin biopsies were cryosectioned at 10 µm, placed onto SuperFrost Plus glass slides (Thermo Fisher Scientific, Waltham, MA, USA) and stored for less than a week at -80˚C prior to high sensitivity library preparation. Optimal RNA integrity on the skin sections was assessed by RNAscope using three housekeeping genes with high (UBC) and medium (PPIB) expressors (SI Appendix, Fig. 3B). We also validated the integrity of the Visium library after sequencing: all samples presented good RNA quality (SI Appendix, Fig. 3C). However, we excluded 2 of the 32 10X Visium samples from further analysis due to poor tissue attachment to the slide.

Visium spatial gene expression slides and reagents were used according to manufacturer instructions (10X Genomics). Optimal permeabilization time for 10um skin sections was 20 minutes. cDNA libraries were quality controlled using the Agilent Bioanalyser. The cDNA libraries were sequenced on the Illumina HiSeq 4000 system, targeting 300 million reads per section with parameters 28cy R1, 8cy i7 index, 0cy i5 index, 91cy read 2. For each frozen sample, FASTQ files were manually aligned with corresponding H&E images and then analyzed with Space Ranger version 1.3.0 which uses the STAR genome aligner version v.2.5.1b and the human reference genome : Homo_sapiens (1000Genomes_hs37d5 + ensembl_75_transcriptome) [star] and Homo_sapiens (GRCh38_15_plus_hs38d1 + ensembl_90_transcriptome) [star].

#### Data Analysis of spatial transcriptomics (10X Visium)

Space ranger outputs were imported into R via Seurat V.4.1.1. All samples were merged to enable a comprehensive analysis. Spots with more than 30% of mitochondrial genes and less than 200 genes per spot were filtered out. Additionally areas identified as necrotic or damaged were identified by a dermatologist experienced in the interpretation of skin frozen sections (M.D.L) and excluded from further analysis.

Raw counts were normalized with the SCTransform() function and dimensionality and clustering steps are the same as in the standard scRNAseq workflow. Data were visualised in UMAP space using DimPlot() or overlaid on the H&E images using SpatialDimPlot() or SpatialFeaturePlot() functions. In order to deconvolute scRNAseq clusters into 10X Visium sections, we used *cell2location v0.1 (22)*. This algorithm comprises two steps. First, we trained a negative binomial regression model to estimate reference transcriptomic profiles for all the cell types profiled with scRNA-seq in the organ. We excluded very lowly expressed genes using the filtering strategy recommended by cell2location authors (cell_count_cutoff=5, cell_percentage_cutoff SFRP2+ fibroblasts=0.03, nonz_mean_cutoff=1.12). Donor information was included as a categorical covariate. Training lasted for 250 epochs and reached convergence according to manual inspection. Next, we estimated the abundance of cell types in the ST slides using reference transcriptomic profiles of different cell types. All slides were analysed jointly. The following cell2location hyperparameters were used: (1) expected cell abundance (N_cells_per_location) = 30; (2) regularisation strength of detection efficiency effect (detection_alpha) = 20. The training was stopped after 50,000 iterations. All other parameters were used at default settings. Cell2location estimates the posterior distribution of cell abundance of every cell type in every spot. Posterior distribution was summarised as 5% quantile, representing the value of cell abundance for which the model has high confidence.

To identify microenvironments of colocalizing cell types, we used non-negative matrix factorisation (NMF). We first normalized the matrix of estimated cell type abundances by dividing it by per-spot total abundances. Resulting matrix Xn of dimensions $n \times c$, where n is the total number of spots in the Visium slides and c is the number of cell types in the reference was decomposed as $X$n = $WZ$, where

W is a $n \times d$ matrix of latent factor values for each spot and Z is a $d \times c$ matrix representing the fraction of abundance of each cell type attributed to each latent factor. Here latent factors correspond to tissue microenvironments defined by a set of colocalized cell types. We use the NMF (61) package for R, setting the number of factors d = Hi NN and using the default algorithm (Brunet2004). NMF coefficients were normalized by a per-factor maximum. We ran NMF 100 times and constructed the coincidence matrix. Then we selected the best run based on lower mean silhouette calculated on coincidence matrix, if more than one run have minimal mean silhouette we selected one with smaller deviance (as reported by nmf function).

For celltype abundance correlation analysis we used per-spot normalized Xn matrix. PCC was calculated for each celltype pair and each sample.

#### *In situ* sequencing

Fresh frozen OCT-embedded skin biopsies were cryosectioned as 10 µm sections and placed onto SuperFrost Plus glass slides (Thermo Fisher Scientific, Waltham, MA, USA) and stored for less than a week at -80˚C prior to high sensitivity library preparation. Samples were fixed with 4% formaldehyde (Merck, Gillingham, Dorset, U.K, ref. F1635) for 30 minutes and then permeabilized for 90 seconds with 0.1 mg/mL pepsin (Merck, Gillingham, Dorset, U.K, ref. 10108057001) in 0.1M HCl (VWR, Leicestershire, U.K, ref. 20255.290). RNA integrity and assay conditions were assessed using MALAT1 (high expressor) and RPLP0 (medium expressor) housekeeping genes only (SI Appendix, Fig. 4E). For library preparation, chimeric padlock probes (targeting directly RNA and containing an anchor sequence as well as a gene-specific barcode) for a custom panel of 81 genes (SI Appendix, Table 2) as well as of 90 genes from 2 pre-defined immune panels (SI Appendix, Table 3, 4) were hybridized overnight at 37°C, then ligated before the rolling circle amplification was performed overnight at 30°C using the High Sensitivity Library Preparation kit for CARTANA technology (10xGenomics) following the manufacturer's instructions. All incubations were performed in SecureSealTM chambers (Grace Biolabs, Bend, OR, USA). To quench autofluorescence background, TrueView (SP-8400 VectorLabs) was used for 2 minutes at room temperature. For tissue sections mounting, Slow Fade Antifade Mountant (Thermo Fisher Scientific, Waltham, MA, USA, ref S36936) was used for optimal handling and imaging. Quality control of the library preparation was performed by applying anchor probes to detect simultaneously all rolling circle amplification products from all genes in all panels. Anchor probes are labelled probes with Cy5 fluorophore (excitation at 650 nm and emission at 670 nm).

All samples passed quality control and were sent to CARTANA Sweden (a subsidiary of 10X Genomics), for *in situ* barcode sequencing, imaging and data processing. Briefly, adapter probes and sequencing pools (containing 4 different fluorescent labels: Alexa Fluor® 488, Cy3, Cy5 and Alexa Fluor® 750) were hybridized to the padlock probes to detect the gene-specific barcodes, through a sequence specific signal for each gene specific rolling circle amplification product. This was followed by imaging and 6 iterations were performed to permit decoding of all genes in the panel. Raw data consisting of 40x images from 5 fluorescent channels (DAPI, Alexa Fluor® 488, Cy3, Cy5 and Alexa Fluor® 750) were each taken as z-stack and flattened to 2D using maximum intensity projection. After image processing and decoding, the results were summarized in a csv file and gene plots were generated using MATLAB.

#### Data Analysis for ISS

Analysis for ISS data was performed for each sample following the SSAM pipeline (v.1.0.2) (31) with minor modifications to the code. Gene reads were loaded and their coordinates transformed from pixels into micrometers (0.16 µm per pixel) to create a SSAM dataset object. The locations of the gene reads were converted into mRNA density through a Kernel Density Estimation (KDE) using a Gaussian kernel and a bandwidth of 2.5. Local maxima of the mRNA densities were found using a total gene expression threshold of 0, a per gene expression threshold of 0, and a search size of 3. Local maxima were employed to calculate the variance stabilization parameters with the sctransform package (v0.3.4) (58), which were subsequently used for gene expression normalization. Cell type identification was performed following SSAM guided analysis and using either the merged face and body or face and BCC processed scRNAseq data as the reference dataset. The raw gene counts for both reference datasets were normalized using sctransform and the average gene expression per cell type was calculated. Local maxima vectors were mapped to the cell type clusters with the most similar gene expression in the scRNAseq reference dataset, using a correlation threshold of 0.15 and a threshold of vector normalisation of 0.0015.

#### Histology

Skin tissue samples were embedded in Tissue-Tek O.C.T. (Life Technologies, Waltham, MA, USA) and stored at –80°C prior to sectioning. Skin sections were sectioned at 10–16 µm using a Thermo Cryostar Nx70 (Thermo Fisher Scientific, Waltham, MA, USA) and placed onto SuperFrost Plus glass slides (Thermo Fisher Scientific, Waltham, MA, USA, ref J2800AMN2). Sections were stained with Harris's hematoxylin solution for 20s at room temperature and were then rinsed in tap water. Next, 0.225% Acid Alcohol (acetic acid and ethanol) in water was used to differentiate the tissue for 10s. Then, skin sections were rinsed with tap water. In the bluing step, the tissue was soaked in Scott's tap water (ref: 3802900, Leica Biosystems, Milton keynes, UK) for 10s and then rinsed with tap water. Staining was performed with 0.125% eosin Y ethanol solution for 10s. Finally, skin sections were dehydrated in IMS for 20s, then cleared in Xylene 30s and mounted with CV mounting medium (ref : 14046430011, Leica biosystem, Germany).

Epidermal thickness (µm) was quantified using a length measuring tool in Fiji software (62). At least 20 measurements were performed per H&E image. The numbers of hair follicles, sebaceous glands and eccrine sweat glands were manually counted per mm length of epidermis. The mean values for epidermal thickness were calculated for each area and plotted on a graph for statistical analysis.

#### Immunohistochemistry

Skin tissue sections were embedded and sectioned as described above. Sections were fixed in 4% paraformaldehyde for 5 minutes, permeabilized in 0.2% Triton (Sigma-Aldrich, St. Louis, MO, USA, ref T92841L) in PBS for 15 minutes and then blocked in blocking buffer solution containing 10% serum, 0.2% fish skin gelatin, 0.1% BSA, and 0.5% Tween-20 (all from Sigma-Aldrich, St. Louis, MO, USA) in PBS and labelled with primary antibodies diluted in blocking buffer overnight at 4°C. Sections were washed with PBS and then labelled with secondary antibodies (all from Thermo Fisher Scientific, Waltham, MA, USA) for 1 hour at room temperature, washed with PBS, and mounted with ProLong™ Gold Antifade Mountant with DAPI (Thermo Fisher Scientific, Waltham, MA, USA, ref P36931).

#### RNAscope

Skin tissue sections were embedded and sectioned as described above. RNAscope experiments were performed using the RNAscope Multiplex Fluorescent Detection Kit v2 (ACDBio, Newark, California, USA, cat. no. 323100) according to the manufacturer's instructions. RNA integrity on the skin sections was assessed by RNAscope with housekeeping control probes (high (UBC), medium (PPIB) and low (POLR2A) expressors). Probes against targeted human mRNA molecules were used (all from ACDBio catalog probes, Newark, California, USA). Opal dyes (Akoya Biosciences, Marlborough, Massachusetts, USA) were used at a dilution of 1:1,000 for the fluorophore step to develop each channel: Opal 520 Reagent Pack (FP1487001KT), Opal 570 Reagent Pack (FP1488001KT) and Opal 650 Reagent Pack (FP1496001KT). Nuclei were counterstained with 4',6-diamidino-2-phenylindole and mounted using ProLong Gold Antifade Mountant (ThermoFisher, Canoga Park, California, cat. no. P36930). Slides were imaged with a Nikon A1 upright confocal microscope (Nikon, Tokyo, Japan) using a 20 dry lens and were further processed using Fiji software (62).

#### Optical Coherence Tomography Imaging

*In vivo* two-dimensional (2D) and three-dimensional (3D) angiographic optical coherence tomography (OCT) images were acquired using a commercially available OCT scanner (VivoSight Dx; Michelson Diagnostics, Kent, UK) on 16 healthy individuals and 11 patients. Images were obtained by placing the hand-held probe directly on the skin (scan area : 6 mm × 6 mm) using a fitted plastic spacer for stability and carefully avoiding compression of the skin. A full scan took 2 minutes. Angiographic OCT scanning generated greyscale images showing the structure of the skin and red areas show the vasculature based on blood flow motion in dermal vessels detected by angiographic OCT acquisition (SI Appendix, Fig. 1B). 3D reconstructions from 500 image frames were generated using Imaris software version 9 (Oxford instrument) for all samples. Total vascular density was analysed using the surface area modality in Imaris software version 9. Papillary and reticular vascular density were calculated using the threshold of mean pixel value in binarized max projection OCT frames.

### Key Resources Table

| Reagent or Resource | Source | Identifier |
|---|---|---|
| **Antibodies** | | |
| Cytokeratin 17 Monoclonal Antibody (E3) | Thermo Fisher Scientific | MA5-13539 |
| CLDN5 polyclonal antibody | ABCAM | AB131259 |
| Donkey anti-rabbit IgG (H+L) Alexa Fluor Plus 647 | Invitrogen | A-32795 |
| Donkey anti-mouse IgG (H+L) Alexa Fluor Plus 594 | Invitrogen | A-32744 |
| Goat anti-chicken IgY (H+L) Alexa Fluor 647 | Invitrogen | A-21449 |
| Goat anti-mouse IgY (H+L) Alexa Fluor 647 | Invitrogen | A-11001 |
| Vimentin monoclonal antibody | Leica | NCL-L-VIM-572 |
| TAGLN polyclonal antibody | Abcam | ab10135 |
| RGS5 polyclonal antibody | Abcam | ab196799 |
| CD146 (NCAM) Mouse monoclonal | Abcam | ab24577 |
| **Critical Commercial Assays** | | |
| Chromium Single Cell 30 Library & Gel Bead Kit v2 | 10x Genomics | PN-120267 |
| RNAscope® Multiplex Fluorescent Reagent Kit | Bio-Techne | 323100 |
| Opal 520 Reagent Pack | Akoya Biosciences | FP1487001KT |
| Opal 570 Reagent Pack | Akoya Biosciences | FP1488001KT |
| Opal 650 Reagent Pack | Akoya Biosciences | FP1496001KT |

| | | |
|---|---|---|
| **RNAscope probes** | | |
| HS-ACTA2 | Bio-Techne | 311811 |
| HS-APOD | Bio-Techne | 445171 |
| HS-DES | Bio-Techne | 403041 |
| HS-POSTN | Bio-Techne | 409181 |
| HS-PPIB - RNAscope® 3-plex Positive Control Probe | Bio-Techne | 320861 |
| HS-PTGDS | Bio-Techne | 431471 |
| HS-SFRP2 | Bio-Techne | 476341 |
| Hs-UBC - RNAscope® 3-plex Positive Control Probe | Bio-Techne | 320861 |
| **Software and Algorithms** | | |
| Cell2location | (22) | https://github.com/BayraktarLab/cell2location |
| Cell Ranger Analysis Pipeline v6.1.1 | 10X Genomics | https://10xgenomics.com/ |
| CellPhoneDB | (36) | https://github.com/ventolab/CellphoneDB/ |
| ShinyGO 0.77 | (57) | http://bioinformatics.sdstate.edu/go/ |
| MATLAB R2022a | MathWorks | https://mathworks.com/ |
| MATLAB script to decode intensity information in situ sequencing data | (30) | https://github.com/Moldia/in_situ_seq |
| sctransform v0.3.4 | (58) | https://github.com/satijalab/sctransform |
| Seurat v4.1.1 | (59) | https://satijalab.org/seurat/install.html |
| Monocle3 | (43) | https://github.com/cole-trapnell-lab/monocle3 |
| Space Ranger Analysis Pipeline v1.3.0 | 10X Genomics | https://10xgenomics.com/ |
| SSAM v1.0.2 | (31) | https://github.com/HiDiHlabs/ssam |

### Supplementary Figure Legends

#### Supplementary Figure 1

**Variation in skin morphology and skin vasculature across anatomical sites**

A) Histological sections of body (upper panel) and facial (lower panel) healthy skin (H&E stain, representative examples are shown).

B) Quantification of differences in epidermal thickness and density of sebaceous glands, hair follicles and eccrine sweat glands per mm length of epidermis. Significant differences (One way ANOVA-test) are indicated (**<0.01,****<0.0001).

C) Optical coherence tomography (OCT) imaging of skin microstructure in vivo is compared for body (upper panel) and facial (lower panel) skin. The intensity of the skin reflectance is shown in greyscale and speckle contrast imaging of blood vessels in red.

D) Quantification of skin vasculature density across a total of 6 anatomical sites: 3 from the body (Abdomen, Forearm, Back) and 3 from the face (Forehead, Nose and Ear) across 16 individuals (upper panel). Comparison of vasculature density in papillary and reticular dermis across a total of 6 anatomical sites (lower panel). Each filled circle represents the result of a single observation. Statistically significant differences (One way ANOVA-test) are indicated (* <0.05, **<0.01, ***<0.001, ****<0.0001).

E) Further vasculature network architecture quantification was performed on face skin areas only, because the vascular network was continuous, permitting quantification of segment length, number of segments, tube thickness and number of branching. No statistically significant differences were found (One way ANOVA-test).

#### Supplementary Figure 2

**Single cell RNA sequencing of human healthy skin and BCC**

A) The source and demographics of the skin samples incorporated into our combined single cell RNA sequencing dataset are described in this table.

B) UMAP plot illustrating integrated single cell sequencing from all the donors. Color indicates the donor source of the cell.

C) Quality control plots showing percent of UMI counts in mitochondrial genes per scRNAseq sample (upper panel) and frequency distribution of UMI counts (log1p-transformed) per scRNAseq sample (lower panel).

D) UMAP plots showing the contribution of each condition (BCC from the face, healthy face and body) to the distinct clusters identified in the integrated dataset. Color indicates the identity of the scRNAseq clusters.

E) Dot Plot showing marker genes specific to each skin cluster. For each cluster, the percentage of cells expressing the marker (diameter) and the average log2 normalized expression (color) is shown.

F) Quantification of chondrocytes (upper panel) and skeletal muscle cells (lower panel) according to facial anatomical sites. Since chondrocytes were only detected in an ear skin sample and skeletal muscle cells in a forehead skin sample statistical significance could not be calculated.

### Supplementary Figure 3

**Developmental gene signatures in human skin cells vary across anatomical sites**

A) Expression of selected somatic mesoderm Hox genes (upper panel) and neural crest mesenchymal genes (lower panel) in our integrated scRNAseq are illustrated in dot plots for which the diameter of the circle corresponds to the percentage of cells expressing the gene and the color reflects the average log2 normalized expression.

B) Average expression of selected somatic mesoderm Hox genes (upper panel) and neural crest mesenchymal genes (lower panel) in global ST profiling.

C, D) Expression of selected neural crest mesenchymal genes in scRNAseq data according to cell types (C) and expression of selected somatic mesoderm Hox genes in scRNAseq data according to cell types (D). Average log2 normalized expression for each cell type is compared between body (blue), facial (red) and BCC (green) conditions. Hox genes were preferentially expressed in body fibroblasts, pericytes, SMC, chondrocytes and VEC. Mesenchymal neural crest genes were preferentially expressed in facial (healthy and BCC) fibroblasts, pericytes, SMC and chondrocytes. Other populations, including melanocytes, Schwann cells and keratinocytes, exhibited less striking differences in marker expression between body, face and BCC face. Skin immune cell populations did not exhibit differential expression of these genes.

### Supplementary Figure 4

**Spatial transcriptomics and *in situ* sequencing of human healthy skin and BCC**

A) The source and demographics of the skin tissue sections used for ST (10X Visium) (black and red) and ISS (red) are described in this table.

B) Multiplex RNAscope of housekeeping gene expression (PPIB and UBC) has been performed on all the skin sections selected of ST by 10X Visium in order to check the presence of high quality RNA throughout tissue sections. One representative sample is shown (healthy face sample).

C) Quality control plots showing percent of UMI counts in mitochondrial genes per ST sample (upper panel) and frequency distribution of UMI counts (log1p-transformed) per ST sample (lower panel).

D) UMI counts in body, face and BCC face ST skin sections used for 10X Visium protocol. Three representative samples are shown.

E) Prior to final library preparation for ISS, RNA integrity and assay conditions were assessed using MALAT1 (high expressor, not shown) and RPLP0 (medium expressor, shown here) housekeeping genes. Three representative samples are shown.

F) Skin specific 165 gene panel in all the sections used for ISS. Each gene is represented by different color dots. Three representative samples are shown.

### Supplementary Figure 5

**Mapping of the main human skin cell types using global spatial transcriptomics and targeted *in situ* sequencing**

A) Using cell2location pipeline, scRNAseq clusters corresponding to the epidermis (suprabasal & basal keratinocytes), dermal stroma (fibroblasts) and endothelium (endothelial cells, pericytes and SMC) were projected onto the global ST skin sections isolated from the same skin biopsies.

B) Three representative ISS images from body, face and BCC face conditions representing the main cell types in human skin highlighting skin tissue architectures and demonstrating the sensitivity of ISS technology. *KRT14, KRT5, KRT10* and *KRT1* marked the epidermis and healthy epithelial structures and tumours within the dermis (blue). *EMB, MGST1, KRT19* and *DCD* marked the appendages such as sebaceous glands and sweat glands (pink). *DCN, COL1A1* and *PDGFRA* marked the dermal stroma (green). *PECAM, CLDN5, ACKR1* and *ACTA2* marked the dermal vasculature (red), DAPI (gray).

C) Using the SSAM pipeline, the same scRNAseq clusters with the same color code were projected onto the same ISS skin sections demonstrating single cell spatial mapping. As in the scRNAseq data, no appendage cluster could be found, therefore we did not spatially project those cells.

### Supplementary Figure 6

**Additional analysis of pericyte populations in healthy skin and basal cell carcinoma**

A) OCT imaging of vascular network of human BCC in vivo. En-face view showing speckle contrast imaging of blood vessels in red via 3D computational reconstructions. Image size 6 mm × 6 mm; scale bar: 500um.

B) Gene ontology terms of different peri-vascular subpopulations (RGS5+ pericytes, TAGLN+ pericytes, SMC).

C) Expression of markers specific to the different pericyte subpopulations on global ST skin sections of human skin from face and body areas and in BCC. Three representative samples are shown.

D) Spatial localisation of ISS reads specific to the different pericyte subpopulations in skin sections of human skin from face and body areas and in BCC. Three representative samples are shown.

E) Heatmap showing the total number of interactions between *RGS5+, TAGLN+* pericytes, SMC and VEC in the scRNAseq dataset obtained with CellPhoneDB a package looking for ligand–receptor interactions in scRNAseq data in combination with predicted microenvironment from ST data (methods). Dark-blue squares (0 interactions) and dark red the maximum number of interactions that was found here (>100).

F) Dotpots representing selected ligand–receptor interactions between pairs of *RGS5*+ pericytes, *TAGLN*+ pericytes and VEC. Only the interaction pairs that are most likely based on predicted microenvironment from ST data are shown. Results are not symmetrical; for example: *RGS5*+ pericytes|VEC with *CCL2_ACKR1* means CCL2 ligand is expressed in *RGS5*+ pericytes, and *ACKR1* receptor in VEC; for VEC | *RGS5*+ pericytes pair, the roles are switched. Thus, each pair is present twice. The total number of interactions found are filtered compared to heatmaps, and only include interactions for which at least one pair of cell types has mean expression of >= 0.5.

### Supplementary Figure 7

**Additional analysis of fibroblast populations in healthy skin and basal cell carcinoma**

A) Gene ontology terms of different fibroblast subpopulations (APOD+ fibroblasts, SFRP2+ fibroblasts, PTGDS+ fibroblasts, POSTN+ fibroblasts).

B) Expression of markers specific to the different fibroblast subpopulations on global ST skin sections of human skin from face and body areas and in BCC. Three representative samples are shown.

C) Spatial localisation of ISS reads specific to the different fibroblast subpopulations in skin sections of human skin from face and body areas and in BCC. Three representative samples are shown.

D) Heatmap showing the total number of interactions between APOD+, SFRP2+, PTGDS+, POSTN+ fibroblasts, VEC and Basal K in the scRNAseq dataset obtained with CellPhoneDB in scRNAseq data in combination with predicted microenvironment from ST data (methods). Dark-blue squares (0 interactions) and dark red the maximum number of interactions that was found here (>150).

E) Dotpots representing selected ligand–receptor interactions between pairs of APOD+, PTGDS+, POSTN+ fibroblasts and VEC. Only the interaction pairs that are most likely based on predicted microenvironment from ST data are shown. Results are not symmetrical; for example: APOD+ fibroblasts|VEC with CCL2_ACKR1 means CCL2 ligand is expressed in APOD+ fibroblasts and ACKR1 receptor in VEC; for VEC | APOD+ fibroblasts pair, the roles are switched. Thus, each pair is present twice. The total number of interactions found are filtered compared to heatmaps, and only include interactions for which at least one pair of cell types has mean expression of >= 0.5.

### Supplementary Figure 8

**Additional analysis of keratinocyte populations in healthy skin and BCC**

A) Microdissected interfollicular epidermis (IFE, light orange) and pilosebaceous units (PSU, dark orange) from healthy scalp skin and integrated in our combined epidermal dataset. We also integrated our sequenced cells with a publicly available healthy scalp skin dataset, in order to perform a high resolution clustering we used those cells as a template for the initial steps of clustering, they are represented in gray (left panel). Quality control plots showing percent of UMI counts in mitochondrial genes and frequency distribution of UMI counts (log1p-transformed) per scRNAseq sample conditions (right panel).

B) UMAP plots showing the contribution of our previous keratinocyte clusters such as basal and suprabasal keratinocytes in our combined epidermal dataset.

C) UMAP plots showing the contribution of each condition (BCC from the face, healthy face and body) to the distinct clusters identified in the integrated epidermal dataset. Color indicates the identity of the scRNAseq clusters.

D) Volcano plot highlighting differential expression of PTCH1/2, HHIP (hedgehog signalling pathway), KRT17, EPCAM and IGKC in BCC basal keratinocytes compared to healthy basal keratinocytes.

E) Prediction of epidermal cell populations onto global ST skin sections of human skin (face and body) and in BCC with cell2location (Methods). Three representative samples are shown. Skin architecture (H&E) is shown in grey; signal intensity is shown by the different colors indicated in each scRNAseq clusters.

F) UMAP plots showing average log2 normalized *KRT17* expression in our epidermal combined dataset.

G) Expression of *KRT17* in ST skin sections of facial, body skin and BCC. Signal intensity corresponds to color. Three representative samples are shown.

H) Monocle trajectory analysis of our epidermal combined dataset.

### Supplementary Table Legends

#### Supplementary Table 1

**Top twenty differentially expressed genes within all skin and BCC cell clusters**

Table showing significantly differentially expressed genes (top20) in each cluster identified through unsupervised clustering in all healthy skin cells (face and body) and BCC cells. Sorted from the highest average log2 normalized gene expression and adjusted p-value.

#### Supplementary Table 2

**List of custom genes for ISS**

List of the 81 custom panel genes, specific to skin subpopulations identified by scRNAseq, for the ISS.

#### Supplementary Table 3

**List of predefined Immune General gene panel for ISS**

List of the predefined panel Immune General (I1A 0.2) genes for the ISS.

#### Supplementary Table 4

**List of predefined Immune Regulation gene panel for ISS**

List of the predefined panel Immune Regulation (I3D 0.2) genes for the ISS.

#### Supplementary Table 5

**Top ten differentially expressed genes in epidermal cell clusters**

Table showing significantly differentially expressed genes (top10) in each cluster identified through unsupervised clustering in epidermal cells in healthy (face and body) and BCC cells. Sorted from the highest average log2 normalized gene expression and adjusted p-value.

### Additional Acknowledgements

We thank Marina de Paula-Silva and Christina Philippeos at King's College London (KCL) for their help with skin sample collection and sectioning. We thank Shanawaz Ali, Kalle Sipila, Matteo Vietri Rudan and Miguel Bernabe Rubio at KCL and Anais Julien at the Karolinska institutet, Sweden, for helpful discussions. We thank Christos Tziotzios for collaboration in sourcing skin scalp samples. We thank Eva Wozniak and Paul Stevens at the Genome Centre, Blizard Institute, QMUL, and Han Lu, and Dasha Freydina at the Genomics Centre, KCL, for assistance with library preparation. We thank the genomic platform and the cellular genetics informatics team at the Wellcome Sanger Institute, Hinxton, in particular Elena Prigmore, Stijn van Dongen and Steven Leonard for sequencing the libraries. We thank Morgane Rouault, Dabashish Chitnis and Hao Xu at 10X genomics, Sweden, for the sequencing and decoding for our ISS slides. We thank staff at the Advanced Sequencing Facility at the Francis Crick Institute, particularly Robert Goldstone and Amelia Edwards. We thank all members of the Dermatological Surgery Unit, Guy's and St Thomas' NHS Foundation Trust in particular Raj Mallipeddi, Faisal Ali, Jack Mann and Clare Kiely for assistance with recruitment of patients. Finally, we would like to thank all the patients who contributed skin samples without which this project would not have been possible.
