## Supplementary figures and images for "Multi-scale spatial mapping of cell populations across anatomical sites in healthy human skin and basal cell carcinoma"

### Supplementary Figure 1

#### Supplementary Figure 1

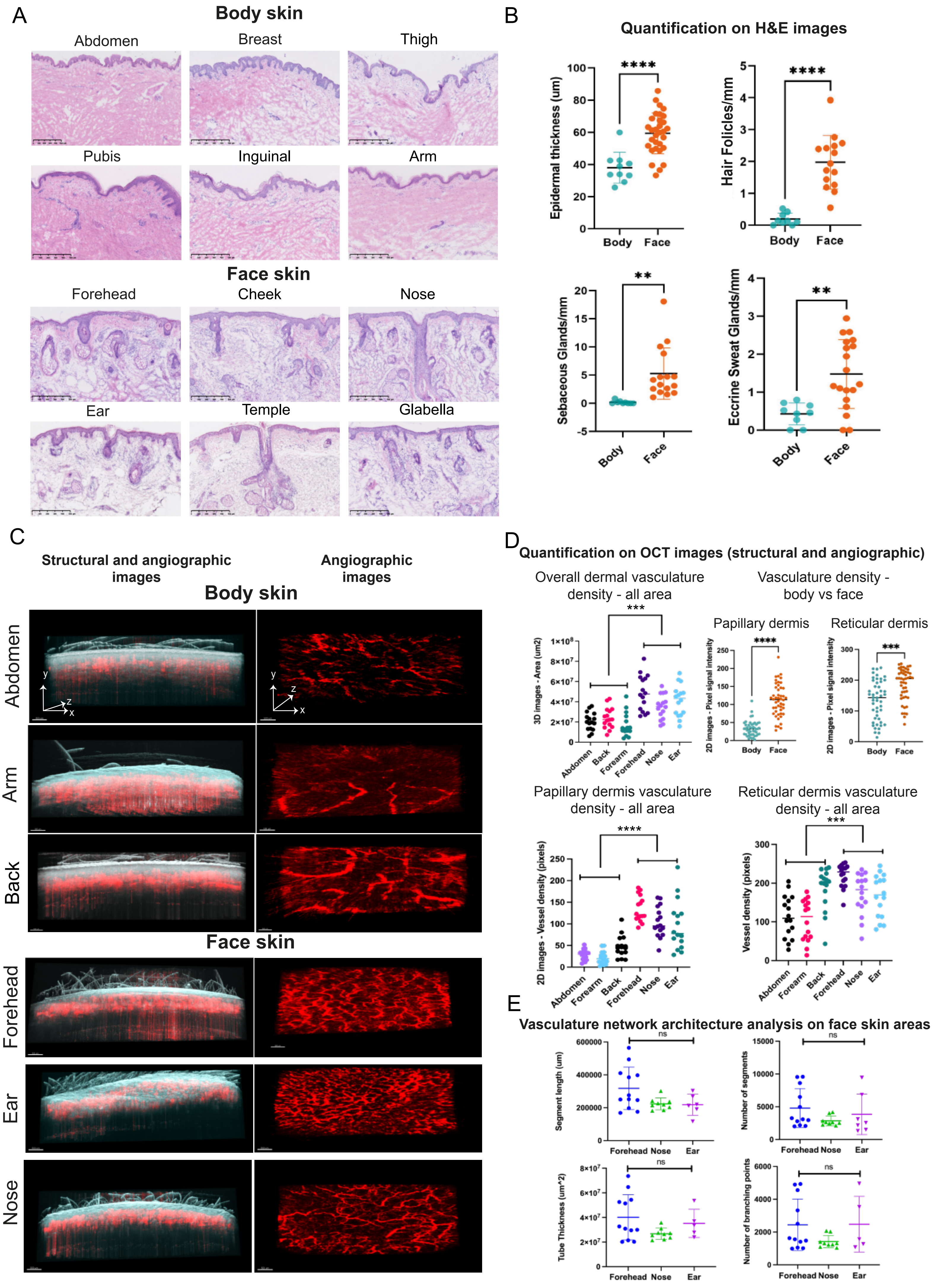

### Supplementary Figure 3

Supplementary Figure 3

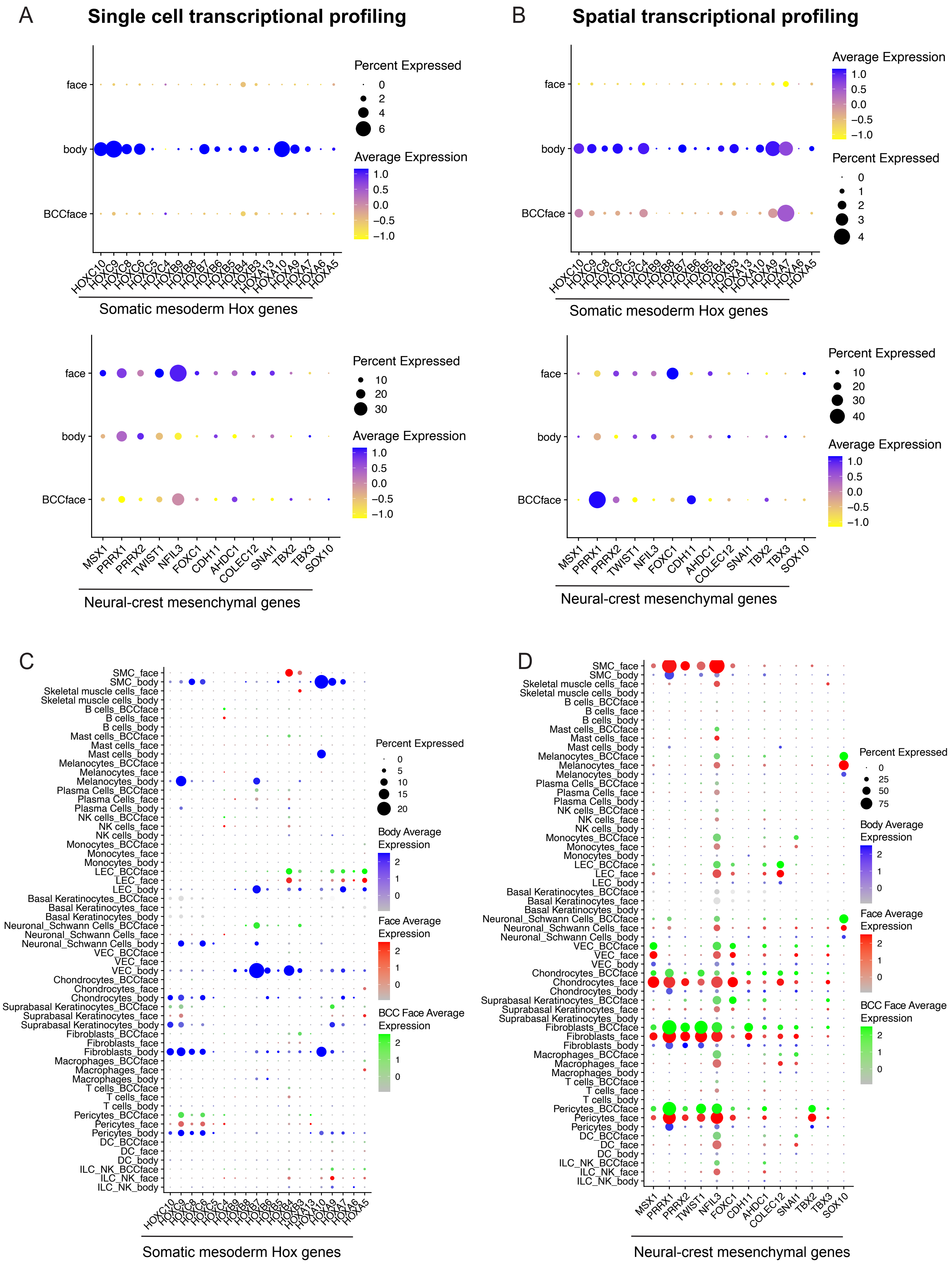

### Supplementary Figure 5

#### Supplementary Figure 5

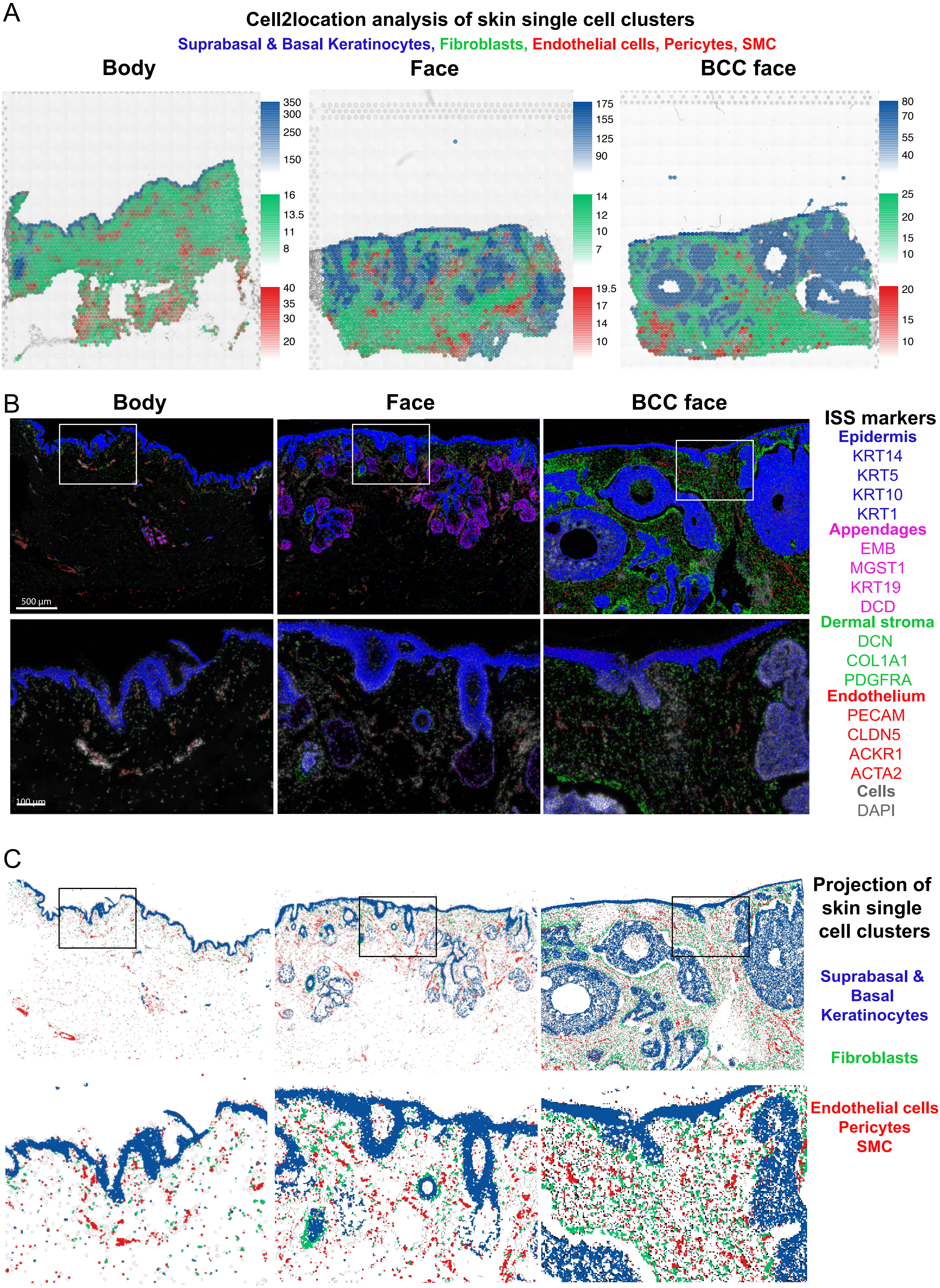

### Supplementary Figure 6

#### Supplementary Figure 6

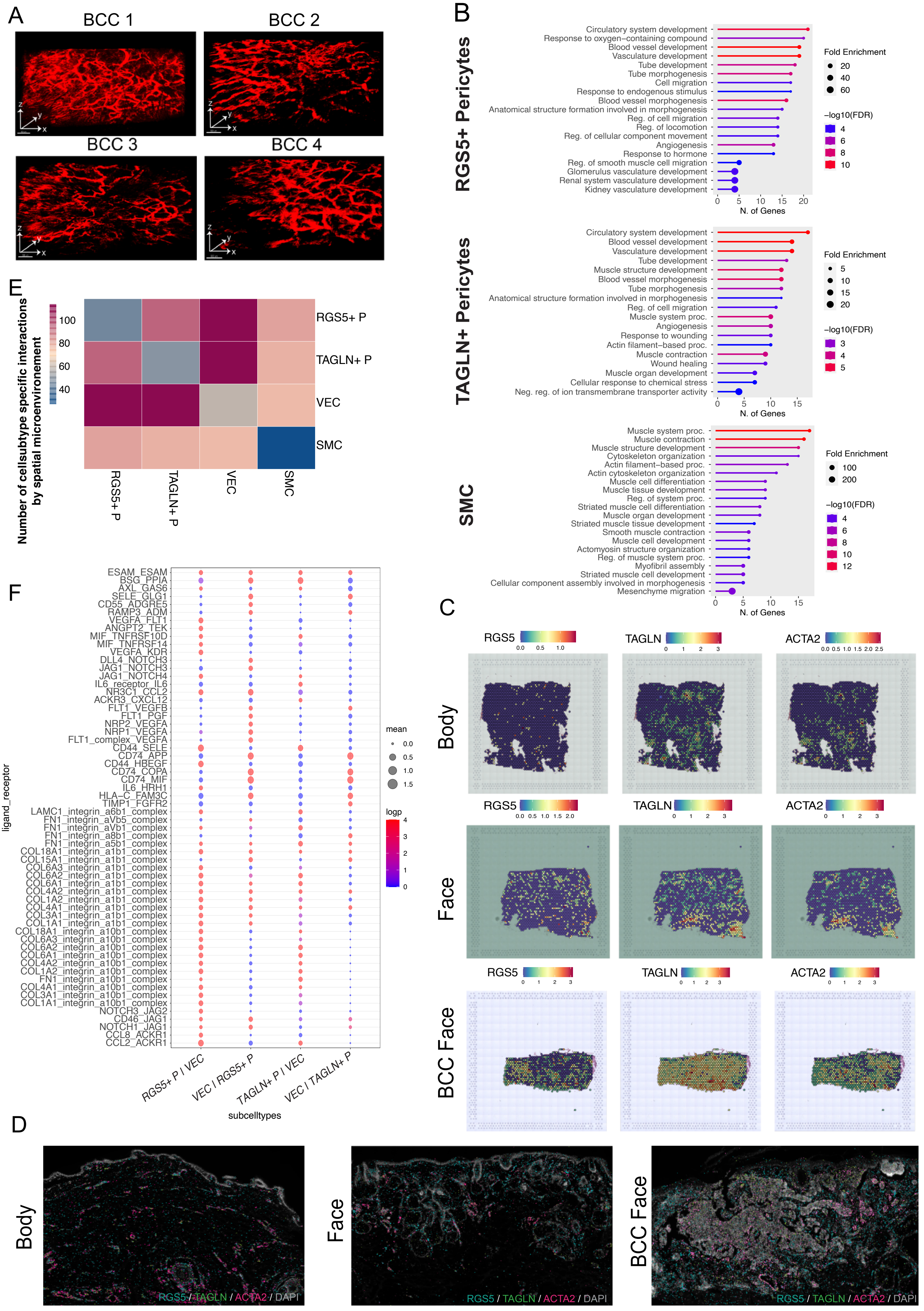

### Supplementary Figure 7

#### Supplementary Figure 7

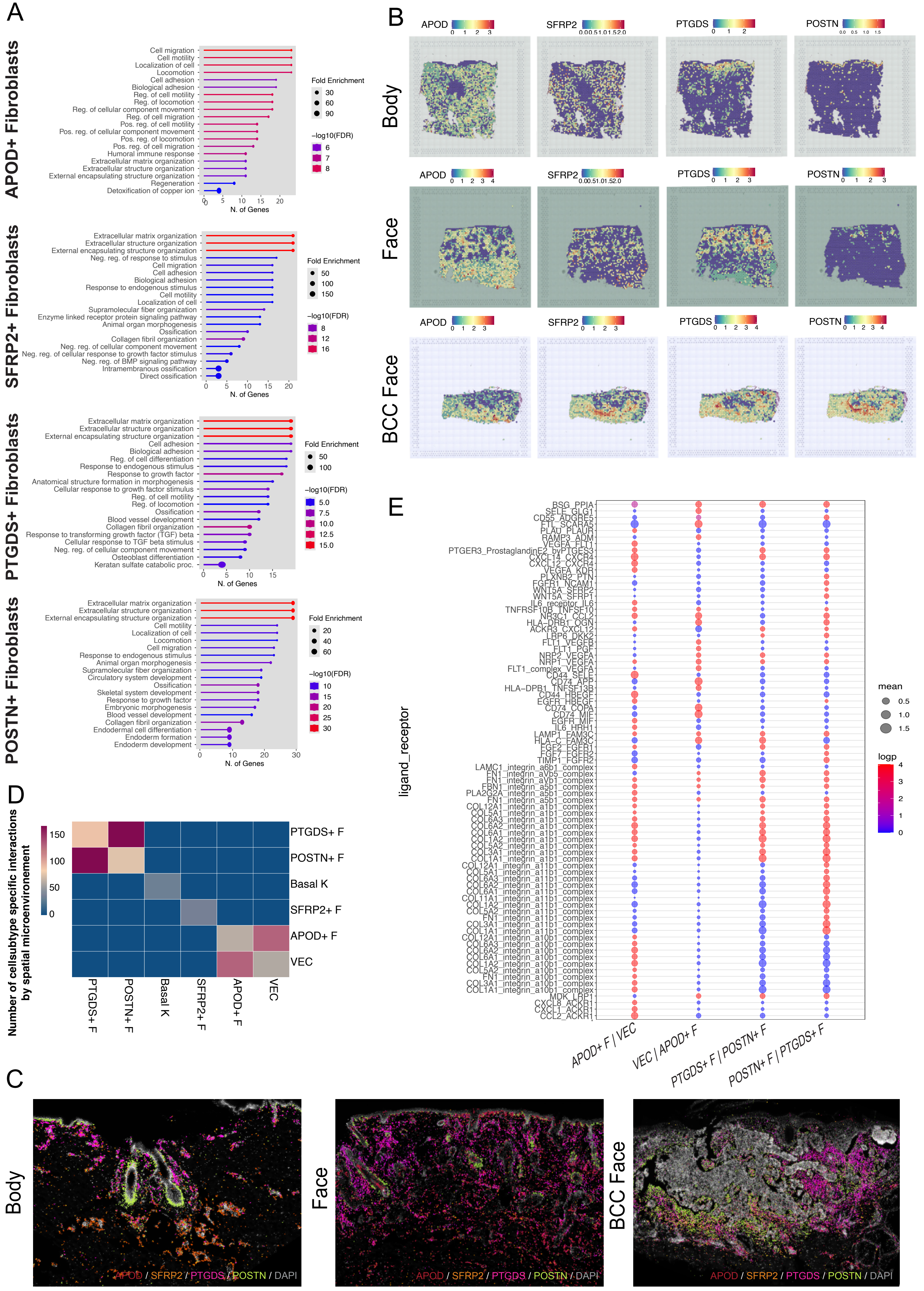

### Supplementary Figure 8

#### Supplementary Figure 8

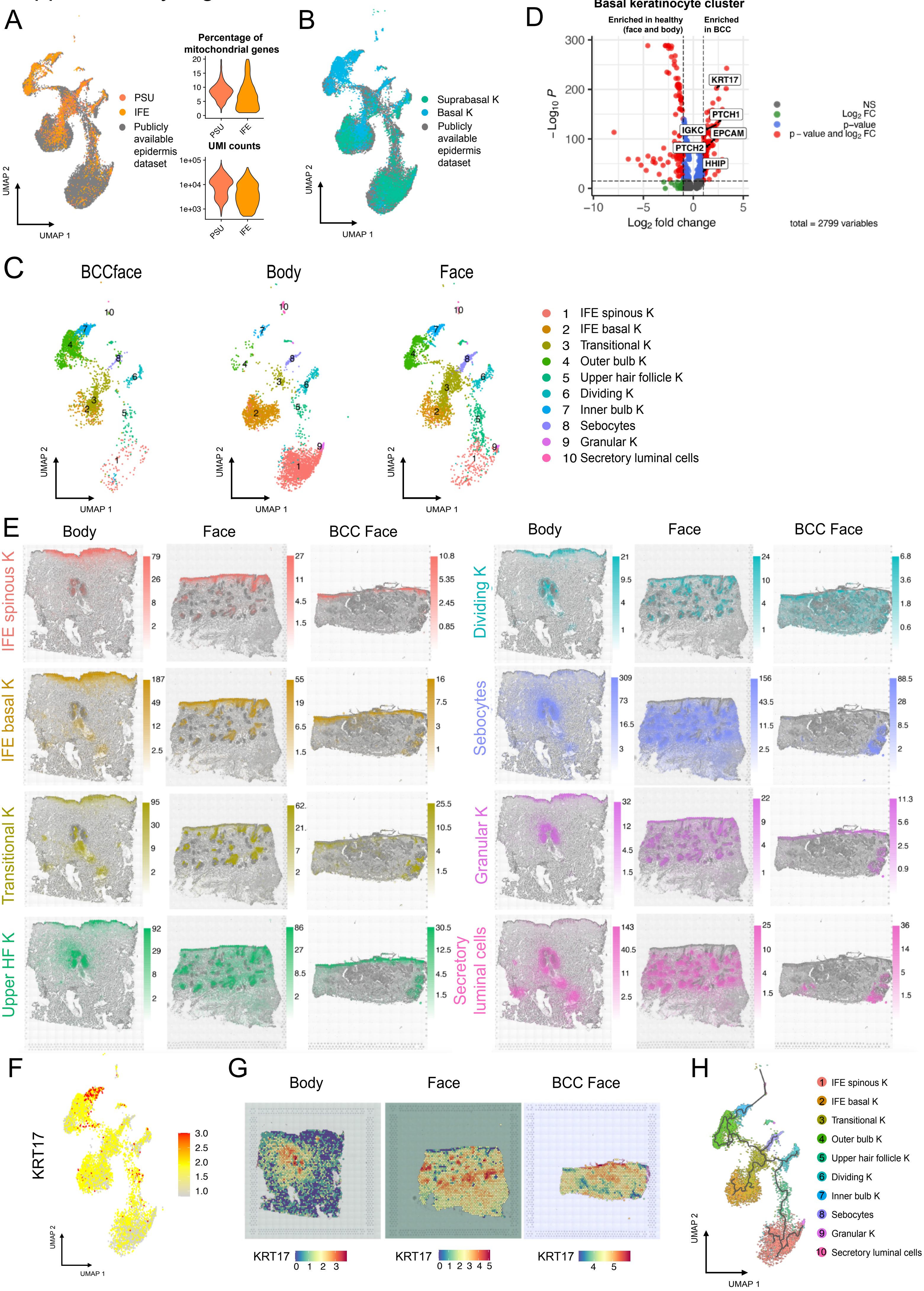
