## Supplementary Figure 2 for "Multi-scale spatial mapping of cell populations across anatomical sites in healthy human skin and basal cell carcinoma"

**A**

| | Name | Disease/ Healthy | Face/Body | Age | Gender | Location | Fitz-patrick scale | N of cells |
|---|---|---|---|---|---|---|---|---|
| Sequenced in this study | face_ear1 | Healthy | Face | 77 | Male | Ear | I/II | 6278 |
| | face_nose1 | Healthy | Face | 56 | Male | Nose | I/II | 6618 |
| | face_cheek1 | Healthy | Face | 85 | Male | Cheek | I/II | 6761 |
| | face_forehead1 | Healthy | Face | 85 | Male | Forehead | I/II | 9499 |
| | face_forehead2 | Healthy | Face | 70 | Male | Forehead | I/II | 1945 |
| | face_cheek2 | Healthy | Face | 62 | Male | Cheek | I/II | 1177 |
| | face_ear2 | Healthy | Face | 86 | Male | Ear | I/II | 3088 |
| | face_temple1 | Healthy | Face | 59 | Male | Temple | I/II | 5105 |
| | face_forehead3 | Healthy | Face | 80 | Male | Forehead | I/II | 8039 |
| | face_temple2 | Healthy | Face | 78 | Male | Temple | I/II | 1344 |
| | face_cheek3b | Healthy | Face | 81 | Female | Cheek | I/II | 4010 |
| | face_forehead4 | Healthy | Face | 90 | Male | Forehead | I/II | 6861 |
| | face_cheek4 | Healthy | Face | 90 | Male | Cheek | I/II | 9250 |
| | face_forehead5 | Healthy | Face | 73 | Male | Forehead | I/II | 5755 |
| | bcc_cheek1 | BCC | Face | 39 | Male | Cheek | I/II | 16303 |
| | bcc_cheek2 | BCC | Face | 85 | Male | Cheek | I/II | 5207 |
| | bcc_ear1 | BCC | Face | 70 | Male | Ear | I/II | 5573 |
| | bcc_ear2 | BCC | Face | 69 | Male | Ear | I/II | 6694 |
| | bcc_forehead1 | BCC | Face | 73 | Male | Forehead | I/II | 10224 |
| | bcc_nose1 | BCC | Face | 91 | Male | Nose | I/II | 6250 |
| | bcc_temple1 | BCC | Face | 51 | Male | Temple | I/II | 6028 |
| | bcc_temple2 | BCC | Face | 59 | Male | Temple | I/II | 2830 |
| Publicly available | body_sole-boldo GSE130973_filtered | Healthy | Body | 25 – 70 | 5 male | Inguinal | n/a | 14033 |
| | body_tabib Control_Human_Skin_6People.csv | Healthy | Body | 23 – 66 | 3 female donors & 3 male donors | Arm | n/a | 6529 |

**B**

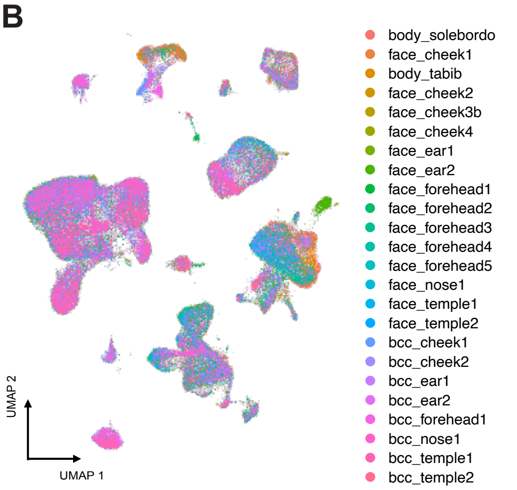


**C**

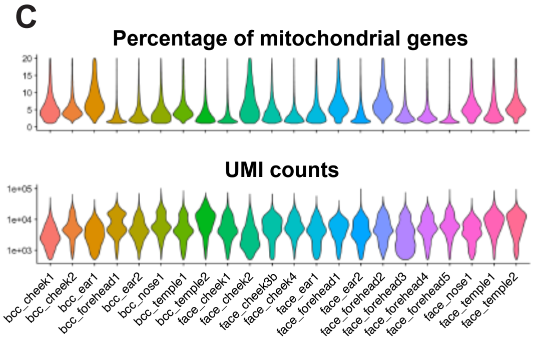


**D**

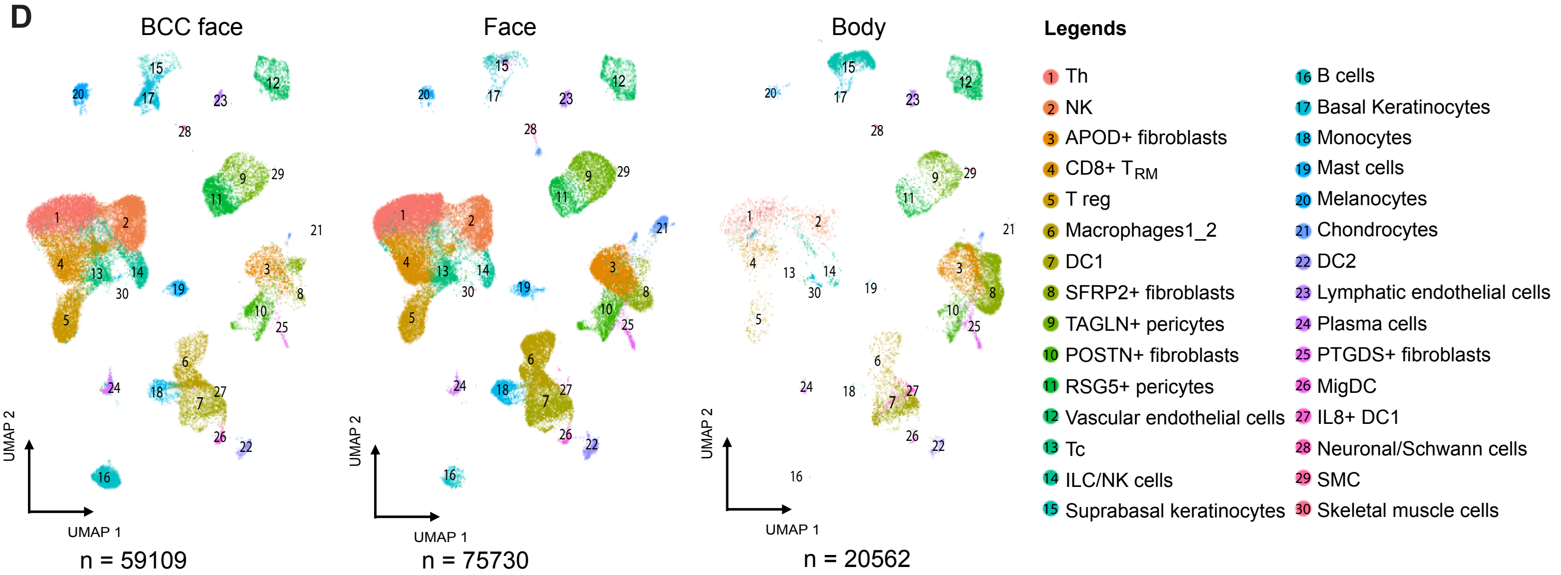


**E**

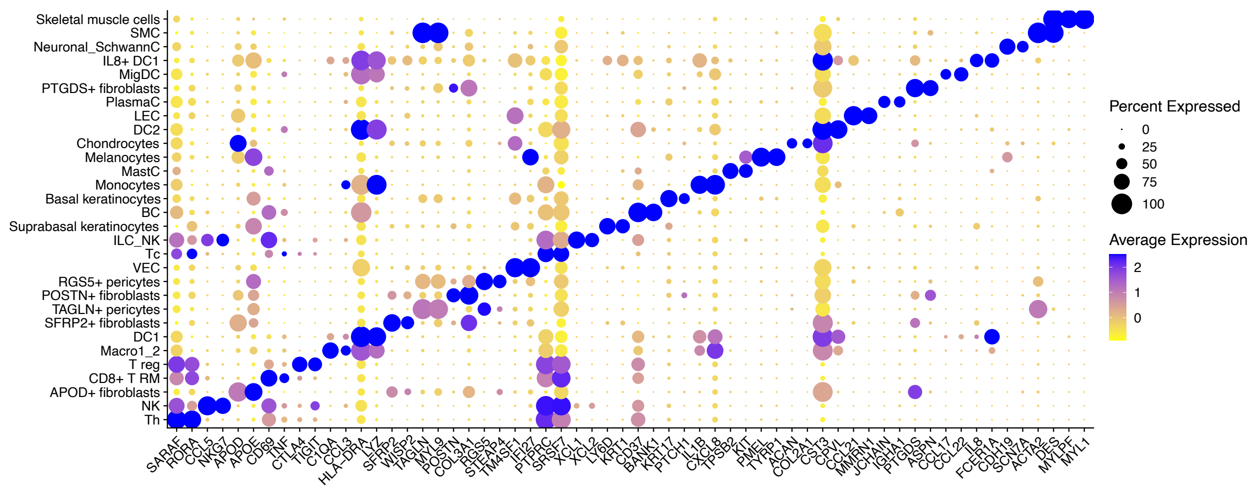


**F**

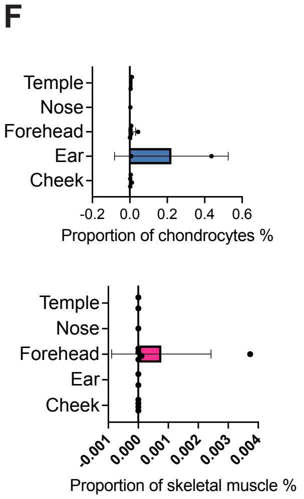
