## Supplementary Figure 4 for "Multi-scale spatial mapping of cell populations across anatomical sites in healthy human skin and basal cell carcinoma"

A 10X visium and *In situ* sequencing samples

| Name | Disease/ Healthy | Face/ Body | Age | Gender | Location | Fitz-patrick scale | N of spots | N of ISS reads |
|---|---|---|---|---|---|---|---|---|
| face_temple1a | Healthy | Face | 68 | Male | Temple | I/II | 986 | |
| face_temple1b | Healthy | Face | 68 | Male | Temple | I/II | 924 | |
| face_temple2a | Healthy | Face | 33 | Female | Temple | I/II | 1520 | |
| face_temple2b | Healthy | Face | 33 | Female | Temple | I/II | 1484 | |
| body_back1a | Healthy | Body | 60 | Male | Back | I/II | 1424 | |
| body_inguinal1a | Healthy | Body | 55 | Male | Inguinal | I/II | 1297 | |
| face_nose1a | Healthy | Face | 77 | Male | Nose | I/II | 1546 | |
| face_glabella1 | Healthy | Face | 68 | Male | Glabella | I/II | 1574 | 947395 |
| face_forehead1a | Healthy | Face | 55 | Male | Forehead | I/II | 1013 | |
| face_cheek1 | Healthy | Face | 39 | Male | Cheek | I/II | 2025 | 1048575 |
| body_abdomen1b | Healthy | Body | 47 | Male | Abdomen | I/II | 1491 | |
| body_thigh1b | Healthy | Body | 53 | Male | Thigh | I/II | 1766 | 310938 |
| body_pubis1 | Healthy | Body | 52 | Male | Pubis | I/II | 1630 | 469219 |
| body_inguinal2 | Healthy | Body | 63 | Male | Inguinal | I/II | 1945 | |
| face_forehead2 | Healthy | Face | 73 | Male | Forehead | I/II | 1957 | 650238 |
| face_forehead1b | Healthy | Face | 55 | Male | Forehead | I/II | 958 | |
| face_nose1b | Healthy | Face | 77 | Male | Nose | I/II | 1639 | 971193 |
| face_cheek2 | Healthy | Face | 76 | Male | Cheek | I/II | 1375 | |
| body_breast1 | Healthy | Body | 42 | Female | Breast | I/II | 2086 | 310513 |
| body_back1b | Healthy | Body | 60 | Male | Back | I/II | 2447 | |
| body_inguinal1b | Healthy | Body | 55 | Male | Inguinal | I/II | 2222 | |
| face_scalp1 | Healthy | Face | 68 | Male | Scalp | I/II | 1342 | |
| bcc_face_cheek1 | BCC | Face | 39 | Male | Cheek | I/II | 811 | 768408 |
| bcc_face_forehead1 | BCC | Face | 73 | Male | Forehead | I/II | 1032 | 1048576 |
| bcc_ face_cheek2 | BCC | Face | 85 | Male | Cheek | I/II | 1868 | 2170139 |
| bcc_ face_nose1 | BCC | Face | 91 | Male | Nose | I/II | 1179 | 3001719 |
| bcc_ face_nose2 | BCC | Face | 72 | Female | Nose | I/II | 1324 | |
| bcc_face_ear1 | BCC | Face | 90 | Male | Ear | I/II | 1123 | |
| bcc_face_forehead2 | BCC | Face | 64 | Female | Forehead | I/II | 907 | |
| bcc_face_nose3 | BCC | Face | 82 | Female | Nose | I/II | 927 | |

B

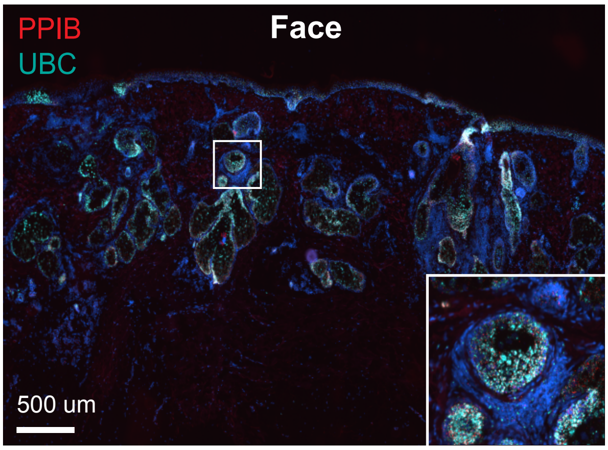


C

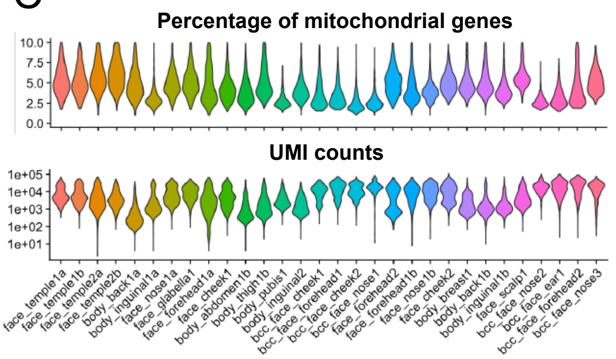


D

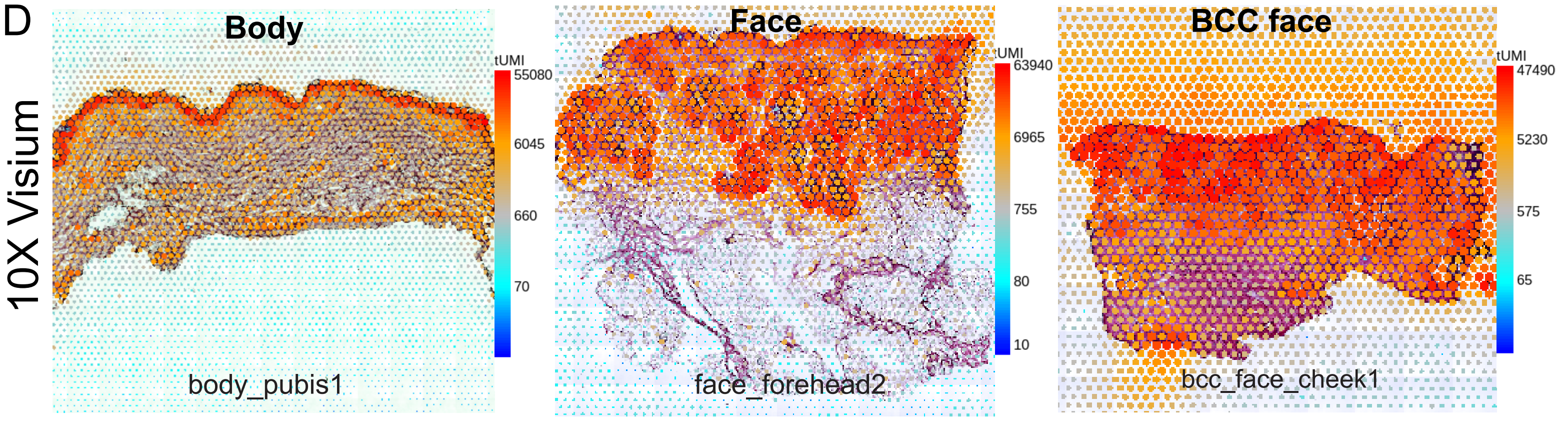


E

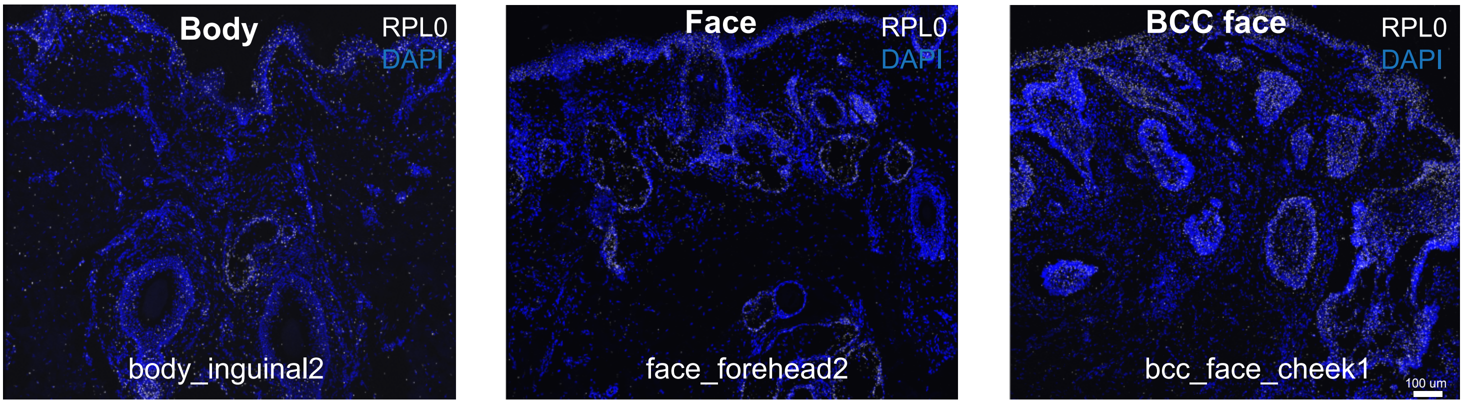


F

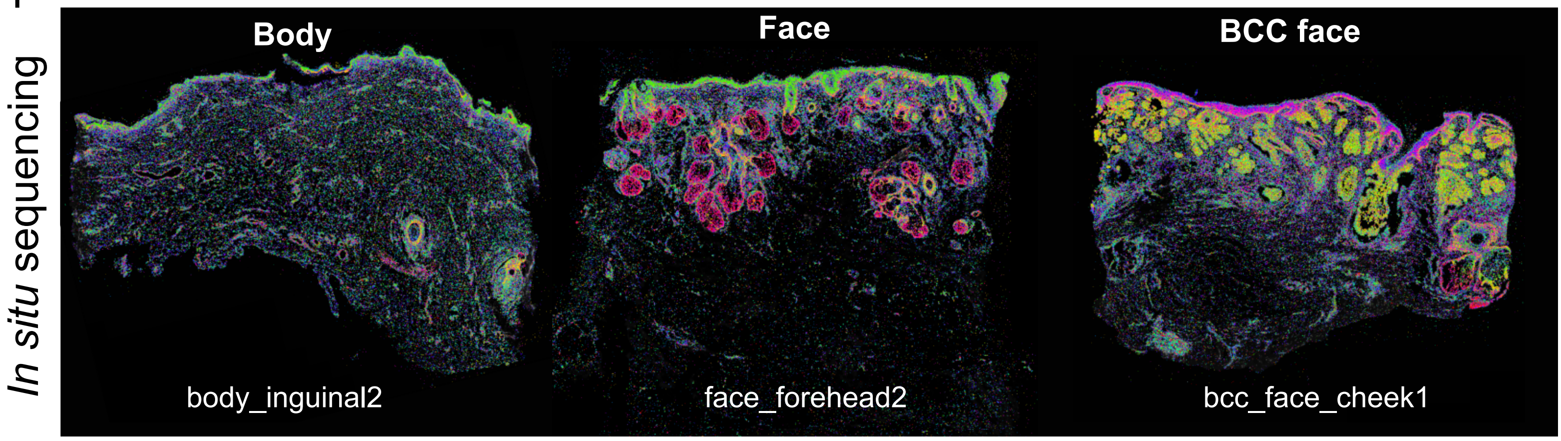
