## Supplementary Table 1 for "Multi-scale spatial mapping of cell populations across anatomical sites in healthy human skin and basal cell carcinoma"

DEG-BCC_and_normal-top20

| | p_val | avg_log2FC | pct.1 | pct.2 | p_val_adj | cluster | gene |
|---|---|---|---|---|---|---|---|
| 1 | 0 | 1.97390835409899 | 0.903 | 0.287 | 0 | Th | IL7R |
| 2 | 0 | 1.78427339536122 | 0.873 | 0.318 | 0 | Th | RORA |
| 3 | 0 | 1.58858723592029 | 0.624 | 0.102 | 0 | Th | TRAT1 |
| 4 | 0 | 1.5805225021889 | 0.918 | 0.484 | 0 | Th | SARAF |
| 5 | 0 | 1.54937084956506 | 0.409 | 0.084 | 0 | Th | KLRB1 |
| 6 | 0 | 1.5472782586616 | 0.804 | 0.205 | 0 | Th | SPOCK2 |
| 7 | 0 | 1.43211040116228 | 0.815 | 0.437 | 0 | Th | NR3C1 |
| 8 | 0 | 1.42382464647342 | 0.937 | 0.694 | 0 | Th | ZFP36L2 |
| 9 | 0 | 1.37676234586136 | 0.974 | 0.506 | 0 | Th | PTPRC |
| 10 | 0 | 1.35835503713398 | 0.97 | 0.622 | 0 | Th | CREM |
| 11 | 0 | 1.35602929586347 | 0.845 | 0.567 | 0 | Th | ANKRD12 |
| 12 | 0 | 1.32925458940398 | 0.866 | 0.407 | 0 | Th | EML4 |
| 13 | 0 | 1.30160326372498 | 0.949 | 0.585 | 0 | Th | EZR |
| 14 | 0 | 1.28612416612016 | 0.784 | 0.311 | 0 | Th | TUBA4A |
| 15 | 0 | 1.26719588332785 | 0.771 | 0.27 | 0 | Th | LEPROTL1 |
| 16 | 0 | 1.26620303547671 | 0.701 | 0.285 | 0 | Th | MBP |
| 17 | 0 | 1.24990951909301 | 0.895 | 0.441 | 0 | Th | ARHGDIB |
| 18 | 0 | 1.21591901599327 | 0.637 | 0.205 | 0 | Th | ALOX5AP |
| 19 | 0 | 1.20783289285518 | 0.429 | 0.086 | 0 | Th | CXCR6 |
| 20 | 0 | 1.18948545440541 | 0.86 | 0.376 | 0 | Th | CNOT6L |
| 21 | 0 | 3.59017509070143 | 0.914 | 0.087 | 0 | NK | CCL5 |
| 22 | 0 | 3.02762660588989 | 0.771 | 0.047 | 0 | NK | NKG7 |
| 23 | 0 | 2.87714324132015 | 0.509 | 0.059 | 0 | NK | IFNG |
| 24 | 0 | 2.68745626632778 | 0.63 | 0.021 | 0 | NK | GZMK |
| 25 | 0 | 2.49775004545892 | 0.533 | 0.082 | 0 | NK | CCL4 |
| 26 | 0 | 2.24026755222372 | 0.776 | 0.19 | 0 | NK | CST7 |
| 27 | 0 | 2.23948303572465 | 0.475 | 0.03 | 0 | NK | CRTAM |
| 28 | 0 | 2.18803866414469 | 0.267 | 0.032 | 0 | NK | CCL4L2 |
| 29 | 0 | 2.00133126351292 | 0.741 | 0.318 | 0 | NK | PIK3R1 |
| 30 | 0 | 1.99096727640418 | 0.357 | 0.027 | 0 | NK | GZMB |
| 31 | 0 | 1.94916236722377 | 0.537 | 0.032 | 0 | NK | CD8A |
| 32 | 0 | 1.83815111737555 | 0.43 | 0.039 | 0 | NK | GZMA |

| | | | | | | | |
|---|---|---|---|---|---|---|---|
| 33 | 0 | 1.77841148555605 | 0.559 | 0.1 | 0 | NK | SH2D1A |
| 34 | 0 | 1.77120461816818 | 0.381 | 0.013 | 0 | NK | GZMH |
| 35 | 0 | 1.7024951647228 | 0.911 | 0.455 | 0 | NK | DUSP2 |
| 36 | 0 | 1.57361356619945 | 0.803 | 0.411 | 0 | NK | CLEC2B |
| 37 | 0 | 1.49580168739309 | 0.778 | 0.364 | 0 | NK | RUNX3 |
| 38 | 0 | 1.46918531159195 | 0.428 | 0.121 | 0 | NK | TNFRSF9 |
| 39 | 0 | 1.46557605559193 | 0.745 | 0.347 | 0 | NK | TUBA4A |
| 40 | 0 | 1.37930959450765 | 0.948 | 0.54 | 0 | NK | PTPRC |
| 41 | 0 | 3.46326653256385 | 0.972 | 0.293 | 0 | APOD+ fibroblasts | CFD |
| 42 | 0 | 3.12279080852043 | 0.931 | 0.181 | 0 | APOD+ fibroblasts | APOD |
| 43 | 0 | 3.10536583546416 | 0.849 | 0.271 | 0 | APOD+ fibroblasts | APOE |
| 44 | 0 | 3.04498968972274 | 0.661 | 0.101 | 0 | APOD+ fibroblasts | PTGDS |
| 45 | 0 | 2.88099768784352 | 0.853 | 0.131 | 0 | APOD+ fibroblasts | CXCL12 |
| 46 | 0 | 2.8634112955035 | 0.946 | 0.177 | 0 | APOD+ fibroblasts | CCDC80 |
| 47 | 0 | 2.80577361896323 | 0.926 | 0.329 | 0 | APOD+ fibroblasts | GSN |
| 48 | 0 | 2.76268416334678 | 0.949 | 0.256 | 0 | APOD+ fibroblasts | MGP |
| 49 | 0 | 2.68339796510441 | 0.869 | 0.124 | 0 | APOD+ fibroblasts | CFH |
| 50 | 0 | 2.64377587388065 | 0.99 | 0.281 | 0 | APOD+ fibroblasts | DCN |
| 51 | 0 | 2.64043942626301 | 0.738 | 0.073 | 0 | APOD+ fibroblasts | FGF7 |
| 52 | 0 | 2.63282028338304 | 0.615 | 0.094 | 0 | APOD+ fibroblasts | TNFAIP6 |
| 53 | 0 | 2.60647605117058 | 0.925 | 0.217 | 0 | APOD+ fibroblasts | CXCL14 |
| 54 | 0 | 2.52757386846696 | 0.786 | 0.114 | 0 | APOD+ fibroblasts | MEG3 |
| 55 | 0 | 2.47764503976724 | 0.911 | 0.629 | 0 | APOD+ fibroblasts | MT2A |
| 56 | 0 | 2.47629879179988 | 0.351 | 0.09 | 0 | APOD+ fibroblasts | CXCL1 |
| 57 | 0 | 2.38902292506913 | 0.877 | 0.202 | 0 | APOD+ fibroblasts | NNMT |
| 58 | 0 | 2.37389178670868 | 0.605 | 0.134 | 0 | APOD+ fibroblasts | MT1M |
| 59 | 0 | 2.28470146526534 | 0.535 | 0.045 | 0 | APOD+ fibroblasts | MEDAG |
| 60 | 0 | 2.21048858636457 | 0.779 | 0.174 | 0 | APOD+ fibroblasts | RND3 |
| 61 | 0 | 2.0710463587906 | 0.798 | 0.311 | 0 | T RM | CD69 |
| 62 | 0 | 1.77820005381196 | 0.434 | 0.126 | 0 | T RM | TNF |
| 63 | 0 | 1.54334508624975 | 0.947 | 0.738 | 0 | T RM | DNAJB1 |
| 64 | 0 | 1.48387861813511 | 0.902 | 0.701 | 0 | T RM | KLF6 |
| 65 | 0 | 1.47027377410847 | 0.674 | 0.344 | 0 | T RM | PTGER4 |
| 66 | 0 | 1.41345622048063 | 0.749 | 0.502 | 0 | T RM | BTG2 |

| | | | | | | | |
|---|---|---|---|---|---|---|---|
| **67** | 0 | 1.38351357457581 | 0.438 | 0.101 | 0 | T RM | CD40LG |
| **68** | 0 | 1.34611437974733 | 0.549 | 0.392 | 0 | T RM | PLIN2 |
| **69** | 0 | 1.32523493342323 | 0.951 | 0.846 | 0 | T RM | HSPA8 |
| **70** | 0 | 1.29962266438229 | 0.373 | 0.114 | 0 | T RM | KLRB1 |
| **71** | 0 | 1.27722662285554 | 0.97 | 0.864 | 0 | T RM | HSPA1A |
| **72** | 0 | 1.27532568812622 | 0.709 | 0.502 | 0 | T RM | DDIT4 |
| **73** | 0 | 1.26541133887579 | 0.915 | 0.686 | 0 | T RM | HSPA1B |
| **74** | 0 | 1.24422050070666 | 0.808 | 0.502 | 0 | T RM | CXCR4 |
| **75** | 0 | 1.2345458578343 | 0.981 | 0.944 | 0 | T RM | UBC |
| **76** | 0 | 1.22295332353711 | 0.727 | 0.373 | 0 | T RM | CD52 |
| **77** | 0 | 1.22076009604185 | 0.987 | 0.957 | 0 | T RM | HSP90AA1 |
| **78** | 0 | 1.21734524096046 | 0.337 | 0.088 | 0 | T RM | TNFSF14 |
| **79** | 0 | 1.19169241642402 | 0.844 | 0.656 | 0 | T RM | SLC2A3 |
| **80** | 0 | 1.1700625386471 | 0.781 | 0.473 | 0 | T RM | DUSP2 |
| **81** | 0 | 2.3928394240666 | 0.722 | 0.071 | 0 | T reg | CTLA4 |
| **82** | 0 | 2.07956020337325 | 0.647 | 0.091 | 0 | T reg | TIGIT |
| **83** | 0 | 2.02346148788152 | 0.837 | 0.391 | 0 | T reg | IL32 |
| **84** | 0 | 1.99891389757546 | 0.657 | 0.192 | 0 | T reg | CARD16 |
| **85** | 0 | 1.96834648975399 | 0.706 | 0.269 | 0 | T reg | PMAIP1 |
| **86** | 0 | 1.93438227394699 | 0.546 | 0.125 | 0 | T reg | TNFRSF4 |
| **87** | 0 | 1.91572855626894 | 0.716 | 0.218 | 0 | T reg | TNFRSF18 |
| **88** | 0 | 1.86797227960874 | 0.517 | 0.053 | 0 | T reg | IKZF2 |
| **89** | 0 | 1.82047511088914 | 0.66 | 0.153 | 0 | T reg | ICOS |
| **90** | 0 | 1.80288891444374 | 0.579 | 0.114 | 0 | T reg | TNFRSF9 |
| **91** | 0 | 1.75717594124963 | 0.563 | 0.076 | 0 | T reg | TBC1D4 |
| **92** | 0 | 1.67781631742143 | 0.649 | 0.173 | 0 | T reg | BATF |
| **93** | 0 | 1.63115268532808 | 0.421 | 0.053 | 0 | T reg | IL2RA |
| **94** | 0 | 1.53226125594729 | 0.818 | 0.382 | 0 | T reg | DUSP4 |
| **95** | 0 | 1.53052230053567 | 0.65 | 0.226 | 0 | T reg | TRBC2 |
| **96** | 0 | 1.51998555449234 | 0.789 | 0.318 | 0 | T reg | CLEC2D |
| **97** | 0 | 1.51779442373243 | 0.66 | 0.203 | 0 | T reg | TRAC |
| **98** | 0 | 1.50853224408966 | 0.517 | 0.181 | 0 | T reg | TRBC1 |
| **99** | 0 | 1.49404759131245 | 0.628 | 0.252 | 0 | T reg | LTB |
| **100** | 0 | 1.49109846091097 | 0.785 | 0.301 | 0 | T reg | CD2 |

| | | | | | | | |
|---|---|---|---|---|---|---|---|
| 101 | 0 | 3.96446107524358 | 0.708 | 0.155 | 0 | Macro1_2 | HMOX1 |
| 102 | 0 | 3.75881997188123 | 0.631 | 0.06 | 0 | Macro1_2 | RNASE1 |
| 103 | 0 | 3.36358367497106 | 0.777 | 0.046 | 0 | Macro1_2 | C1QA |
| 104 | 0 | 3.26651970326482 | 0.793 | 0.252 | 0 | Macro1_2 | CXCL8 |
| 105 | 0 | 3.04676317704746 | 0.447 | 0.075 | 0 | Macro1_2 | CCL3 |
| 106 | 0 | 3.01304703247624 | 0.823 | 0.246 | 0 | Macro1_2 | CTSL |
| 107 | 0 | 2.99278054360039 | 0.908 | 0.226 | 0 | Macro1_2 | CTSB |
| 108 | 0 | 2.94059416501109 | 0.664 | 0.079 | 0 | Macro1_2 | MMP9 |
| 109 | 0 | 2.90882739275502 | 0.758 | 0.19 | 0 | Macro1_2 | CXCL3 |
| 110 | 0 | 2.70665709595213 | 0.838 | 0.074 | 0 | Macro1_2 | FCGR2A |
| 111 | 0 | 2.66915601039518 | 0.998 | 0.96 | 0 | Macro1_2 | FTL |
| 112 | 0 | 2.66187194663018 | 0.802 | 0.052 | 0 | Macro1_2 | C5AR1 |
| 113 | 0 | 2.5865199648891 | 0.379 | 0.073 | 0 | Macro1_2 | EREG |
| 114 | 0 | 2.58027743683602 | 0.85 | 0.105 | 0 | Macro1_2 | AIF1 |
| 115 | 0 | 2.54915378841449 | 0.52 | 0.026 | 0 | Macro1_2 | SDS |
| 116 | 0 | 2.54411560399545 | 0.983 | 0.468 | 0 | Macro1_2 | HLA-DRA |
| 117 | 0 | 2.53567501250212 | 0.633 | 0.028 | 0 | Macro1_2 | C1QB |
| 118 | 0 | 2.52250019139752 | 0.576 | 0.161 | 0 | Macro1_2 | SELENOP |
| 119 | 0 | 2.47184377021033 | 0.652 | 0.024 | 0 | Macro1_2 | C1QC |
| 120 | 0 | 2.44947922401187 | 0.909 | 0.285 | 0 | Macro1_2 | CTSZ |
| 121 | 0 | 3.51381982801513 | 0.997 | 0.474 | 0 | DC1 | HLA-DRA |
| 122 | 0 | 3.41129125961984 | 0.719 | 0.182 | 0 | DC1 | G0S2 |
| 123 | 0 | 3.40308286979823 | 0.989 | 0.392 | 0 | DC1 | HLA-DPB1 |
| 124 | 0 | 3.28057867007514 | 0.976 | 0.229 | 0 | DC1 | HLA-DQA1 |
| 125 | 0 | 3.27975830821445 | 0.988 | 0.383 | 0 | DC1 | HLA-DPA1 |
| 126 | 0 | 3.26634776166907 | 0.99 | 0.432 | 0 | DC1 | HLA-DRB1 |
| 127 | 0 | 3.08558663191892 | 0.91 | 0.16 | 0 | DC1 | LYZ |
| 128 | 0 | 2.93095038912384 | 0.746 | 0.112 | 0 | DC1 | C15orf48 |
| 129 | 0 | 2.92875963713173 | 0.996 | 0.572 | 0 | DC1 | CD74 |
| 130 | 0 | 2.90277589876416 | 0.964 | 0.29 | 0 | DC1 | HLA-DQB1 |
| 131 | 0 | 2.7682291409221 | 0.785 | 0.208 | 0 | DC1 | HLA-DRB5 |
| 132 | 0 | 2.72234743355066 | 0.73 | 0.035 | 0 | DC1 | FCER1A |
| 133 | 0 | 2.44012314808838 | 0.976 | 0.496 | 0 | DC1 | CST3 |
| 134 | 0 | 2.35116337827933 | 0.879 | 0.236 | 0 | DC1 | CD83 |

| | | | | | | | |
|---|---|---|---|---|---|---|---|
| 135 | 0 | 2.23835400230716 | 0.7 | 0.265 | 0 | DC1 | CXCL8 |
| 136 | 0 | 2.21537846770206 | 0.681 | 0.06 | 0 | DC1 | IL1R2 |
| 137 | 0 | 2.21411521738248 | 0.885 | 0.294 | 0 | DC1 | CTSZ |
| 138 | 0 | 2.15272905738606 | 0.922 | 0.146 | 0 | DC1 | TYROBP |
| 139 | 0 | 2.14883606682868 | 0.846 | 0.365 | 0 | DC1 | INSIG1 |
| 140 | 0 | 2.08085914351176 | 0.818 | 0.11 | 0 | DC1 | LST1 |
| 141 | 0 | 4.11726694515631 | 0.837 | 0.087 | 0 | SFRP2+ fibroblasts | SFRP2 |
| 142 | 0 | 3.87451889153788 | 0.991 | 0.295 | 0 | SFRP2+ fibroblasts | DCN |
| 143 | 0 | 3.68021131521306 | 0.869 | 0.127 | 0 | SFRP2+ fibroblasts | FBLN1 |
| 144 | 0 | 3.66493446641538 | 0.927 | 0.231 | 0 | SFRP2+ fibroblasts | CXCL14 |
| 145 | 0 | 3.38214841661242 | 0.935 | 0.309 | 0 | SFRP2+ fibroblasts | CFD |
| 146 | 0 | 3.17075367651847 | 0.861 | 0.118 | 0 | SFRP2+ fibroblasts | MMP2 |
| 147 | 0 | 3.13135321143601 | 0.941 | 0.253 | 0 | SFRP2+ fibroblasts | COL1A2 |
| 148 | 0 | 3.01042203762743 | 0.929 | 0.34 | 0 | SFRP2+ fibroblasts | GSN |
| 149 | 0 | 2.93858030121234 | 0.859 | 0.16 | 0 | SFRP2+ fibroblasts | PLAC9 |
| 150 | 0 | 2.86246337694966 | 0.826 | 0.14 | 0 | SFRP2+ fibroblasts | LUM |
| 151 | 0 | 2.82220812319606 | 0.863 | 0.243 | 0 | SFRP2+ fibroblasts | COL1A1 |
| 152 | 0 | 2.77459195300157 | 0.775 | 0.197 | 0 | SFRP2+ fibroblasts | COL3A1 |
| 153 | 0 | 2.7252916775885 | 0.571 | 0.041 | 0 | SFRP2+ fibroblasts | WISP2 |
| 154 | 0 | 2.70470735350393 | 0.572 | 0.081 | 0 | SFRP2+ fibroblasts | IGFBP6 |
| 155 | 0 | 2.66482267108736 | 0.777 | 0.109 | 0 | SFRP2+ fibroblasts | CTSK |
| 156 | 0 | 2.6633590530971 | 0.742 | 0.104 | 0 | SFRP2+ fibroblasts | MFAP4 |
| 157 | 0 | 2.65883675538538 | 0.944 | 0.243 | 0 | SFRP2+ fibroblasts | COL6A2 |
| 158 | 0 | 2.58951634641883 | 0.333 | 0.029 | 0 | SFRP2+ fibroblasts | SLPI |
| 159 | 0 | 2.57643442723845 | 0.822 | 0.188 | 0 | SFRP2+ fibroblasts | SERPINF1 |
| 160 | 0 | 2.43285429633306 | 0.573 | 0.052 | 0 | SFRP2+ fibroblasts | SEPP1 |
| 161 | 0 | 4.43656894906895 | 0.972 | 0.117 | 0 | TAGLN+ pericytes | TAGLN |
| 162 | 0 | 4.27077942773467 | 0.882 | 0.071 | 0 | TAGLN+ pericytes | ACTA2 |
| 163 | 0 | 3.96088452617759 | 0.954 | 0.127 | 0 | TAGLN+ pericytes | TPM2 |
| 164 | 0 | 3.84280185093923 | 0.945 | 0.131 | 0 | TAGLN+ pericytes | MYL9 |
| 165 | 0 | 3.41310348685049 | 0.945 | 0.255 | 0 | TAGLN+ pericytes | C11orf96 |
| 166 | 0 | 3.30958282820359 | 0.978 | 0.257 | 0 | TAGLN+ pericytes | ADIRF |
| 167 | 0 | 3.2107971518011 | 0.819 | 0.023 | 0 | TAGLN+ pericytes | MYH11 |
| 168 | 0 | 3.01037772752229 | 0.937 | 0.172 | 0 | TAGLN+ pericytes | SPARCL1 |

| | | | | | | | |
|---|---|---|---|---|---|---|---|
| 169 | 0 | 2.90404680717784 | 0.844 | 0.162 | 0 | TAGLN+ pericytes | ADAMTS1 |
| 170 | 0 | 2.8709683337411 | 0.62 | 0.059 | 0 | TAGLN+ pericytes | RGS5 |
| 171 | 0 | 2.79410799353959 | 0.615 | 0.111 | 0 | TAGLN+ pericytes | MT1A |
| 172 | 0 | 2.72986179918426 | 0.788 | 0.131 | 0 | TAGLN+ pericytes | CRISPLD2 |
| 173 | 0 | 2.67107573078146 | 0.513 | 0.12 | 0 | TAGLN+ pericytes | IL6 |
| 174 | 0 | 2.6440177061596 | 0.952 | 0.448 | 0 | TAGLN+ pericytes | DSTN |
| 175 | 0 | 2.57011885056351 | 0.573 | 0.024 | 0 | TAGLN+ pericytes | AVPR1A |
| 176 | 0 | 2.54978815878258 | 0.848 | 0.113 | 0 | TAGLN+ pericytes | MCAM |
| 177 | 0 | 2.49887419981743 | 0.741 | 0.24 | 0 | TAGLN+ pericytes | CCL2 |
| 178 | 0 | 2.48586858421709 | 0.481 | 0.003 | 0 | TAGLN+ pericytes | RERGL |
| 179 | 0 | 2.47149368373905 | 0.982 | 0.324 | 0 | TAGLN+ pericytes | IGFBP7 |
| 180 | 0 | 2.45520949523911 | 0.969 | 0.285 | 0 | TAGLN+ pericytes | CALD1 |
| 181 | 0 | 4.4436121906627 | 0.935 | 0.251 | 0 | POSTN+ fibroblasts | COL1A1 |
| 182 | 0 | 3.89630155252647 | 0.899 | 0.203 | 0 | POSTN+ fibroblasts | COL3A1 |
| 183 | 0 | 3.72659731013533 | 0.961 | 0.264 | 0 | POSTN+ fibroblasts | COL1A2 |
| 184 | 0 | 3.41060282189462 | 0.657 | 0.054 | 0 | POSTN+ fibroblasts | POSTN |
| 185 | 0 | 2.89247808837419 | 0.919 | 0.217 | 0 | POSTN+ fibroblasts | COL6A1 |
| 186 | 0 | 2.78908074104753 | 0.841 | 0.257 | 0 | POSTN+ fibroblasts | SPARC |
| 187 | 0 | 2.63729085630616 | 0.939 | 0.255 | 0 | POSTN+ fibroblasts | COL6A2 |
| 188 | 0 | 2.59406613067044 | 0.786 | 0.132 | 0 | POSTN+ fibroblasts | COL6A3 |
| 189 | 0 | 2.46111912373621 | 0.826 | 0.151 | 0 | POSTN+ fibroblasts | LUM |
| 190 | 0 | 2.38104670135235 | 0.823 | 0.137 | 0 | POSTN+ fibroblasts | MEG3 |
| 191 | 0 | 2.19164105069816 | 0.642 | 0.08 | 0 | POSTN+ fibroblasts | COL5A2 |
| 192 | 0 | 2.12228523036517 | 0.615 | 0.153 | 0 | POSTN+ fibroblasts | CTGF |
| 193 | 0 | 2.10053633943176 | 0.682 | 0.12 | 0 | POSTN+ fibroblasts | HTRA1 |
| 194 | 0 | 2.09345001456348 | 0.499 | 0.031 | 0 | POSTN+ fibroblasts | ASPN |
| 195 | 0 | 2.0659487727879 | 0.565 | 0.051 | 0 | POSTN+ fibroblasts | MFAP5 |
| 196 | 0 | 1.87253552087119 | 0.396 | 0.015 | 0 | POSTN+ fibroblasts | COL11A1 |
| 197 | 0 | 1.87074087018723 | 0.505 | 0.071 | 0 | POSTN+ fibroblasts | DIO2 |
| 198 | 0 | 1.84865153679926 | 0.479 | 0.055 | 0 | POSTN+ fibroblasts | COL5A1 |
| 199 | 0 | 1.84425795859932 | 0.817 | 0.21 | 0 | POSTN+ fibroblasts | CCDC80 |
| 200 | 0 | 1.828934266552 | 0.691 | 0.142 | 0 | POSTN+ fibroblasts | TWIST1 |
| 201 | 0 | 3.95860317842631 | 0.824 | 0.053 | 0 | RGS5+ pericytes | RGS5 |
| 202 | 0 | 3.2556360039452 | 0.802 | 0.211 | 0 | RGS5+ pericytes | RGS16 |

| | | | | | | | |
|---|---|---|---|---|---|---|---|
| 203 | 0 | 3.1135443021037 | 0.785 | 0.118 | 0 | RGS5+ pericytes | ID4 |
| 204 | 0 | 2.86470296126541 | 0.538 | 0.052 | 0 | RGS5+ pericytes | TCIM |
| 205 | 0 | 2.84857712681214 | 0.825 | 0.103 | 0 | RGS5+ pericytes | NR2F2 |
| 206 | 0 | 2.81686920271023 | 0.329 | 0.051 | 0 | RGS5+ pericytes | CCL19 |
| 207 | 0 | 2.60326150374859 | 0.627 | 0.044 | 0 | RGS5+ pericytes | STEAP4 |
| 208 | 0 | 2.57671187929255 | 0.678 | 0.244 | 0 | RGS5+ pericytes | CCL2 |
| 209 | 0 | 2.5013871171113 | 0.955 | 0.288 | 0 | RGS5+ pericytes | CALD1 |
| 210 | 0 | 2.49273745810784 | 0.584 | 0.062 | 0 | RGS5+ pericytes | KCNE4 |
| 211 | 0 | 2.43199120249369 | 0.73 | 0.136 | 0 | RGS5+ pericytes | CRISPLD2 |
| 212 | 0 | 2.4205324896934 | 0.795 | 0.119 | 0 | RGS5+ pericytes | PDGFRB |
| 213 | 0 | 2.32924046547758 | 0.554 | 0.051 | 0 | RGS5+ pericytes | NDUFA4L2 |
| 214 | 0 | 2.22961326059577 | 0.577 | 0.117 | 0 | RGS5+ pericytes | ADAMTS4 |
| 215 | 0 | 2.19836760832914 | 0.65 | 0.049 | 0 | RGS5+ pericytes | ANGPT2 |
| 216 | 0 | 2.17500327954868 | 0.924 | 0.329 | 0 | RGS5+ pericytes | IGFBP7 |
| 217 | 0 | 2.16961319769152 | 0.713 | 0.17 | 0 | RGS5+ pericytes | ADAMTS1 |
| 218 | 0 | 2.14256227527685 | 0.493 | 0.035 | 0 | RGS5+ pericytes | GJA4 |
| 219 | 0 | 2.10471080885966 | 0.856 | 0.261 | 0 | RGS5+ pericytes | C11orf96 |
| 220 | 0 | 2.095435153084 | 0.808 | 0.163 | 0 | RGS5+ pericytes | PRRX1 |
| 221 | 0 | 4.62085162708587 | 0.925 | 0.108 | 0 | VEC | TM4SF1 |
| 222 | 0 | 4.39011775581363 | 0.463 | 0.02 | 0 | VEC | SELE |
| 223 | 0 | 3.52399915357541 | 0.921 | 0.163 | 0 | VEC | IFI27 |
| 224 | 0 | 3.16584241332455 | 0.361 | 0.015 | 0 | VEC | STC1 |
| 225 | 0 | 3.09996075683418 | 0.77 | 0.053 | 0 | VEC | AQP1 |
| 226 | 0 | 2.82425711548053 | 0.843 | 0.258 | 0 | VEC | TSC22D1 |
| 227 | 0 | 2.80935447435569 | 0.786 | 0.098 | 0 | VEC | GNG11 |
| 228 | 0 | 2.78981216775848 | 0.27 | 0.01 | 0 | VEC | DARC |
| 229 | 0 | 2.73898449186426 | 0.689 | 0.134 | 0 | VEC | SPRY1 |
| 230 | 0 | 2.69004615924103 | 0.605 | 0.016 | 0 | VEC | CLDN5 |
| 231 | 0 | 2.53904623214288 | 0.454 | 0.015 | 0 | VEC | C2CD4B |
| 232 | 0 | 2.49587533605536 | 0.254 | 0.004 | 0 | VEC | ACKR1 |
| 233 | 0 | 2.46094974835291 | 0.893 | 0.18 | 0 | VEC | SPARCL1 |
| 234 | 0 | 2.45363871274056 | 0.613 | 0.007 | 0 | VEC | PLVAP |
| 235 | 0 | 2.40596473072647 | 0.64 | 0.073 | 0 | VEC | MCTP1 |
| 236 | 0 | 2.38569801839529 | 0.337 | 0.011 | 0 | VEC | CSF3 |

| | | | | | | | |
|---|---|---|---|---|---|---|---|
| 237 | 0 | 2.37982791346583 | 0.422 | 0.01 | 0 | VEC | RBP7 |
| 238 | 0 | 2.37226863617029 | 0.558 | 0.072 | 0 | VEC | RCAN1 |
| 239 | 0 | 2.35785084472881 | 0.571 | 0.048 | 0 | VEC | ADAMTS9 |
| 240 | 0 | 2.27574269268504 | 0.557 | 0.044 | 0 | VEC | CALCRL |
| 241 | 0 | 1.42981892533727 | 0.999 | 0.999 | 0 | Tc | MALAT1 |
| 242 | 0 | 1.33048269754277 | 0.789 | 0.57 | 0 | Tc | PTPRC |
| 243 | 0 | 1.32603138015463 | 0.755 | 0.734 | 0 | Tc | SRSF7 |
| 244 | 0 | 1.32128444509003 | 0.768 | 0.76 | 0 | Tc | MTRNR2L1 |
| 245 | 1.6668 | 1.26528955757821 | 0.666 | 0.574 | 5.5000816 | Tc | STK4 |
| 246 | 1.9011 | 1.3695737430568 | 0.543 | 0.474 | 6.2730021 | Tc | EML4 |
| 247 | 1.4454 | 1.48062806744763 | 0.508 | 0.471 | 4.7693414 | Tc | SMCHD1 |
| 248 | 2.0690 | 1.2500829241857 | 0.526 | 0.533 | 6.8269916 | Tc | CHD2 |
| 249 | 2.8898 | 1.40602373476933 | 0.422 | 0.361 | 9.5352056 | Tc | RNF213 |
| 250 | 2.1927 | 1.23402158985789 | 0.377 | 0.295 | 7.2353222 | Tc | IKZF1 |
| 251 | 5.3673 | 1.22193773897327 | 0.367 | 0.285 | 1.7710229 | Tc | ACAP1 |
| 252 | 9.9988 | 1.2149653823264 | 0.469 | 0.399 | 3.2992087 | Tc | RORA |
| 253 | 7.3053 | 1.29136598348125 | 0.408 | 0.344 | 2.4104794 | Tc | SYNE2 |
| 254 | 9.2532 | 1.23822919414785 | 0.435 | 0.414 | 3.0532033 | Tc | NKTR |
| 255 | 2.0730 | 1.33112679332914 | 0.553 | 0.61 | 6.8402060 | Tc | ANKRD12 |
| 256 | 7.8366 | 1.26614338642381 | 0.413 | 0.375 | 2.5857911 | Tc | AC058791.1 |
| 257 | 2.6911 | 1.26914154375853 | 0.465 | 0.466 | 8.8798207 | Tc | TSPYL2 |
| 258 | 2.7030 | 1.23424265592246 | 0.315 | 0.3 | 8.9189381 | Tc | SLC5A3 |
| 259 | 3.2754 | 1.52764706018505 | 0.352 | 0.362 | 1.0807600 | Tc | PLCG2 |
| 260 | 8.7701 | 1.22546070999504 | 0.297 | 0.293 | 2.8938126 | Tc | ANKRD28 |
| 261 | 0 | 4.76788797356367 | 0.813 | 0.039 | 0 | ILC_NK | XCL1 |
| 262 | 0 | 4.42259940350476 | 0.651 | 0.038 | 0 | ILC_NK | GNLY |
| 263 | 0 | 4.34564839709996 | 0.664 | 0.032 | 0 | ILC_NK | XCL2 |
| 264 | 0 | 2.43467528365319 | 0.555 | 0.022 | 0 | ILC_NK | KLRD1 |
| 265 | 0 | 2.314287294876 | 0.401 | 0.046 | 0 | ILC_NK | GZMB |
| 266 | 0 | 2.05188206485461 | 0.583 | 0.098 | 0 | ILC_NK | NKG7 |
| 267 | 0 | 1.99778309492716 | 0.573 | 0.062 | 0 | ILC_NK | CTSW |
| 268 | 0 | 1.96967188337752 | 0.401 | 0.006 | 0 | ILC_NK | KLRC1 |
| 269 | 0 | 1.75288835015594 | 0.525 | 0.17 | 0 | ILC_NK | AREG |
| 270 | 0 | 1.64441419028777 | 0.501 | 0.122 | 0 | ILC_NK | KLRB1 |

| | | | | | | | |
|---|---|---|---|---|---|---|---|
| 271 | 0 | 1.58970098045589 | 0.341 | 0.062 | 0 | ILC_NK | CRTAM |
| 272 | 0 | 1.57397736529042 | 0.574 | 0.148 | 0 | ILC_NK | CCL5 |
| 273 | 0 | 1.55407093485227 | 0.621 | 0.156 | 0 | ILC_NK | CD7 |
| 274 | 0 | 1.54598069958049 | 0.942 | 0.704 | 0 | ILC_NK | REL |
| 275 | 0 | 1.4753321181558 | 0.692 | 0.243 | 0 | ILC_NK | TNFRSF18 |
| 276 | 0 | 1.39377717206013 | 0.684 | 0.346 | 0 | ILC_NK | PIK3R1 |
| 277 | 0 | 1.39110063488152 | 0.652 | 0.245 | 0 | ILC_NK | SYTL3 |
| 278 | 0 | 1.37856893364807 | 0.799 | 0.436 | 0 | ILC_NK | CLEC2B |
| 279 | 0 | 1.37160871843447 | 0.579 | 0.218 | 0 | ILC_NK | ADGRE5 |
| 280 | 0 | 1.34475891050692 | 0.781 | 0.334 | 0 | ILC_NK | CD69 |
| 281 | 0 | 5.58097669518395 | 0.932 | 0.208 | 0 | Suprabasal keratinc | KRT14 |
| 282 | 0 | 5.24818832326327 | 0.776 | 0.046 | 0 | Suprabasal keratinc | DMKN |
| 283 | 0 | 5.2041147068356 | 0.971 | 0.098 | 0 | Suprabasal keratinc | SFN |
| 284 | 0 | 4.99447654884469 | 0.666 | 0.037 | 0 | Suprabasal keratinc | KRT1 |
| 285 | 0 | 4.83420579869808 | 0.829 | 0.285 | 0 | Suprabasal keratinc | KRT10 |
| 286 | 0 | 4.50959869997311 | 0.828 | 0.201 | 0 | Suprabasal keratinc | S100A2 |
| 287 | 0 | 4.38228446319702 | 0.969 | 0.152 | 0 | Suprabasal keratinc | PERP |
| 288 | 0 | 4.29128074936503 | 0.552 | 0.018 | 0 | Suprabasal keratinc | KRTDAP |
| 289 | 0 | 4.05867072153456 | 0.779 | 0.024 | 0 | Suprabasal keratinc | LY6D |
| 290 | 0 | 3.98490188238559 | 0.937 | 0.06 | 0 | Suprabasal keratinc | S100A14 |
| 291 | 0 | 3.9843726690773 | 0.871 | 0.031 | 0 | Suprabasal keratinc | LGALS7B |
| 292 | 0 | 3.79796644801845 | 0.82 | 0.117 | 0 | Suprabasal keratinc | KRT5 |
| 293 | 0 | 3.6313619915386 | 0.892 | 0.047 | 0 | Suprabasal keratinc | TACSTD2 |
| 294 | 0 | 3.42796666405753 | 0.766 | 0.012 | 0 | Suprabasal keratinc | MIR205HG |
| 295 | 0 | 3.24484602854855 | 0.89 | 0.123 | 0 | Suprabasal keratinc | AQP3 |
| 296 | 0 | 3.12725530407241 | 0.71 | 0.069 | 0 | Suprabasal keratinc | LYPD3 |
| 297 | 0 | 2.82923134584601 | 0.828 | 0.108 | 0 | Suprabasal keratinc | GNB2L1 |
| 298 | 0 | 2.76980225550851 | 0.794 | 0.021 | 0 | Suprabasal keratinc | SERPINB5 |
| 299 | 0 | 2.74616725238925 | 0.854 | 0.063 | 0 | Suprabasal keratinc | DSP |
| 300 | 0 | 2.67089898157187 | 0.497 | 0.026 | 0 | Suprabasal keratinc | KRT16 |
| 301 | 0 | 4.07370185859762 | 0.971 | 0.009 | 0 | BC | MS4A1 |
| 302 | 0 | 2.81321820058107 | 0.836 | 0.008 | 0 | BC | BANK1 |
| 303 | 0 | 2.34098701740075 | 0.939 | 0.361 | 0 | BC | CD37 |
| 304 | 0 | 2.33174947890381 | 0.672 | 0.093 | 0 | BC | LY9 |

| | | | | | | | |
|---|---|---|---|---|---|---|---|
| 305 | 0 | 2.07345726851767 | 0.677 | 0.011 | 0 | BC | CD79A |
| 306 | 0 | 2.00135878874789 | 0.778 | 0.156 | 0 | BC | MEF2C |
| 307 | 0 | 1.82270270461117 | 0.684 | 0.121 | 0 | BC | SWAP70 |
| 308 | 0 | 1.82071151557367 | 0.493 | 0.096 | 0 | BC | CCR7 |
| 309 | 0 | 1.77737232413247 | 0.602 | 0.044 | 0 | BC | RALGPS2 |
| 310 | 0 | 1.74560390282892 | 0.529 | 0.009 | 0 | BC | TNFRSF13B |
| 311 | 0 | 1.68432140810676 | 0.272 | 0.007 | 0 | BC | IGHM |
| 312 | 0 | 1.6452222931062 | 0.519 | 0.006 | 0 | BC | LINC00926 |
| 313 | 0 | 1.60985017512494 | 0.818 | 0.253 | 0 | BC | CD83 |
| 314 | 0 | 1.59039397378308 | 0.701 | 0.198 | 0 | BC | PARP14 |
| 315 | 0 | 1.58925005908667 | 0.418 | 0.051 | 0 | BC | XIST |
| 316 | 0 | 1.58054834746853 | 0.536 | 0.01 | 0 | BC | TNFRSF13C |
| 317 | 0 | 1.56719300342792 | 0.814 | 0.265 | 0 | BC | LTB |
| 318 | 0 | 1.50114971571299 | 0.923 | 0.249 | 0 | BC | HLA-DQA1 |
| 319 | 0 | 1.45647176592788 | 0.875 | 0.386 | 0 | BC | CD52 |
| 320 | 0 | 1.44262931128644 | 0.488 | 0.049 | 0 | BC | BCL11A |
| 321 | 0 | 4.99118795194765 | 0.821 | 0.119 | 0 | Basal keratinocytes | KRT17 |
| 322 | 0 | 3.73005716926308 | 0.882 | 0.118 | 0 | Basal keratinocytes | KRT5 |
| 323 | 0 | 3.70760248015657 | 0.788 | 0.068 | 0 | Basal keratinocytes | DSP |
| 324 | 0 | 3.51280923987983 | 0.729 | 0.046 | 0 | Basal keratinocytes | KRT15 |
| 325 | 0 | 3.06078137713857 | 0.786 | 0.211 | 0 | Basal keratinocytes | DST |
| 326 | 0 | 2.96370361497919 | 0.921 | 0.211 | 0 | Basal keratinocytes | KRT14 |
| 327 | 0 | 2.71217811707978 | 0.613 | 0.075 | 0 | Basal keratinocytes | SPON2 |
| 328 | 0 | 2.7016251436436 | 0.805 | 0.315 | 0 | Basal keratinocytes | SOX4 |
| 329 | 0 | 2.64259691449427 | 0.57 | 0.036 | 0 | Basal keratinocytes | DAPL1 |
| 330 | 0 | 2.61666353212792 | 0.541 | 0.032 | 0 | Basal keratinocytes | EDIL3 |
| 331 | 0 | 2.53484113617689 | 0.603 | 0.011 | 0 | Basal keratinocytes | GJB6 |
| 332 | 0 | 2.49357265267839 | 0.677 | 0.07 | 0 | Basal keratinocytes | IRX2 |
| 333 | 0 | 2.45385801934551 | 0.749 | 0.206 | 0 | Basal keratinocytes | S100A2 |
| 334 | 0 | 2.28328998086465 | 0.618 | 0.046 | 0 | Basal keratinocytes | CLDN1 |
| 335 | 0 | 2.26473327329529 | 0.556 | 0.012 | 0 | Basal keratinocytes | TFAP2B |
| 336 | 0 | 2.2414993571333 | 0.841 | 0.158 | 0 | Basal keratinocytes | PERP |
| 337 | 0 | 2.21392974051517 | 0.491 | 0.028 | 0 | Basal keratinocytes | PTCH1 |
| 338 | 0 | 2.2027161343959 | 0.566 | 0.02 | 0 | Basal keratinocytes | BNC2 |

| | | | | | | | |
|---|---|---|---|---|---|---|---|
| 339 | 0 | 2.10988571016691 | 0.677 | 0.146 | 0 | Basal keratinocytes | NFIB |
| 340 | 0 | 2.04416150773796 | 0.533 | 0.084 | 0 | Basal keratinocytes | ABI3BP |
| 341 | 0 | 5.16885528980248 | 0.526 | 0.046 | 0 | Monocytes | SERPINB2 |
| 342 | 0 | 4.77504233409099 | 0.869 | 0.122 | 0 | Monocytes | IL1B |
| 343 | 0 | 4.56773061079668 | 0.911 | 0.076 | 0 | Monocytes | EREG |
| 344 | 0 | 4.29349108588412 | 0.773 | 0.225 | 0 | Monocytes | THBS1 |
| 345 | 0 | 4.12801256418339 | 0.949 | 0.273 | 0 | Monocytes | CXCL8 |
| 346 | 0 | 4.0957355450031 | 0.864 | 0.094 | 0 | Monocytes | S100A9 |
| 347 | 0 | 3.41238029005307 | 0.783 | 0.078 | 0 | Monocytes | S100A8 |
| 348 | 0 | 3.33277382658996 | 0.968 | 0.488 | 0 | Monocytes | TIMP1 |
| 349 | 0 | 3.11503351918304 | 0.815 | 0.013 | 0 | Monocytes | AQP9 |
| 350 | 0 | 3.06632425799644 | 0.917 | 0.188 | 0 | Monocytes | BCL2A1 |
| 351 | 0 | 3.06335901985058 | 0.788 | 0.214 | 0 | Monocytes | CXCL3 |
| 352 | 0 | 3.00804258684543 | 0.474 | 0.09 | 0 | Monocytes | CCL20 |
| 353 | 0 | 3.00118268020535 | 0.956 | 0.183 | 0 | Monocytes | LYZ |
| 354 | 0 | 2.99538387753592 | 0.983 | 0.558 | 0 | Monocytes | SOD2 |
| 355 | 0 | 2.94660834842722 | 0.658 | 0.13 | 0 | Monocytes | PTGS2 |
| 356 | 0 | 2.81535794048853 | 0.635 | 0.093 | 0 | Monocytes | IL1RN |
| 357 | 0 | 2.58404507733825 | 0.907 | 0.247 | 0 | Monocytes | CTSS |
| 358 | 0 | 2.57581265193771 | 0.898 | 0.296 | 0 | Monocytes | CXCL2 |
| 359 | 0 | 2.50103072952671 | 0.802 | 0.217 | 0 | Monocytes | BASP1 |
| 360 | 0 | 2.47085783761669 | 0.803 | 0.195 | 0 | Monocytes | PPIF |
| 361 | 0 | 6.08750091431936 | 0.919 | 0.035 | 0 | MastC | TPSAB1 |
| 362 | 0 | 5.26614976944717 | 0.755 | 0.032 | 0 | MastC | TPSB2 |
| 363 | 0 | 4.73972855670069 | 0.823 | 0.028 | 0 | MastC | GATA2 |
| 364 | 0 | 4.65189792635313 | 0.767 | 0.011 | 0 | MastC | CPA3 |
| 365 | 0 | 4.57823593531994 | 0.789 | 0.063 | 0 | MastC | HPGD |
| 366 | 0 | 4.54427223756633 | 0.646 | 0.017 | 0 | MastC | CTSG |
| 367 | 0 | 4.22839070617712 | 0.511 | 0.007 | 0 | MastC | ADCYAP1 |
| 368 | 0 | 3.81435436234703 | 0.696 | 0.008 | 0 | MastC | HDC |
| 369 | 0 | 3.60564555925552 | 0.807 | 0.384 | 0 | MastC | GLUL |
| 370 | 0 | 3.46683163298305 | 0.641 | 0.023 | 0 | MastC | KIT |
| 371 | 0 | 3.17551725206878 | 0.521 | 0.008 | 0 | MastC | HPGDS |
| 372 | 0 | 3.14782065574649 | 0.615 | 0.027 | 0 | MastC | IL1RL1 |

| | | | | | | | |
|---|---|---|---|---|---|---|---|
| 373 | 0 | 3.14737145239979 | 0.843 | 0.437 | 0 | MastC | RGS2 |
| 374 | 0 | 2.84955227705726 | 0.663 | 0.288 | 0 | MastC | SGK1 |
| 375 | 0 | 2.78720637864792 | 0.441 | 0.06 | 0 | MastC | LTC4S |
| 376 | 0 | 2.69157323070548 | 0.441 | 0.003 | 0 | MastC | GCSAML |
| 377 | 0 | 2.64564183006913 | 0.447 | 0.008 | 0 | MastC | SLC18A2 |
| 378 | 0 | 2.60936572575186 | 0.715 | 0.501 | 0 | MastC | FOSB |
| 379 | 0 | 2.56883473978621 | 0.543 | 0.124 | 0 | MastC | ARHGAP18 |
| 380 | 0 | 2.50027458223343 | 0.524 | 0.155 | 0 | MastC | ACSL4 |
| 381 | 0 | 5.07617280173123 | 0.786 | 0.023 | 0 | Melanocytes | DCT |
| 382 | 0 | 4.49240461232203 | 0.909 | 0.043 | 0 | Melanocytes | PMEL |
| 383 | 0 | 4.37836448896651 | 0.841 | 0.032 | 0 | Melanocytes | TYRP1 |
| 384 | 0 | 4.3648775297787 | 0.884 | 0.07 | 0 | Melanocytes | MITF |
| 385 | 0 | 3.79419078192113 | 0.886 | 0.032 | 0 | Melanocytes | MLANA |
| 386 | 0 | 3.33655224710305 | 0.783 | 0.179 | 0 | Melanocytes | IFI27 |
| 387 | 0 | 2.94848472337057 | 0.703 | 0.049 | 0 | Melanocytes | PDLIM3 |
| 388 | 0 | 2.89701893588777 | 0.885 | 0.234 | 0 | Melanocytes | CRYAB |
| 389 | 0 | 2.71404236228067 | 0.723 | 0.02 | 0 | Melanocytes | GCNT2 |
| 390 | 0 | 2.6410657825678 | 0.735 | 0.111 | 0 | Melanocytes | PHACTR1 |
| 391 | 0 | 2.53100824216055 | 0.649 | 0.005 | 0 | Melanocytes | TRPM1 |
| 392 | 0 | 2.44430142222226 | 0.706 | 0.016 | 0 | Melanocytes | FMN1 |
| 393 | 0 | 2.21355176034964 | 0.797 | 0.277 | 0 | Melanocytes | CDC42EP3 |
| 394 | 0 | 2.15729708864752 | 0.659 | 0.031 | 0 | Melanocytes | TFAP2A |
| 395 | 0 | 2.0607166375331 | 0.792 | 0.163 | 0 | Melanocytes | GPNMB |
| 396 | 0 | 2.02151723344576 | 0.652 | 0.096 | 0 | Melanocytes | PLEKHA5 |
| 397 | 0 | 2.00193375498247 | 0.589 | 0.063 | 0 | Melanocytes | QPCT |
| 398 | 0 | 1.98461045582056 | 0.565 | 0.051 | 0 | Melanocytes | FRZB |
| 399 | 0 | 1.97518105346031 | 0.618 | 0.021 | 0 | Melanocytes | KIT |
| 400 | 0 | 1.90256148737118 | 0.603 | 0.025 | 0 | Melanocytes | NSG1 |
| 401 | 0 | 6.59065712661464 | 0.789 | 0.171 | 0 | Chondrocytes | CLU |
| 402 | 0 | 5.52143387237343 | 0.693 | 0.023 | 0 | Chondrocytes | C2orf40 |
| 403 | 0 | 5.29737037624442 | 0.596 | 0.009 | 0 | Chondrocytes | CYTL1 |
| 404 | 0 | 5.0947020700734 | 0.802 | 0.223 | 0 | Chondrocytes | APOD |
| 405 | 0 | 4.65344485343183 | 0.549 | 0.023 | 0 | Chondrocytes | COCH |
| 406 | 0 | 3.85432295356444 | 0.602 | 0.007 | 0 | Chondrocytes | RBP4 |

| | | | | | | | |
|---|---|---|---|---|---|---|---|
| 407 | 0 | 3.49361395501006 | 0.591 | 0.002 | 0 | Chondrocytes | SNORC |
| 408 | 0 | 3.47345562907293 | 0.588 | 0.021 | 0 | Chondrocytes | FGFBP2 |
| 409 | 0 | 3.34482424000627 | 0.335 | 0.046 | 0 | Chondrocytes | PLA2G2A |
| 410 | 0 | 3.33663029386008 | 0.569 | 0.049 | 0 | Chondrocytes | SERPINA1 |
| 411 | 0 | 3.10261500443301 | 0.727 | 0.194 | 0 | Chondrocytes | SOD3 |
| 412 | 0 | 3.09902882466575 | 0.792 | 0.295 | 0 | Chondrocytes | MGP |
| 413 | 0 | 2.89567199293784 | 0.762 | 0.278 | 0 | Chondrocytes | MT1X |
| 414 | 0 | 2.72618726335502 | 0.857 | 0.238 | 0 | Chondrocytes | NNMT |
| 415 | 0 | 2.56557273337263 | 0.759 | 0.131 | 0 | Chondrocytes | FN1 |
| 416 | 0 | 2.49344448209784 | 0.894 | 0.148 | 0 | Chondrocytes | MEG3 |
| 417 | 0 | 2.48103249896551 | 0.555 | 0.036 | 0 | Chondrocytes | SLPI |
| 418 | 0 | 2.44648325805506 | 0.576 | 0.006 | 0 | Chondrocytes | COL9A3 |
| 419 | 0 | 2.35140431221102 | 0.498 | 0.021 | 0 | Chondrocytes | S100A1 |
| 420 | 0 | 2.34798659074876 | 0.636 | 0.127 | 0 | Chondrocytes | BGN |
| 421 | 0 | 3.45766458782067 | 0.999 | 0.416 | 0 | DC2 | HLA-DPB1 |
| 422 | 0 | 3.42645512396262 | 0.998 | 0.407 | 0 | DC2 | HLA-DPA1 |
| 423 | 0 | 3.09220037973741 | 0.89 | 0.087 | 0 | DC2 | CPVL |
| 424 | 0 | 3.01279421542905 | 1 | 0.59 | 0 | DC2 | CD74 |
| 425 | 0 | 2.8036437159205 | 1 | 0.496 | 0 | DC2 | HLA-DRA |
| 426 | 0 | 2.76052126769937 | 0.96 | 0.666 | 0 | DC2 | TXN |
| 427 | 0 | 2.75353842727868 | 0.999 | 0.455 | 0 | DC2 | HLA-DRB1 |
| 428 | 0 | 2.69109038026844 | 0.792 | 0.231 | 0 | DC2 | HLA-DRB5 |
| 429 | 0 | 2.64552326941928 | 0.988 | 0.26 | 0 | DC2 | HLA-DQA1 |
| 430 | 0 | 2.59302833988216 | 0.833 | 0.019 | 0 | DC2 | DNASE1L3 |
| 431 | 0 | 2.59205087508369 | 0.923 | 0.08 | 0 | DC2 | C1orf54 |
| 432 | 0 | 2.52737865239077 | 0.995 | 0.516 | 0 | DC2 | CST3 |
| 433 | 0 | 2.46853764998852 | 0.823 | 0.201 | 0 | DC2 | PPIF |
| 434 | 0 | 2.45833968861617 | 0.714 | 0.042 | 0 | DC2 | S100B |
| 435 | 0 | 2.4014090732121 | 0.981 | 0.318 | 0 | DC2 | HLA-DQB1 |
| 436 | 0 | 2.35982900501939 | 0.969 | 0.191 | 0 | DC2 | LYZ |
| 437 | 0 | 2.30728383193237 | 0.939 | 0.318 | 0 | DC2 | CTSZ |
| 438 | 0 | 2.09462890409546 | 0.844 | 0.287 | 0 | DC2 | SGK1 |
| 439 | 0 | 2.06845460928201 | 0.903 | 0.221 | 0 | DC2 | BASP1 |
| 440 | 0 | 2.05801647969909 | 0.823 | 0.06 | 0 | DC2 | LGALS2 |

| | | | | | | | |
|---|---|---|---|---|---|---|---|
| 441 | 0 | 6.50694082987718 | 0.928 | 0.031 | 0 | LEC | CCL21 |
| 442 | 0 | 4.82875983175065 | 0.918 | 0.01 | 0 | LEC | TFF3 |
| 443 | 0 | 3.96977973798143 | 0.762 | 0.033 | 0 | LEC | FABP4 |
| 444 | 0 | 3.51434897228565 | 0.838 | 0.03 | 0 | LEC | CLDN5 |
| 445 | 0 | 3.50129019255492 | 0.946 | 0.135 | 0 | LEC | TFPI |
| 446 | 0 | 3.48246549988087 | 0.8 | 0.005 | 0 | LEC | MMRN1 |
| 447 | 0 | 3.31490209288042 | 0.895 | 0.116 | 0 | LEC | GNG11 |
| 448 | 0 | 2.93261382174279 | 0.616 | 0.068 | 0 | LEC | ANGPT2 |
| 449 | 0 | 2.78161332311526 | 0.838 | 0.101 | 0 | LEC | PPFIBP1 |
| 450 | 0 | 2.66354224486309 | 0.651 | 0.106 | 0 | LEC | AKAP12 |
| 451 | 0 | 2.44133979306039 | 0.571 | 0.013 | 0 | LEC | LYVE1 |
| 452 | 0 | 2.41866052489429 | 0.708 | 0.054 | 0 | LEC | RAMP2 |
| 453 | 0 | 2.41751200240576 | 0.665 | 0.022 | 0 | LEC | PROX1 |
| 454 | 0 | 2.32623971967264 | 0.419 | 0.027 | 0 | LEC | C2CD4B |
| 455 | 0 | 2.31759598881207 | 0.947 | 0.348 | 0 | LEC | IGFBP7 |
| 456 | 0 | 2.21235370609766 | 0.491 | 0.021 | 0 | LEC | CAVIN2 |
| 457 | 0 | 2.18512106122603 | 0.635 | 0.021 | 0 | LEC | ECSCR |
| 458 | 0 | 2.15714858014658 | 0.589 | 0.038 | 0 | LEC | SNCG |
| 459 | 0 | 2.08751573087681 | 0.804 | 0.132 | 0 | LEC | TM4SF1 |
| 460 | 1.4722 | 2.17617633715224 | 0.456 | 0.122 | 4.8577650 | LEC | PDK4 |
| 461 | 0 | 8.76118723418525 | 0.865 | 0.236 | 0 | PlasmaC | IGKC |
| 462 | 0 | 8.37959437410917 | 0.62 | 0.135 | 0 | PlasmaC | IGLC2 |
| 463 | 0 | 8.16711270296258 | 0.464 | 0.072 | 0 | PlasmaC | IGLC3 |
| 464 | 0 | 7.71973041456078 | 0.7 | 0.065 | 0 | PlasmaC | IGHG1 |
| 465 | 0 | 7.33085802090256 | 0.515 | 0.056 | 0 | PlasmaC | IGHA1 |
| 466 | 0 | 6.78729134194579 | 0.706 | 0.039 | 0 | PlasmaC | IGHG4 |
| 467 | 0 | 6.53206901330586 | 0.45 | 0.021 | 0 | PlasmaC | IGHG2 |
| 468 | 0 | 6.30303281837895 | 0.689 | 0.04 | 0 | PlasmaC | IGHG3 |
| 469 | 0 | 5.82712633149606 | 0.556 | 0.013 | 0 | PlasmaC | JCHAIN |
| 470 | 0 | 5.67063974848476 | 0.415 | 0.017 | 0 | PlasmaC | IGHGP |
| 471 | 0 | 4.73443932384808 | 0.295 | 0.007 | 0 | PlasmaC | IGHA2 |
| 472 | 0 | 3.64823133441335 | 0.919 | 0.011 | 0 | PlasmaC | MZB1 |
| 473 | 0 | 3.24592073228867 | 0.953 | 0.574 | 0 | PlasmaC | SSR4 |
| 474 | 0 | 2.79692001468341 | 0.968 | 0.585 | 0 | PlasmaC | HERPUD1 |

| | | | | | | | |
|---|---|---|---|---|---|---|---|
| 475 | 0 | 2.62466438044901 | 0.826 | 0.005 | 0 | PlasmaC | DERL3 |
| 476 | 0 | 2.23674573163142 | 0.743 | 0.023 | 0 | PlasmaC | CD79A |
| 477 | 0 | 2.03258706921937 | 0.861 | 0.2 | 0 | PlasmaC | SEC11C |
| 478 | 0 | 1.91419985236179 | 0.932 | 0.576 | 0 | PlasmaC | CYBA |
| 479 | 0 | 1.88653837443592 | 0.794 | 0.132 | 0 | PlasmaC | FKBP11 |
| 480 | 0 | 1.81946238369264 | 0.296 | 0.001 | 0 | PlasmaC | IGLV3-1 |
| 481 | 0 | 5.51705057464824 | 0.663 | 0.026 | 0 | PTGDS+ fibroblasts | COCH |
| 482 | 0 | 3.64887914485224 | 0.875 | 0.133 | 0 | PTGDS+ fibroblasts | PTGDS |
| 483 | 0 | 3.34173438037836 | 0.758 | 0.041 | 0 | PTGDS+ fibroblasts | ASPN |
| 484 | 0 | 3.03692477873514 | 0.858 | 0.168 | 0 | PTGDS+ fibroblasts | LUM |
| 485 | 0 | 2.92530479024869 | 0.782 | 0.178 | 0 | PTGDS+ fibroblasts | IGFBP5 |
| 486 | 0 | 2.92501927596324 | 0.696 | 0.053 | 0 | PTGDS+ fibroblasts | OGN |
| 487 | 0 | 2.6923298080604 | 0.987 | 0.281 | 0 | PTGDS+ fibroblasts | COL1A2 |
| 488 | 0 | 2.66057921104993 | 0.885 | 0.136 | 0 | PTGDS+ fibroblasts | CTSK |
| 489 | 0 | 2.5527728817423 | 0.746 | 0.13 | 0 | PTGDS+ fibroblasts | MFAP4 |
| 490 | 0 | 2.49020953628413 | 0.9 | 0.26 | 0 | PTGDS+ fibroblasts | TCF4 |
| 491 | 0 | 2.44216124221075 | 0.981 | 0.267 | 0 | PTGDS+ fibroblasts | COL1A1 |
| 492 | 0 | 2.4000298346685 | 0.674 | 0.098 | 0 | PTGDS+ fibroblasts | DPT |
| 493 | 0 | 2.36259758494527 | 0.792 | 0.133 | 0 | PTGDS+ fibroblasts | HTRA1 |
| 494 | 0 | 2.34630601942005 | 0.803 | 0.227 | 0 | PTGDS+ fibroblasts | CYR61 |
| 495 | 0 | 2.30981708034088 | 0.398 | 0.071 | 0 | PTGDS+ fibroblasts | POSTN |
| 496 | 0 | 2.29296528686013 | 0.81 | 0.336 | 0 | PTGDS+ fibroblasts | ID3 |
| 497 | 0 | 2.21060205406567 | 0.537 | 0.017 | 0 | PTGDS+ fibroblasts | CRABP1 |
| 498 | 0 | 2.16337580524533 | 0.526 | 0.005 | 0 | PTGDS+ fibroblasts | TNN |
| 499 | 0 | 2.14448051798028 | 0.838 | 0.2 | 0 | PTGDS+ fibroblasts | SPARCL1 |
| 500 | 0 | 2.08980190851712 | 0.715 | 0.105 | 0 | PTGDS+ fibroblasts | THY1 |
| 501 | 0 | 3.89657142078375 | 0.478 | 0.012 | 0 | MigDC | CCL17 |
| 502 | 0 | 3.73181797452988 | 0.665 | 0.019 | 0 | MigDC | CCL22 |
| 503 | 0 | 3.22990018673112 | 0.868 | 0.103 | 0 | MigDC | CCR7 |
| 504 | 0 | 3.12427514921844 | 0.992 | 0.566 | 0 | MigDC | BIRC3 |
| 505 | 0 | 3.04294169778411 | 0.756 | 0.207 | 0 | MigDC | G0S2 |
| 506 | 0 | 3.04204961076576 | 0.977 | 0.667 | 0 | MigDC | TXN |
| 507 | 0 | 2.98882819112485 | 0.964 | 0.265 | 0 | MigDC | CD83 |
| 508 | 0 | 2.69599700195276 | 0.798 | 0.058 | 0 | MigDC | GPR157 |

| | | | | | | | |
|---|---|---|---|---|---|---|---|
| 509 | 0 | 2.68829305995615 | 0.904 | 0.105 | 0 | MigDC | DAPP1 |
| 510 | 0 | 2.59464580509343 | 0.915 | 0.255 | 0 | MigDC | MARCKSL1 |
| 511 | 0 | 2.51368353435081 | 0.847 | 0.121 | 0 | MigDC | FSCN1 |
| 512 | 0 | 2.51361613595539 | 0.968 | 0.42 | 0 | MigDC | HLA-DPB1 |
| 513 | 0 | 2.47188258729292 | 0.827 | 0.052 | 0 | MigDC | POGLUT1 |
| 514 | 0 | 2.46816533111286 | 0.907 | 0.225 | 0 | MigDC | BASP1 |
| 515 | 0 | 2.42802238884829 | 0.489 | 0.015 | 0 | MigDC | IDO1 |
| 516 | 0 | 2.42643576812917 | 0.847 | 0.015 | 0 | MigDC | LAMP3 |
| 517 | 0 | 2.42188078491225 | 0.85 | 0.08 | 0 | MigDC | CSF2RA |
| 518 | 0 | 2.29560980218616 | 0.951 | 0.355 | 0 | MigDC | MARCKS |
| 519 | 0 | 2.27861514864376 | 0.788 | 0.061 | 0 | MigDC | RASSF4 |
| 520 | 0 | 2.20928866633571 | 0.675 | 0.065 | 0 | MigDC | LGALS2 |
| 521 | 0 | 5.07814716870938 | 0.62 | 0.041 | 0 | IL8+ DC1 | IL8 |
| 522 | 0 | 3.89936374159999 | 0.383 | 0.015 | 0 | IL8+ DC1 | IGLL5 |
| 523 | 0 | 3.47830233076511 | 0.361 | 0.013 | 0 | IL8+ DC1 | IGJ |
| 524 | 0 | 2.73155873051111 | 0.476 | 0.012 | 0 | IL8+ DC1 | RP11-1143( |
| 525 | 0 | 2.60797906948086 | 0.617 | 0.042 | 0 | IL8+ DC1 | HLA-DQB2 |
| 526 | 0 | 2.60660641475643 | 0.678 | 0.068 | 0 | IL8+ DC1 | FCER1A |
| 527 | 0 | 2.3469264050843 | 0.622 | 0.097 | 0 | IL8+ DC1 | HLA-DQA2 |
| 528 | 0 | 2.20975393286655 | 0.495 | 0.011 | 0 | IL8+ DC1 | AGPAT9 |
| 529 | 0 | 2.20285201731227 | 0.907 | 0.322 | 0 | IL8+ DC1 | HLA-DQB1 |
| 530 | 0 | 2.18884250544599 | 0.856 | 0.207 | 0 | IL8+ DC1 | G0S2 |
| 531 | 0 | 2.17191816228512 | 0.583 | 0.082 | 0 | IL8+ DC1 | SELK |
| 532 | 0 | 2.1493594068883 | 0.874 | 0.195 | 0 | IL8+ DC1 | LYZ |
| 533 | 0 | 2.13337702130384 | 0.626 | 0.1 | 0 | IL8+ DC1 | ATP5E |
| 534 | 0 | 2.10041900594542 | 0.709 | 0.089 | 0 | IL8+ DC1 | IL1R2 |
| 535 | 1.599( | 2.11982327176099 | 0.943 | 0.42 | 5.2762196 | IL8+ DC1 | HLA-DPB1 |
| 536 | 1.0054 | 2.46747323161587 | 0.979 | 0.519 | 3.3176162 | IL8+ DC1 | CST3 |
| 537 | 2.6678 | 2.40899788048021 | 0.953 | 0.499 | 8.8029435 | IL8+ DC1 | HLA-DRA |
| 538 | 4.979( | 2.3277676982778 | 0.957 | 0.592 | 1.6429010 | IL8+ DC1 | CD74 |
| 539 | 2.802( | 2.9537023467075 | 0.663 | 0.235 | 9.2457595 | IL8+ DC1 | HLA-DRB5 |
| 540 | 5.938( | 2.25606935014473 | 0.842 | 0.387 | 1.9593201 | IL8+ DC1 | INSIG1 |
| 541 | 0 | 4.02236412858995 | 0.707 | 0.001 | 0 | Neuronal_Schwann( | NRXN1 |
| 542 | 0 | 4.0001650365123 | 0.763 | 0.012 | 0 | Neuronal_Schwann( | CDH19 |

| | | | | | | | |
|---|---|---|---|---|---|---|---|
| 543 | 0 | 3.77019196927296 | 0.94 | 0.242 | 0 | Neuronal_Schwann | CRYAB |
| 544 | 0 | 3.57879909971573 | 0.817 | 0.102 | 0 | Neuronal_Schwann | GPM6B |
| 545 | 0 | 3.48318109629984 | 0.635 | 0.046 | 0 | Neuronal_Schwann | S100B |
| 546 | 0 | 3.40307406072423 | 0.529 | 0.01 | 0 | Neuronal_Schwann | SCN7A |
| 547 | 0 | 3.26361477826963 | 0.859 | 0.257 | 0 | Neuronal_Schwann | PMP22 |
| 548 | 0 | 3.11595835946705 | 0.386 | 0.006 | 0 | Neuronal_Schwann | MPZ |
| 549 | 0 | 3.05022979351115 | 0.6 | 0.023 | 0 | Neuronal_Schwann | PLP1 |
| 550 | 0 | 2.7764355805904 | 0.481 | 0.036 | 0 | Neuronal_Schwann | ITGB8 |
| 551 | 0 | 2.62084674849791 | 0.542 | 0.009 | 0 | Neuronal_Schwann | NTM |
| 552 | 0 | 2.50026330300495 | 0.506 | 0.027 | 0 | Neuronal_Schwann | MATN2 |
| 553 | 0 | 2.47303106156721 | 0.523 | 0.038 | 0 | Neuronal_Schwann | CADM1 |
| 554 | 0 | 2.42422415810791 | 0.546 | 0.029 | 0 | Neuronal_Schwann | HSPA12A |
| 555 | 0 | 2.40327628617998 | 0.442 | 0.043 | 0 | Neuronal_Schwann | ANK3 |
| 556 | 9.0901 | 2.54955269062491 | 0.793 | 0.222 | 2.9994017 | Neuronal_Schwann | DST |
| 557 | 1.1805 | 2.46565437708094 | 0.494 | 0.088 | 3.8953028 | Neuronal_Schwann | RCAN1 |
| 558 | 6.3090 | 2.37389065937943 | 0.83 | 0.273 | 2.0817328 | Neuronal_Schwann | SPARC |
| 559 | 8.0703 | 2.52281627632109 | 0.847 | 0.333 | 2.6629043 | Neuronal_Schwann | CD9 |
| 560 | 9.1923 | 2.69795640215635 | 0.598 | 0.181 | 3.0331208 | Neuronal_Schwann | IGFBP5 |
| 561 | 0 | 6.1809828500975 | 0.97 | 0.011 | 0 | SMC | DES |
| 562 | 0 | 5.60040943954112 | 0.976 | 0.011 | 0 | SMC | ACTG2 |
| 563 | 0 | 5.54210890225559 | 0.994 | 0.104 | 0 | SMC | ACTA2 |
| 564 | 0 | 5.37952021653091 | 0.988 | 0.09 | 0 | SMC | MYLK |
| 565 | 0 | 4.94939996468265 | 0.988 | 0.056 | 0 | SMC | MYH11 |
| 566 | 0 | 4.58356614863666 | 0.994 | 0.023 | 0 | SMC | CNN1 |
| 567 | 0 | 4.54565290580232 | 0.88 | 0.001 | 0 | SMC | PCP4 |
| 568 | 0 | 4.16644408597815 | 0.988 | 0.125 | 0 | SMC | CSRP1 |
| 569 | 0 | 3.67997606176232 | 0.964 | 0.047 | 0 | SMC | PPP1R14A |
| 570 | 0 | 3.62541419474664 | 0.843 | 0.077 | 0 | SMC | SELM |
| 571 | 0 | 3.37658633751743 | 0.614 | 0.004 | 0 | SMC | ACTA1 |
| 572 | 0 | 3.21730623988112 | 0.873 | 0.063 | 0 | SMC | SMTN |
| 573 | 0 | 3.05328844608668 | 0.795 | 0.043 | 0 | SMC | NRN1 |
| 574 | 0 | 2.95589517545024 | 0.783 | 0.057 | 0 | SMC | SYNPO2 |
| 575 | 9.7598 | 5.5635682077539 | 0.994 | 0.152 | 3.2203675 | SMC | TAGLN |
| 576 | 4.5451 | 5.51753737769328 | 1 | 0.162 | 1.4997140 | SMC | TPM2 |

| | | | | | | | |
|---|---|---|---|---|---|---|---|
| 577 | 4.2985 | 5.28632992161244 | 0.994 | 0.165 | 1.4183429 | SMC | MYL9 |
| 578 | 4.9380 | 2.97978677071222 | 0.91 | 0.144 | 1.6293516 | SMC | CKB |
| 579 | 8.5687 | 4.16162141881816 | 1 | 0.2 | 2.8273461 | SMC | TPM1 |
| 580 | 3.4632 | 3.08050770617349 | 1 | 0.469 | 1.1427335 | SMC | DSTN |
| 581 | 0 | 8.18547545646324 | 0.969 | 0.005 | 0 | Skeletal muscle cell | ACTA1 |
| 582 | 0 | 7.38558923692975 | 0.875 | 0.01 | 0 | Skeletal muscle cell | TNNC2 |
| 583 | 0 | 7.05905205042728 | 0.875 | 0.014 | 0 | Skeletal muscle cell | MYLPF |
| 584 | 0 | 7.05217681541108 | 0.969 | 0.004 | 0 | Skeletal muscle cell | MB |
| 585 | 0 | 7.04220814026433 | 0.969 | 0.002 | 0 | Skeletal muscle cell | MYL1 |
| 586 | 0 | 7.02109465746987 | 0.938 | 0.002 | 0 | Skeletal muscle cell | CKM |
| 587 | 0 | 6.95341590386022 | 0.938 | 0.006 | 0 | Skeletal muscle cell | TCAP |
| 588 | 0 | 6.93415396684556 | 0.875 | 0.009 | 0 | Skeletal muscle cell | TNNI2 |
| 589 | 0 | 6.74253382219968 | 0.844 | 0.011 | 0 | Skeletal muscle cell | TNNT3 |
| 590 | 0 | 6.43417083941691 | 0.938 | 0.001 | 0 | Skeletal muscle cell | CSRP3 |
| 591 | 0 | 6.12097572732393 | 1 | 0.007 | 0 | Skeletal muscle cell | ENO3 |
| 592 | 0 | 6.09714754726781 | 0.969 | 0.001 | 0 | Skeletal muscle cell | COX6A2 |
| 593 | 0 | 5.9965124307846 | 1 | 0.012 | 0 | Skeletal muscle cell | DES |
| 594 | 0 | 5.46419916570855 | 0.344 | 0.001 | 0 | Skeletal muscle cell | MYL2 |
| 595 | 0 | 5.42331525270779 | 0.812 | 0.001 | 0 | Skeletal muscle cell | ANKRD1 |
| 596 | 0 | 5.24914562303869 | 0.906 | 0.002 | 0 | Skeletal muscle cell | SLN |
| 597 | 0 | 5.2387067463363 | 0.781 | 0.001 | 0 | Skeletal muscle cell | MYH2 |
| 598 | 3.0162 | 5.13044791062127 | 0.469 | 0.007 | 9.9522906 | Skeletal muscle cell | TNNT1 |
| 599 | 7.1916 | 5.62275034810634 | 1 | 0.162 | 2.3729558 | Skeletal muscle cell | TPM2 |
| 600 | 1.5795 | 6.26709181297112 | 0.844 | 0.201 | 5.2119970 | Skeletal muscle cell | TPM1 |
