## Supplementary Table 2 for "Multi-scale spatial mapping of cell populations across anatomical sites in healthy human skin and basal cell carcinoma"

Supplementary Table 2 : List of custom genes for ISS

| Gene full name | Gene Symbol | Gene ID | Aliases | ENSEMBL GeneID | NCBI Accession number |
|---|---|---|---|---|---|
| keratin 5 | KRT5 | 3852 | K5; CK5; DDD; DDD1; EBS2; KRT5A | ENSG00000186081 | NG_008297.1 |
| keratin 14 | KRT14 | 3861 | K14; NFJ; CK14; EBS3; EBS4 | ENSG00000186847 | NG_008624.1 |
| caveolin 1 | CAV1 | 857 | CGL3; PPH3; BSCL3; LCCNS; VIP21; MSTP085 | ENSG00000105974 | NG_012051.1 |
| caveolin 2 | CAV2 | 858 | CAV | ENSG00000105971 | NG_029920.1 |
| collagen type XVII alpha 1 chain | COL17A1 | 1308 | ERED; BP180; BPA-2; BPAG2; LAD-1; BA16H23.2 | ENSG00000065618 | NG_007069.1 |
| integrin subunit beta 1 | ITGB1 | 3688 | CD29; FNRB; MDF2; VLAB; GPIIA; MSK12; VLA-BETA | ENSG00000150093 | NG_029012.1 |
| CD46 molecule | CD46 | 4179 | MCP; TLX; AHUS2; MIC10; TRA2.10 | ENSG00000117335 | NG_009296.1 |
| delta like canonical Notch ligand 1 | DLL1 | 28514 | DL1; Delta; DELTA1; NEDBAS | ENSG00000198719 | NG_027940.1 |
| leucine rich repeats and immunoglobulin like domains 1 | LRIG1 | 26018 | LIG1; LIG-1 | ENSG00000144749 | not found |
| mitogen-activated protein kinase 1 | MAPK1 | 5594 | ERK; p38; p40; p41; ERK2; ERT1; NS13; ERK-2; MAPK2; PRKM1; PRKM2; P42MAPK; p41mapk; p42-MAPK | ENSG00000100030 | NG_023054.2 |
| keratin 10 | KRT10 | 3858 | BIE; EHK; K10; KPP; BCIE; CK10 | ENSG00000186395 | NG_008405.1 |
| keratin 1 | KRT1 | 3848 | CK1, EHK, EHK1, EPPK, K1, KRT1A, NEPPK | ENSG00000167768 | NG_008364.2 |

| | | | | | |
|---|---|---|---|---|---|
| cyclin dependent kinase 1 | CDK1 | 983 | CDC2, CDC28A, P34CDC2 | ENSG00000170312 | NG_029877.1 |
| proliferating cell nuclear antigen | PCNA | 5111 | ATLD2 | ENSG00000132646 | NG_047066.1 |
| intercellular adhesion molecule 1 | ICAM1 | 3383 | BB2, CD54, P3.58 | ENSG00000090339 | NG_012083.1 |
| C-C motif chemokine ligand 20 | CCL20 | 6364 | CKb4; LARC; ST38; MIP3A; Exodus; MIP-3a; SCYA20; MIP-3-alpha | ENSG00000115009 | not found |
| lymphocyte antigen 6 family member D | LY6D | 8581 | E48; Ly-6D | ENSG00000167656 | not found |
| keratin 17 | KRT17 | 3872 | PC; K17; PC2; 39.1; CK-17; PCHC1 | ENSG00000128422 | NG_008625.1 |
| embigin | EMB | 133418 | GP70 | ENSG00000170571 | not found |
| microsomal glutathione S-transferase 1 | MGST1 | 4257 | MGST; GST12; MGST-I | ENSG00000008394 | not found |
| Tyrosinase-related protein 1 | TRYP1 | 7306 | TRP; CAS2; CATB; GP75; OCA3; TRP1; TYRP; b-PROTEIN | ENSG00000107165 | NG_011705.1 |
| premelanosome protein | PMEL | 6490 | P1; SI; SIL; ME20; P100; SILV; ME20M; gp100; ME20-M; PMEL17; D12S53E | ENSG00000185664 | NG_028086.1 |
| formin 1 | FMN1 | 342184 | LD; FMN | ENSG00000248905 | NG_042863.1 |
| calpain 3 | CAPN3 | 825 | p94; CANP3; LGMD2; nCL-1; CANPL3; LGMD2A; LGMDD4; LGMDR1 | ENSG00000092529 | NG_008660.1 |
| empty spiracles homeobox 2 | EMX2 | 2018 | | ENSG00000170370 | NG_013009.1 |
| major facilitator superfamily domain containing 12 | MFSD12 | 126321 | PP3501; C19orf28 | ENSG00000161091 | not found |

| | | | | | |
|---|---|---|---|---|---|
| myelin protein zero | MPZ | 4359 | P0; CHM; DSS; MPP; CHN2; CMT1; CMT1B; CMT2I; CMT2J; CMT4E; CMTDI3; CMTDID; HMSNIB | ENSG00000158887 | NG_008055.1 |
| dickkopf WNT signaling pathway inhibitor 3 | DKK3 | 27122 | RIG; REIC | ENSG00000050165 | not found |
| dermcidin | DCD | 117159 | PIF; AIDD; DSEP; HCAP; DCD-1 | ENSG00000161634 | not found |
| keratin 19 | KRT19 | 3880 | K19; CK19; K1CS | ENSG00000171345 | NG_012285.1 |
| sodium voltage-gated channel alpha subunit 7 | SCN7A | 6332 | NaG; SCN6A; Nav2.1; Nav2.2 | ENSG00000136546 | NG_031928.1 |
| myogenic factor 5 | MYF5 | 4617 | EORVA; bHLHc2 | ENSG00000111049 | not found |
| decorin | DCN | 1634 | CSCD; PG40; PGII; PGS2; DSPG2; SLRR1B | ENSG00000011465 | NG_011672.1 |
| platelet derived growth factor receptor alpha | PDGFRA | 5156 | CD140A; PDGFR2; PDGFR-2 | ENSG00000134853 | NG_009250.1 |
| collagen type I alpha 1 chain | COL1A1 | 1277 | OI1; OI2; OI3; OI4; EDSC; CAFYD; EDSARTH1 | ENSG00000108821 | NG_007400.1 |
| fibroblast growth factor 7 | FGF7 | 2252 | KGF; HBGF-7 | ENSG00000140285 | NG_029159.1 |
| collagen type VI alpha 5 chain | COL6A5 | 256076 | VWA4; COL29A1 | ENSG00000172752 | NG_021424.1 |
| prostaglandin D2 synthase | PTGDS | 5730 | PDS; PGD2; PGDS; LPGDS; PGDS2; L-PGDS | ENSG00000107317 | not found |
| APC down-regulated 1 | APCDD1 | 147495 | HHS; HTS; B7323; HYPT1; DRAPC1; FP7019 | ENSG00000154856 | NG_027685.1 |
| C-C motif chemokine ligand 2 | CCL2 | 6347 | HC11; MCAF; MCP1; MCP-1; SCYA2; GDCF-2; SMC-CF; HSMCR30 | ENSG00000108691 | NG_012123.1 |
| apolipoprotein D | APOD | 347 | none | ENSG00000189058 | not found |

| | | | | | |
|---|---|---|---|---|---|
| asporin | ASPN | 54829 | OS3; PLAP1; PLAP-1; SLRR1C | ENSG00000106819 | NG_023430.2 |
| WNT inhibitory factor 1 | WIF1 | 11197 | WIF-1 | ENSG00000156076 | not found |
| periostin | POSTN | 10631 | PN; OSF2; OSF-2; PDLPOSTN | ENSG00000133110 | not found |
| secretory leukocyte peptidase inhibitor | SLPI | 6590 | ALP; MPI; ALK1; BLPI; HUSI; WAP4; WFDC4; HUSI-I | ENSG00000124107 | NG_028137.1 |
| cellular communication network factor 5 | CCN5 | 8839 | CT58; WISP2; CTGF-L | ENSG00000064205 | not found |
| microfibril associated protein 5 | MFAP5 | 8076 | AAT9; MP25; MAGP2; MAGP-2; MFAP-5 | ENSG00000197614 | not found |
| prostaglandin-endoperoxide synthase | PTGS2 | 5743 | COX2; COX-2; PHS-2; PGG/HS; PGHS-2; hCox-2; GRIPGHS | ENSG00000073756 | NG_028206.2 |
| HIC ZBTB transcriptional repressor 1 | HIC1 | 3090 | hic-1; ZBTB29; ZNF901 | ENSG00000177374 | NG_027689.1 |
| secreted frizzled related protein 1 | SFRP1 | 6422 | FRP; FRP1; FrzA; FRP-1; SARP2 | ENSG00000104332 | not found |
| platelet and endothelial cell adhesion molecule 1 | PECAM1 | 5175 | CD31; PECA1; GPIIA'; PECAM-1; endoCAM; CD31/EndoCAM | ENSG00000261371 | NG_047009.1 |
| selectin E | SELE | 6401 | ELAM; ESEL; CD62E; ELAM1; LECAM2 | ENSG00000007908 | NG_012124.1 |
| claudin 5 | CLDN5 | 7122 | AWAL; BEC1; TMVCF; TMDVCF; CPETRL1 | ENSG00000184113 | not found |
| fms related receptor tyrosine kinase 1 | FLT1 | 2321 | FLT; FLT-1; VEGFR1; VEGFR-1 | ENSG00000102755 | NG_012003.1 |
| hes related family bHLH transcription factor with YRPW motif 1 | HEY1 | 23462 | CHF2; OAF1; HERP2; HESR1; HRT-1; NERP2; hHRT1; BHLHb31 | ENSG00000164683 | not found |
| CD93 molecule | CD93 | 22918 | C1QR1; C1qRP; CDw93; | ENSG00000125810 | not found |

| | | | | | |
|---|---|---|---|---|---|
| | | | ECSM3; MXRA4; C1qR(P); dJ737E23.1 | | |
| endomucin | EMCN | 51705 | EMCN2; MUC14 | ENSG00000164035 | not found |
| interferon alpha inducible protein 27 | IFI27 | 3429 | P27; ISG12; FAM14D; ISG12A | ENSG00000165949 | not found |
| atypical chemokine receptor 1 (Duffy blood group) | ACKR1 | 2532 | FY; Dfy; GPD; DARC; GpFy; CCBP1; CD234; WBCQ1; DARC/ACKR1 | ENSG00000213088 | NG_011626.3 |
| synuclein gamma | SNCG | 6623 | SR; BCSG1 | ENSG00000173267 | NG_008783.1 |
| lymphatic vessel endothelial hyaluronan receptor 1 | LYVE1 | 10894 | HAR; XLKD1; LYVE-1; CRSBP-1 | ENSG00000133800 | not found |
| prospero homeobox 1 | PROX1 | 5629 | none | ENSG00000117707 | not found |
| C-C motif chemokine ligand 21 | CCL21 | 6366 | ECL; SLC; CKb9; TCA4; 6Ckine; SCYA21 | ENSG00000137077 | not found |
| caveolae associated protein 2 | CAVIN2 | 8436 | SDR; SDPR; PS-p68; cavin-2 | ENSG00000168497 | not found |
| multimerin 1 | MMRN1 | 22915 | ECM; MMRN; GPIa*; EMILIN4 | ENSG00000138722 | NG_032895.2 |
| actin alpha 2, smooth muscle | ACTA2 | 59 | ACTSA | ENSG00000107796 | NG_011541.1 |
| transgelin | TAGLN | 6876 | SM22; SMCC; TAGLN1; WS3-10; SM22-alpha | ENSG00000149591 | not found |
| RERG like | RERGL | 79785 | none | ENSG00000111404 | NG_052618.1 |
| myosin heavy chain 11 | MYH11 | 4629 | AAT4, FAA4, SMHC, SMMHC | ENSG00000133392 | NG_009299.1 |
| metalloreductase | STEAP4 | 79689 | TIARP; STAMP2; SchLAH; TNFAIP9 | ENSG00000127954 | NG_028313.1 |
| regulator of G protein signaling 5 | RGS5 | 8490 | MST092; MST106; MST129; MSTP032; MSTP092; MSTP106; MSTP129 | ENSG00000143248 | NG_027731.2 |

| | | | | | |
|---|---|---|---|---|---|
| desmin | DES | 1674 | CSM1; CSM2; CDCD3; LGMD1D; LGMD1E; LGMD2R | ENSG00000175084 | NG_008043.1 |
| actin gamma 2, smooth muscle | ACTG2 | 72 | ACT; ACTE; VSCM; ACTA3; ACTL3; ACTSG | ENSG00000163017 | NG_034140.1 |
| T cell immunoreceptor with Ig and ITIM domains | TIGIT | 201633 | VSIG9; VSTM3; WUCAM | ENSG00000181847 | not found |
| SPARC (osteonectin), cwcv and kazal like domains proteoglycan 2 | SPOCK2 | 9806 | testican-2 | ENSG00000107742 | not found |
| C-type lectin domain family 2 member D | CLEC2D | 29121 | CLAX; LLT1; OCIL | ENSG00000069493 | not found |
| cathepsin W | CTSW | 1521 | LYPN | ENSG00000172543 | not found |
| RUNX family transcription factor 3 | RUNX3 | 864 | AML2; CBFA3; PEBP2aC | ENSG00000020633 | not found |
| serpin family B member 2 | SERPINB2 | 5055 | PAI; PAI2; PAI-2; PLANH2; HsT1201 | ENSG00000197632 | not found |
| twist family bHLH transcription factor 1 | TWIST1 | 7291 | CRS; CSO; SCS; ACS3; CRS1; BPES2; BPES3; SWCOS; TWIST; bHLHa38 | ENSG00000122691 | NG_008114.2 |
| SRY-box transcription factor 10 | SOX10 | 6663 | DOM; WS4; PCWH; WS2E; WS4C | ENSG00000100146 | NG_007948.1 |
