## Supplementary Table 3 for "Multi-scale spatial mapping of cell populations across anatomical sites in healthy human skin and basal cell carcinoma"

Supplementary Table 3: List of predefined Immune General gene panel

| Gene full name | Gene symbol | HGNC ID | Alias | ENSEMBL GeneID human | Annotation |
|---|---|---|---|---|---|
| protein tyrosine phosphatase receptor type C | PTPRC | HGNC: 9666 | B220;CD45R;Cd45;L-CA;Ly-5;Lyt-4;T200;loc;B220;CD45;CD45R;GP180;L-CA;LCA;LY5;T200 | ENSG00000081237 | hematopoietic lineage |
| CD34 molecule | CD34 | HGNC: 1662 | AU040960 | ENSG00000174059 | hematopoietic stem cells and endothelial cells |
| CD40 molecule | CD40 | HGNC: 11919 | AI326936;Bp50;GP39;HIGM1;IGM;IMD3;T-BAM;TRAP;Tnfrsf5;p50;Bp50;CDW40;TNFRSF5;p50 | ENSG00000101017 | antigen presenting cells (APC) |
| CD80 molecule | CD80 | HGNC: 1700 | B71;Cd28l;Ly-53;Ly53;MIC17;TSA1;B7;B7-1;B7.1;BB1;CD28LG;CD28LG1;LAB7 | ENSG00000121594 | antigen presenting cells (APC) |
| Fc receptor, IgG, high affinity I | | | AI323638;AV092959;CD64;FcgammaRI;IGGHAFC | | macrophages and monocytes |
| Fc fragment of IgG receptor Ia | FCGR1A | HGNC: 3613 | CD64;CD64A;FCRI;IGFR1 | ENSG00000150337 | macrophages and monocytes |
| S100 calcium binding protein A8 | S100A8 | HGNC: 10498 | 60B8Ag;AI323541;B8Ag;CFAg;CP-10;Caga;MRP8;p8;60B8AG;CAGA;CFAG;CGLA;CP-10;L1Ag;MA387;MIF;MRP8;NIF;P8 | ENSG00000143546 | monocytes |
| CD14 molecule | CD14 | HGNC: 1628 | | ENSG00000170458 | CD14+ monocytes |
| Fc receptor, IgG, low affinity III | | | CD16 | | CD16+ monocytes |
| Fc fragment of IgG receptor IIIa | FCGR3A | HGNC: 3619 | CD16;CD16A;FCG3;FCGR3;FCGRIII;FCR-10;FCRIII;FCRIIIA;IGFR3;IMD20 | ENSG00000203747 | CD16+ monocytes |

| | | | | | |
|---|---|---|---|---|---|
| membrane spanning 4-domains A2 | MS4A2 | HGNC: 7316 | FcRB;Fce1b;Fcer1b;Fcrbeta;Ms4a1;fcERI;APY;ATOPY;FCER1B;FCERI;IGEL;IGER;IGHER;MS4A1 | ENSG00000149534 | mast cells |
| tryptase alpha/beta 1 | TPSAB1 | HGNC: 12019 | MMCP-7;Mcp-7;Mcp7;Mcpt7;TPS1;TPS2;TPSB1;TPSB2;Tryptase-2 | ENSG00000172236 | mast cells |
| apolipoprotein E | APOE | HGNC: 613 | AI255918;Apo-E;AD2;APO-E;ApoE4;LDLCQ5;LPG | ENSG00000130203 | macrophages |
| complement C1q A chain | C1QA | HGNC: 1241 | AI255395;Adic;C1q | ENSG00000173372 | macrophages |
| complement C1q B chain | C1QB | HGNC: 1242 | Adia | ENSG00000173369 | macrophages |
| CD163 molecule | CD163 | HGNC: 1631 | CD163v2;CD163v3;M130;MM130;SCARI1 | ENSG00000177575 | macrophages |
| CD68 molecule | CD68 | HGNC: 1693 | Lamp4;Scard1;gp110;GP110;LAMP4;SCARD1 | ENSG00000129226 | macrophages |
| folate receptor beta | FOLR2 | HGNC: 3793 | FBP2;FR-P3;FR-beta;Folbp-2;Folbp2;BETA-HFR;FBP;FBP/PL-1;FOLR1;FR-BETA;FR-P3;FRbeta | ENSG00000165457 | resident macrophages |
| CEA cell adhesion molecule 8 | CEACAM8 | HGNC: 1820 | CD66b;CD67;CGM6;NCA-95 | ENSG00000124469 | granulocytes |
| fucosyltransferase 4 | FUT4 | HGNC: 4015 | AI451562;CD15;FAL;FucT-IV;LeX;SSEA-1;Ssea1;CD15;ELFT;FCT3A;FUC-TIV;FUTIV;LeX;SSEA-1 | ENSG00000196371 | neutrophils and eosinophils |
| interleukin 3 receptor subunit alpha | IL3RA | 08385 | CD123;CDw123;SUT-1;CD123;IL3R;IL3RAY;IL3RX;IL3RY;hIL-3Ra | 430000 | basophils and plasmacytoid dendritic cells (pDC) |
| CD209 molecule | CD209 | HGNC: 1641 | CDSIGN;CLEC4L;DC-SIGN;DC-SIGN1 | ENSG00000090659 | dendritic cells (DC) |
| CD209a antigen | | | CD209;CDSIGN;CIRE;DC-SIGN;DC-SIGN1;Dcsign;SIGN-R1;SIGNR5 | | dendritic cells (DC) |
| integrin subunit alpha X | ITGAX | HGNC: 6152 | AI449405;Cd11c;Cr4;N418;CD11C;SLEB6 | ENSG00000140678 | dendritic cells (DC) |
| C-type lectin domain containing 9A | CLEC9A | HGNC: 26705 | 9830005G06Rik;DNGR-1;CD370;DNGR-1;DNGR1;UNQ9341 | ENSG00000197992 | type 1 dendritic cells (DC1) |
| CD1c molecule | CD1C | HGNC: 1636 | BDCA1;CD1;CD1A;R7 | ENSG00000158481 | type 2 dendriti |

| | | | | | |
|---|---|---|---|---|---|
| | | | | | c cells (DC2) |
| C-type lectin domain containing 10A | CLEC10A | HGNC: 16916 | CD301a;M-ASGP-BP-1;Mgl;Mgl1;CD301;CLECSF13;CLECSF14;HML;HML2;MGL | ENSG00000132514 | type 2 dendritic cells (DC2) |
| indoleamine 2,3-dioxygenase 1 | IDO1 | HGNC: 6059 | Ido;Indo;IDO;IDO-1;INDO | ENSG00000131203 | IDO1+ myeloid dendritic cells (mDC) |
| leukocyte specific transcript 1 | LST1 | HGNC: 14189 | B144;B144;D6S49E;LST-1 | ENSG00000204482 | innate lymphoid cells (ILC) |
| CD7 molecule | CD7 | HGNC: 1695 | GP40;LEU-9;TP41;Tp40 | ENSG00000173762 | natural killer (NK) cells |
| killer cell lectin like receptor C2 | KLRC2 | HGNC: 6375 | NKG2C;CD159c;NKG2-C;NKG2C | ENSG00000205809 | natural killer (NK) cells |
| killer cell lectin like receptor D1 | KLRD1 | HGNC: 6378 | CD94;CD94 | ENSG00000134539 | natural killer (NK) cells |
| transmembrane immune signaling adaptor TYROBP | TYROBP | HGNC: 12449 | DAP12;KARAP;Ly83;DAP12;KARAP;PLOSL;PLOSL1 | ENSG00000011600 | natural killer (NK) cells |
| natural killer cell granule protein 7 | NKG7 | HGNC: 7830 | 2500004F03Rik;GIG1;GMP-17;p15-TIA-1 | ENSG00000105374 | natural killer (NK) cells/ NKT cells |
| CD3d molecule | CD3D | HGNC: 1673 | T3d;CD3-DELTA;IMD19;T3D | ENSG00000167286 | T cells |
| CD3e molecule | CD3E | HGNC: 1674 | AI504783;CD3;CD3epsilon;T3e;IMD18;T3E;TCRE | ENSG00000198851 | T cells |
| CD3g molecule | CD3G | HGNC: 1675 | Ctg-3;Ctg3;T3g;CD3-GAMMA;IMD17;T3G | ENSG00000160654 | T cells |
| CD28 molecule | CD28 | HGNC: 1653 | Tp44 | ENSG00000178562 | T cells, T cell activation |
| CD4 molecule | CD4 | HGNC: 1678 | L3T4;Ly-4;CD4mut | ENSG00000010610 | CD4+ T cells |
| CD8a molecule | CD8A | HGNC: 1706 | BB154331;Ly-2;Ly-35;Ly-B;Lyt-2;CD8;Leu2;p32 | ENSG00000153563 | CD8+ T cells |
| CD8b molecule | CD8B | HGNC: 1707 | CD8B1;LEU2;LY3;LYT3;P37 | ENSG00000172116 | CD8+ T cells |

| | | | | | |
|---|---|---|---|---|---|
| CD8 antigen, beta chain 1 | | | Cd8b;Ly-3;Ly-C;Lyt-3 | | CD8+ T cells |
| interleukin 2 receptor subunit alpha | IL2RA | HGNC: 6008 | CD25;Il2r;Ly-43;CD25;IDDM10;IL2R;IMD41;TCGFR;p55 | ENSG00000134460 | mouse regulatory T cells (Treg); human Th1 effector |
| killer cell lectin like receptor B1 | KLRB1 | HGNC: 6373 | 4930431A04Rik;Gm4696;Klrb1g;Klrb6;Ly-55;Ly55;NKR-P1G;Nkrp-1e;Nkrp1g;CD161;CLEC5B;NKR;NKR-P1;NKR-P1A;NKRP1A;hNKR-P1A | ENSG00000111796 | mucosal-associated invariant T cells (MAIT) |
| forkhead box P3 | FOXP3 | HGNC: 6106 | JM2;scurfin;sf;AIID;DIETER;IPEX;JM2;PIDX;XPID | ENSG00000049768 | regulatory T cells (Treg) |
| C-X-C motif chemokine receptor 3 | CXCR3 | HGNC: 4540 | Cd183;Cmkar3;CD182;CD183;CKR-L2;CMKAR3;GPR9;IP10-R;Mig-R;MigR | ENSG00000186810 | type 1 T helper cells (Th1) |
| C-C motif chemokine receptor 4 | CCR4 | HGNC: 1605 | C-C CKR-4;CHEMR1;Cmkbr4;LESTR;Sdf1r;CC-CKR-4;CD194;CKR4;CMKBR4;ChemR13;HGCN:14099;K5-5 | ENSG00000183813 | type 2 T helper cells (Th2) |
| joining chain of multimeric IgA and IgM | JCHAIN | HGNC: 5713 | 9530090F24Rik;AI323815;Igj;Jch;IGCJ;IGJ;JCH | ENSG00000132465 | mucosal-targeted plasma cells |
| G protein-coupled receptor 183 | GPR183 | HGNC: 3128 | Ebi2;EBI2;hEBI2 | ENSG00000169508 | memory cells |
| CD19 molecule | CD19 | HGNC: 1633 | AW495831;B4;CVID3 | ENSG00000177455 | B cells |
| CD79a molecule | CD79A | HGNC: 1698 | Ig-alpha;Iga;Igalpha;Ly-54;Ly54;mb-1;IGA;MB-1 | ENSG00000105369 | B cells |
| CD79b molecule | CD79B | HGNC: 1699 | B29;Ig-beta;Igb;Igbeta;AGM6;B29;IGB | ENSG00000007312 | B cells |
| membrane spanning 4-domains A1 | MS4A1 | HGNC: 7315 | AA960661;Cd20;Ly-44;Ms4a2;B1;Bp35;CD20;CVID5;LEU-16;MS4A2;S7 | ENSG00000156738 | B cells |

| | | | | | |
|---|---|---|---|---|---|
| immunoglobulin kappa constant | IGKC | HGNC: 5716 | Igk-C;HCAK1;IGKCD;Km | ENSG00000211592 | plasma cells |
