## Supplementary Table 4 for "Multi-scale spatial mapping of cell populations across anatomical sites in healthy human skin and basal cell carcinoma"

### Supplementary Table 3: List of predefined Immune Regulation gene panel

| Gene full name | Gene symbol | HGNC ID | Alias | ENSEMBL GeneID human | Annota-tion |
|---|---|---|---|---|---|
| interleukin 7 | IL7 | HGNC:6023 | A630026I06Rik;Il-7;hlb368;IL-7 | ENSG00000104432 | differentiation of hematopoietic stem cells into lymphoid progenitor cells |
| C-C motif chemokine receptor 5 | CCR5 | HGNC:1606 | AM4-7;CD195;Cmkbr5;CC-CKR-5;CCCKR5;CCR-5;CD195;CKR-5;CKR5;CMKBR5;IDDM22 | ENSG00000160791 | granulocyte lineage differentiation |
| basic leucine zipper ATF-like transcription factor 3 | BATF3 | HGNC:28915 | 9130211I03Rik;Snft;JDP1;JUNDM1;SNFT | ENSG00000123685 | dendritic cell (DC) development, potential target in immunosupressed tumor |
| interleukin 7 receptor | IL7R | HGNC:6024 | CD127;IL-7Ralpha;CD127;CDW127;IL-7R-alpha;IL7RA;ILRA | ENSG00000168685 | V(D)J recombination |
| lymphoid enhancer binding factor 1 | LEF1 | HGNC:6551 | 3000002B05;AI451430;Lef-1;LEF-1;TCF10;TCF1ALPHA;TCF7L3 | ENSG00000138795 | B and T cell differentiation |
| C-C motif chemokine receptor 6 | CCR6 | HGNC:1607 | CC-CKR-6;CCR-6;Cmkbr6;KY411;BN-1;C-C CKR-6;CC-CKR-6;CCR-6;CD196;CKR-L3;CKRL3;CMKBR6;DCR2;DRY6;GPR29;GPRCY4;STRL22 | ENSG00000112486 | B cell maturation and differentation |
| interleukin 2 | IL2 | HGNC:6001 | Il-2;IL-2;TCGF;lymphokine | ENSG00000109471 | T cell differentiation and immune tolerance, Th1 effector |

| | | | | | |
|---|---|---|---|---|---|
| neural cell adhesion molecule 1 | NCAM1 | HGNC:7656 | CD56;E-NCAM;NCAM-1;Ncam;CD56;MSK39;NCAM | ENSG00000149294 | T cell expansion, neural development |
| T-box transcription factor 21 | TBX21 | HGNC:11599 | TBT1;Tbet;Tblym;T-PET;T-bet;TBET;TBLYM | ENSG00000073861 | controls expression of Th1 cytokine IFNG |
| interferon gamma | IFNG | HGNC:5438 | IFN-g;Ifg;IFG;IFI | ENSG00000111537 | Th1 development, diverse functions |
| interleukin 5 | IL5 | HGNC:6016 | Il-5;EDF;IL-5;TRF | ENSG00000113525 | Th2 development |
| interleukin 4 | IL4 | HGNC:6014 | BSF-1;Il-4;BCGF-1;BCGF1;BSF-1;BSF1;IL-4 | ENSG00000113520 | Th2 development, diverse functions |
| GATA binding protein 3 | GATA3 | HGNC:4172 | Gata-3;jal;HDR;HDRS | ENSG00000107485 | Th2 helper cell development |
| C-C motif chemokine receptor 7 | CCR7 | HGNC:1608 | CC-CKR-7;CCR-7;CD197;Cdw197;Cmkbr7;EBI1;Ebi1h;BLR2;CC-CKR-7;CCR-7;CD197;CDw197;CMKBR7;EBI1 | ENSG00000126353 | dendritic cell (DC) activation |
| CD44 molecule (Indian blood group) | CD44 | HGNC:1681 | AU023126;AW121933;AW146109;HERMES;Ly-24;Pgp-1;CDW44;CSPG8;ECMR-III;HCELL;HUTCH-I;IN;LHR;MC56;MDU2;MDU3;MIC4;Pgp1 | ENSG00000026508 | lymphocyte activation |
| peroxisome proliferator activated receptor gamma | PPARG | HGNC:9236 | Nr1c3;PPAR-gamma;PPAR-gamma2;PPARgamma;PPARgamma2;CIMT1;GLM1;NR1C3;PPARG1;PPARG2;PPARG5;PPARgamma | ENSG00000132170 | M2 macrophage activation, lipotoxicity |
| icos ligand | | | AU044799;B7-H2;B7RP-1;B7h;BG071784;Gl50;GL50;GL50-B;ICOS-L;Icoslg;LICOS;Ly115l;mKIAA0653 | | B and T cell activation |
| inducible T cell costim | ICOSLG | HGNC:17087 | B7-H2;B7H2;B7RP-1;B7RP1;B7h;CD275;GL50;ICOS-L;ICOSL;LICOS | ENSG00000160223 | B and T cell activation |

| | | | | | |
|---|---|---|---|---|---|
| ulator ligand | | | | | |
| CD40 ligand | CD40LG | HGNC:11935 | CD154;CD40-L;Cd40l;HIGM1;IGM;IMD3;Ly-62;Ly62;T-BAM;TRAP;Tnfsf5;Tnlg8b;gp39;CD154;CD40L;HIGM1;IGM;IMD3;T-BAM;TNFSF5;TRAP;gp39;hCD40L | ENSG00000102245 | activated T cells |
| interleukin 17A | IL17A | HGNC:5981 | Ctla-8;Ctla8;IL-17;IL-17A;Il17;CTLA-8;CTLA8;IL-17;IL-17A;IL17 | ENSG00000112115 | activated T cells |
| CD86 molecule | CD86 | HGNC:1705 | B7;B7-2;B7.2;B70;CLS1;Cd28l2;ETC-1;Ly-58;Ly58;MB7;MB7-2;TS/A-2;B7-2;B7.2;B70;CD28LG2;LAB72 | ENSG00000114013 | T cell activation |
| C-X-C motif chemokine receptor 5 | CXCR5 | HGNC:1060 | Blr1;CXC-R5;CXCR-5;Gpcr6;MDR15;BLR1;CD185;MDR15 | ENSG00000160683 | follicular helper T cells |
| eomesodermin | EOMES | HGNC:3372 | C77258;TBR-2;Tbr2;TBR2 | ENSG00000163508 | T cell exhaustion |
| CD83 molecule | CD83 | HGNC:1703 | BL11;HB15 | ENSG00000112149 | antigen presenting and immune stimulation |
| triggering receptor expressed on myeloid cells 2 | TREM2 | HGNC:17761 | TREM-2;Trem2a;Trem2b;Trem2c;PLOSL2;TREM-2;Trem2a;Trem2b;Trem2c | ENSG00000095970 | anti-inflammation |
| complement C3 | C3 | HGNC:1318 | AI255234;ASP;HSE-MSF;Plp;AHUS5;ARMD9;ASP;C3a;C3b;CPAMD1;HEL-S-62p | ENSG00000125730 | complement system |
| C-X-C motif chemokine receptor 6 | CXCR6 | HGNC:16647 | BB217514;BONZO;STRL33;BONZO;CD186;STRL33;TYMSTR | ENSG00000172215 | induced by inflammatory stimuli, T cell recruiting signal |
| tumor necrosis factor | TNF | HGNC:11892 | DIF;TNF-a;TNF-alpha;TNFSF2;TNFalpha;Tnfa;Tnfsf1a;Tnlg1f;DIF;TNF-alpha;TNFA;TNFSF2;TNLG1F | ENSG00000232810 | pro-inflammation, diverse functions |
| interleukin 22 | IL22 | HGNC:14900 | IL-22;IL-22a;ILTIFa;Iltif;IL-21;IL-22;IL-D110;IL-TIF;ILTIF;TIFIL-23;TIFa;zcyto18 | ENSG00000127318 | chronic inflammation, effect on stromal and |

| | | | | | |
|---|---|---|---|---|---|
| | | | | | epithelial cells |
| Fc fragment of IgG receptor IIa | FCGR2A | HGNC:3616 | CD32;CD32A;CDw32;FCG2;FCGR2;FCGR2A1;FcGR;IGFR2 | ENSG00000143226 | low affinity inhibitory receptor for immunoglobulin gamma Fc region |
| Fc fragment of IgG receptor IIb | FCGR2B | HGNC:3618 | AI528646;CD32;F630109E10Rik;Fc[g]RII;FcgRII;Fcgr2;Fcgr2a;Fcr-2;Fcr-3;Ly-17;Ly-m20;LyM-1;Lym-1;fcRII;CD32;CD32B;FCG2;FCGR2;FCGR2C;FcRII-c;IGFR2 | ENSG00000072694 | low affinity inhibitory receptor for immunoglobulin gamma Fc region |
| C-X-C motif chemokine ligand 5 | CXCL5 | HGNC:10642 | AMCF-II;Cxcl6;ENA-78;GCP-2;LIX;Scyb5;Scyb6;ENA-78;SCYB5 | ENSG00000163735 | chemotaxis of neutrophils |
| C-X3-C motif chemokine receptor 1 | CX3CR1 | HGNC:2558 | CCRL1;CMKBRL1;CMKDR1;GPR13;GPRV28;V28 | ENSG00000168329 | immune cell recruitment |
| chemokine (C-X-C motif) ligand 15 | | | Il8;Scyb15;lungkine;weche | | immune cell recruitment |
| C-X-C motif chemokine ligand 8 | CXCL8 | HGNC:6025 | GCP-1;GCP1;IL8;LECT;LUCT;LYNAP;MDNCF;MONAP;NAF;NAP-1;NAP1 | ENSG00000169429 | immune cell recruitment |
| C-C motif chemokine receptor 2 | CCR2 | HGNC:1603 | Cc-ckr-2;Ccr2a;Ccr2b;Ckr2;Ckr2a;Ckr2b;Cmkbr2;mJe-r;CC-CKR-2;CCR-2;CCR2A;CCR2B;CD192;CKR2;CKR2A;CKR2B;CMKBR2;MCP-1-R | ENSG00000121807 | migration of monocytes |
